# An imbalance between proliferation and differentiation underlies the development of microRNA-defective pineoblastoma

**DOI:** 10.1101/2024.04.23.590638

**Authors:** Claudette R. Fraire, Kavita Desai, Uma A. Obalapuram, Lindsay K. Mendyka, Veena Rajaram, Teja Sebastian, Yemin Wang, Kenan Onel, Jeon Lee, Stephen X. Skapek, Kenneth S. Chen

## Abstract

Mutations in the microRNA processing genes *DICER1* and *DROSHA* drive several cancers that resemble embryonic progenitors. To understand how microRNAs regulate tumorigenesis, we ablated *Drosha* or *Dicer1* in the developing pineal gland to emulate the pathogenesis of pineoblastoma, a brain tumor that resembles undifferentiated precursors of the pineal gland. Accordingly, these mice develop pineal tumors marked by loss of microRNAs, including the let-7/miR-98-5p family, and de-repression of microRNA target genes. Pineal tumors driven by loss of *Drosha* or *Dicer1* mimic tumors driven by *Rb1* loss, as they exhibit upregulation of S-phase genes and homeobox transcription factors that regulate pineal development. Blocking proliferation of these tumors facilitates expression of pinealocyte maturation markers, with a concomitant reduction in embryonic markers. Select embryonic markers remain elevated, however, as the microRNAs that normally repress these target genes remain absent. One such microRNA target gene is the oncofetal transcription factor *Plagl2*, which regulates expression of pro-growth genes, and inhibiting their signaling impairs tumor growth. Thus, we demonstrate that tumors driven by loss of microRNA processing may be therapeutically targeted by inhibiting downstream drivers of proliferation.

## INTRODUCTION

Mutations in microRNA processing genes, such as *DROSHA*, *DICER1*, and *DGCR8*, drive ∼2% of childhood cancers [1, 2]. These mutations arise in a family of cancers that resemble embryonic progenitors, including Wilms tumor, pleuropulmonary blastoma, rhabdomyosarcoma, and the brain tumor pineoblastoma [3–11]. It is thought that loss of microRNA production leads to de-repression of the genes that are normally repressed by these microRNAs, leading to tumor formation [12, 13]. However, microRNA loss leads to transcriptome-wide changes, with different effects in each cell type, and it is challenging to separate genes upregulated through loss of direct microRNA targeting from secondary effects. In addition, little is known about the specific microRNAs or microRNA target genes that regulate the balance between proliferation and differentiation in each cancer type. As a result, there is currently no way to therapeutically target these mutations.

Pineoblastoma is a rare brain tumor that resembles embryonic neurogenic progenitors of the pineal gland, and it is treated with chemotherapy, surgery, and radiation [14]. However, the morbidities associated with each of these treatment modalities limits their utility; cranial radiation therapy, for instance, is generally avoided in young children. Most other *DICER1/DROSHA*-related tumors, which exhibit point mutations in these genes that incompletely impair microRNA production, but pineoblastoma can uniquely exhibit complete loss of *DROSHA* or *DICER1*, leading to complete loss of microRNA production [11, 15]. Specifically, recent molecular studies have identified three mutational patterns in pineoblastoma: (1) bi-allelic loss of *DROSHA*, *DICER1*, or *DGCR8*; (2) bi-allelic loss of *RB1*; or (3) activation of *MYC* or *FOXR2* [16–19]. The RB1 tumor suppressor normally regulates E2F transcription factors to allow progression through the G1/S cell cycle checkpoint [20], and MYC orchestrates tumorigenesis by binding “E-box” motifs in promoters or enhancers of tumor-promoting genes [21]. However, the microRNAs and microRNA target genes that mediate tumorigenesis in tumors driven by loss of *DROSHA* or *DICER1* remain poorly understood.

Previous studies have used *Rb1* loss, cyclin D1 upregulation, and *Dicer1* loss to model the development of pineoblastoma in mice [22–27]. To dissect how microRNA loss drives formation of pineoblastoma, we generated mice with loss of microRNAs in the developing pineal gland. Here we show that loss of *Drosha* or *Dicer1* mimics loss of the *Rb1* tumor suppressor by driving cell cycle progression through de-repression of cyclin D2 (*Ccnd2*). Uncontrolled proliferation itself restrains tumor cells in an undifferentiated state, and restraining proliferation leads them to upregulate differentiation markers. In addition to being directly targeted by microRNAs, we find that *Ccnd2* is also indirectly regulated by microRNAs through the transcription factor *Plagl2,* and blocking other Plagl2 target genes also slows tumor growth. Lastly, we show that human pineoblastoma tumors exhibit parallel expression patterns.

Together, our results demonstrate the potential for therapeutic targeting of the downstream effects of microRNA loss in *DROSHA-* or *DICER1-*mutant cancers.

## RESULTS

### Pineal tumors driven by loss of Drosha, Dicer1, or Rb1 resemble embryonic pineal progenitors

Prior studies had demonstrated that mice develop pineoblastoma upon pineal-directed ablation of *Rb1* and *p53* (*IrbpCre*;*p53*^flox/flox^;*Rb1*^flox/flox^, hereafter abbreviated as IPRb1) [27]. These studies used a Cre recombinase line driven by the promoter of interphotoreceptor retinoid binding protein (*Irbp*, also known as *Rbp3*), which is expressed in retinal and pineal progenitors [28, 29]. To understand how microRNA loss regulates pineal tumor formation, we developed mice with analogous loss of *Drosha* and *p53* (*IrbpCre*;*p53*^flox/flox^;*Drosha*^flox/flox^, abbreviated as IPDrosha) (**Fig. 1A,B**). These mice were healthy at birth, but at 5-10 months of age, they began to exhibit signs of brain tumor formation. Within days of developing an enlarged skull, they become moribund. Similarly, pineal tumors also arise upon conditionally deleting *Dicer1* and *p53* (*IrbpCre*;*p53*^flox/flox^;*Dicer1*^flox/flox^, or IPDicer1) (**Fig. 1A,B**). By microscopic evaluation, IPDrosha, IPDicer1, and IPRb1 tumors contain large, pleomorphic, primitive cells with a high nucleus-to-cytoplasm ratio and rare inconspicuous nucleoli that resemble embryonic pineal progenitors. These cells are arranged as sheets, in a vaguely nodular pattern, with nuclear molding, frequent mitosis, and areas of necrosis. These cells stain positive for markers of proliferation and neuronal lineage (Ki-67 and synaptophysin, respectively) and negative for a glial lineage marker (glial fibrillary acidic protein, or GFAP) (**Fig. 1C,D**). Furthermore, just as *DROSHA/DICER1*-related pineoblastomas arise later in childhood than *RB1*-driven pineoblastoma, IPDrosha and IPDicer1 tumors appeared to arise at later ages than IPRb1 tumors (median age of onset 269 days, 239 days, and 182 days, respectively), though we observed tumors as early as ∼5 months of age in all three groups. We did not observe any histological abnormalities in the retina, where *IrbpCre* is also active (not shown).

**Figure 1.**
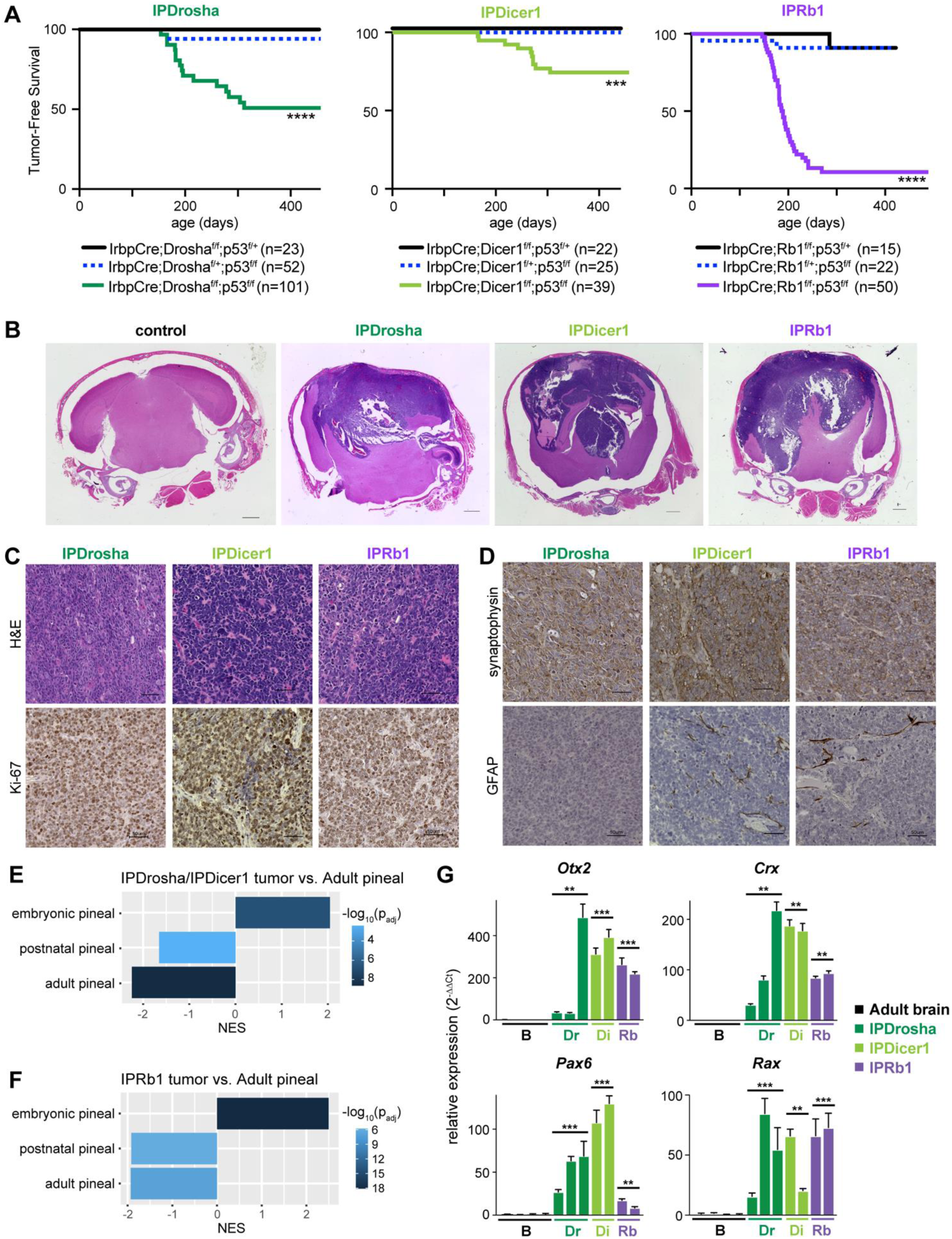
Pineal tumors driven by loss of *Drosha*, *Dicer1*, or *Rb1* resemble embryonic pineal progenitors. **(A)** Kaplan-Meier plots showing tumor-free survival of IPDrosha, IPDicer1, and IPRb1 tumors. IrbpCre;Drosha^f/f^;p53^f/f^ (IPDrosha, dark green) mice have a tumor incidence of 36% at 1 year; IrbpCre;Dicer1^f/f^;p53^f/f^ (IPDicer1, light green) mice have a tumor incidence of 34% at 1 year; IrbpCre;Rb1^f/f^;p53^f/f^ (IPRb1, purple) mice have a tumor incidence of 90% at 1 year (***p<0.001 and ****p<0.0001; vs. control genotypes; by log-rank test). **(B)** Photomicrographs of brains from control (Drosha^f/f^;p53^f/f^), IPDrosha, IPDicer1, and IPRb1 mice show expansile pineal tumors expanding into the skull above and normal brain below (scale bar 1000 µm). **(C)** Photomicrographs of IPDrosha, IPDicer1, and IPRb1 tumors (scale bar 50 µm). Top row shows H&E, and bottom row shows immunostaining for Ki-67. **(D)** Photomicrographs of IPDrosha, IPDicer1, and IPRb1 tumors (scale bar 50 µm), with immunostaining for synaptophysin (top) and GFAP (bottom). **(E)** Gene set enrichment analysis (GSEA) of enrichment for pineal development markers comparing IPDrosha/IPDicer1 tumors with normal adult pineal gland. **(F)** GSEA of enrichment for pineal development markers comparing IPRb1 tumor with normal adult pineal gland. **(G)** qPCR for pineal homeobox genes (*Otx2, Crx, Pax6*, and *Rax*) in brain, IPDrosha, IPDicer1, and IPRb1 tumors. Values shown are mean ± SD (**p < 0.01 and **p < 0.001; vs. normal brain; two-sided t-test). See also Supplementary Figure S1.

To understand how these tumors arise, we next profiled gene expression and histone modification patterns, focusing on acetylation of histone H3 lysine 27 (H3K27ac), which marks active promoters and enhancers, and trimethylation of histone H3 lysine 4 (H3K4me3), which marks active promoters. We found that IPDrosha and IPRb1 tumors retain the gene expression patterns of embryonic pineal progenitors, with enrichment for embryonic pineal markers and depletion for adult markers (**Fig. 1E-F**, **Suppl. Fig. S1A-D**). Specifically, the pineal homeobox transcription factors *Otx2, Crx, Pax6*, and *Rax* are expressed at high levels in IPDrosha, IPDicer1, and IPRb1 tumors, reflecting their shared pineal progenitor identity (**Fig. 1G**) [30]. Accordingly, the genomic regions surrounding *Otx2, Crx, Rax*, and *Pax6* are marked by broad H3K27ac signal, sometimes termed “super-enhancers” [31] (**Suppl. Fig. S1E**). Broad H3K27ac signal at these homeobox transcription factors indicates the importance of these genes to maintain embryonic identity in these pineal tumors.

Next, we examined whether an embryonic pineal expression signature was unique to these tumors, or whether other murine cancer models might also exhibit a similar signature. To do this, we identified publicly available RNA-seq datasets of mouse models of cancer with paired non-cancer control tissue. Specifically, we examined data from embryonic neuronal brain tumors (*SHH*-driven medulloblastoma and Group 3 [*MYC/SMARCA4*-driven] medulloblastoma), glial brain tumors (*MYCN*-driven high-grade glioma, glioblastoma allografts, and histone H3.3 K27M diffuse midline glioma), and embryonic neuroectodermal extracranial tumors (TH-*MYCN* neuroblastoma). Combined with our comparisons of IPDrosha/IPDicer1 tumors and IPRb1 tumors with normal adult pineal glands, this constituted a total of eight comparisons between a murine cancer model and paired non-cancer control tissue.

We used four proliferation-related Molecular Signatures Database (MSigDB) hallmark gene sets to define a proliferation signature [32]. For all eight comparisons, this proliferation signature was significantly enriched in tumors compared to normal tissues (**Suppl. Fig. S2A**). We then removed proliferation markers from our developmental pineal marker sets to identify proliferation-independent embryonic and adult pineal markers. In this analysis, IPDrosha/IPDicer1 and IPRb1 tumors were the only tumors to show a statistically significant enrichment for proliferation-independent embryonic pineal markers (**Suppl. Fig. S2B**). Similarly, IPDrosha/IPDicer1 and IPRb1 tumors showed the most significant depletion of proliferation-independent adult pineal markers (**Suppl. Fig. S2C**). Together, these results indicate that IPDrosha, IPDicer1, and IPRb1 tumors all arise from a similar pineal gland progenitor. Furthermore, IPDrosha/IPDicer1 tumors resemble embryonic pineal precursors lacking microRNAs, and IPRb1 tumors resemble embryonic pineal precursors lacking cell cycle controls.

### Drosha/Dicer1-driven tumors lack microRNAs and overexpress microRNA target genes

To understand how IPRb1 tumors form with intact microRNA processing and, conversely, how IPDrosha tumors form with intact Rb1 function, we next investigated differences between IPRb1 and IPDrosha tumors. We first performed small RNA sequencing in IPDrosha tumors, IPRb1 tumors, matched normal brain, and age-matched normal adult pineal glands to identify the microRNAs regulating normal pineal gland development. As expected, IPDrosha tumors are devoid of canonical microRNAs (**Fig. 2A,B**, **Suppl. Fig. S3A**, **Suppl. Data 1**). Some microRNAs can be produced without *Drosha*, through a non-canonical pathway, if they are processed by the splicing machinery (called “miRtrons”) or if they are transcribed as a capped microRNA [33, 34]. These “Drosha-independent” microRNAs remain expressed in IPDrosha tumors at levels similar to matched normal brain, age-matched adult pineal glands, or IPRb1 tumors (**Suppl. Fig. S3B-D**). As expected, IPDicer1 tumors similarly expressed undetectable levels of microRNAs (**Fig. 2C**). Together, these findings are consistent with *Drosha* loss.

**Figure 2.**
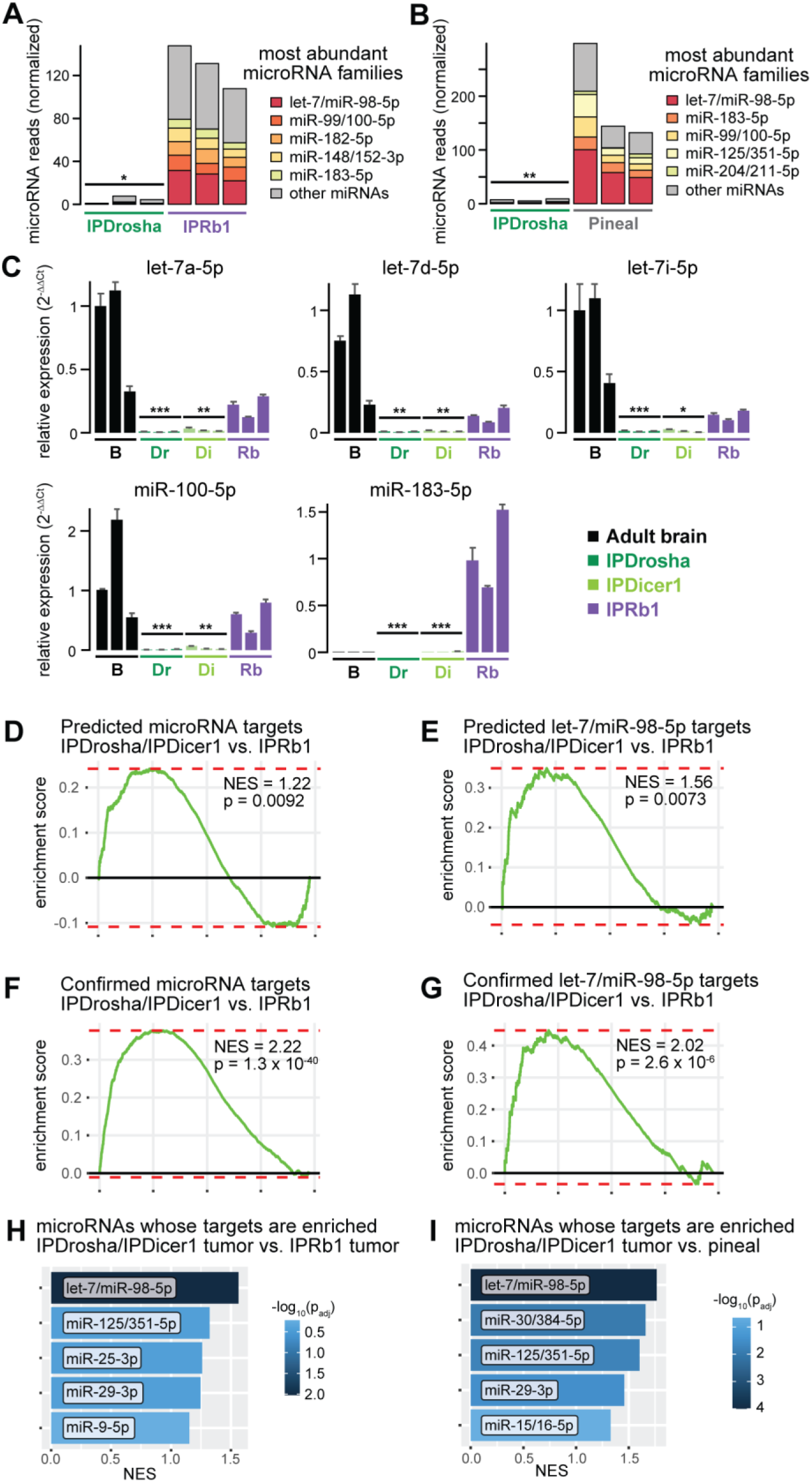
*Drosha/Dicer1*-driven tumors lack microRNAs and overexpress microRNA target genes. **(A)** Small RNA sequencing displaying the most abundant microRNA families in IPRb1 tumors and IPDrosha tumors. **(B)** Small RNA sequencing displaying the most abundant microRNA families in adult pineal glands and IPDrosha tumors. **(C)** TaqMan miRNA expression of let-7a-5p, let-7d-5p, let-7i-5p, miR-100-5p, and miR-183-5p in normal adult brain, IPDrosha, IPDicer1, and IPRb1 tumors. **(D,E)** GSEA of enrichment for predicted targets of all microRNAs expressed in IPRb1 tumors **(D)** or predicted let-7/miR-98-5p targets **(E)** in IPDrosha/IPDicer1 vs. IPRb1 tumors. **(F,G)** GSEA for enrichment of confirmed top targets of microRNAs **(F)** or let-7/miR-98-5p **(G)** in IPDrosha/IPDicer1 vs. IPRb1 tumors. **(H-I)** GSEA showing enrichment for target genes of individual microRNA families in total RNA sequencing of IPDrosha/IPDicer1 vs. IPRb1 tumors **(H)** or adult pineal gland **(I)**. See also **Supplementary Figures S3** and **S4**.

We identified the most abundant microRNAs in age-matched adult pineal glands and in IPRb1 tumors, which resemble embryonic pineal progenitors. Members of the let-7/miR-98-5p, miR-99/100-5p, and miR-183-5p families are among the most abundant microRNA families in both IPRb1 tumors and adult pineal glands (**Fig. 2A,B, Suppl. Fig. S3E-F**). The let-7/miR-98-5p and miR-99/100-5p families are co-transcribed from the same genomic loci in an evolutionarily conserved manner [35]. They are not expressed in neural progenitors but are induced late in neural or retinal development to downregulate progenitor genes [36–38]. In IPRb1 tumors and adult pineal glands, we detected nearly all let-7/miR-98-5p family members (**Suppl. Fig. S3G-H**). The depletion of mature let-7/miR-98-5p microRNAs was seen in IPDrosha tumors despite broad H3K27ac and H3K4me3 at the host genes of let-7/miR-98-5p transcripts (**Suppl. Fig. S3I**). Similarly, miR-183-5p is one of the most abundant microRNAs in the adult retina and pineal gland, where it regulates genes involved in the circadian rhythm [39–41]. Germline mutations in the region encoding this microRNA produce retinal degeneration in humans [40].

To understand which microRNAs have the greatest effect on gene expression in IPDrosha tumors, we next examined the expression of genes predicted to be targeted by the 50 microRNAs expressed in IPRb1 tumors but not in IPDrosha tumors. Based on TargetScan calculations of sequence context and conservation, we found 781 genes predicted to be targeted by these 50 microRNAs [42]. These predicted microRNA targets are expressed at higher levels in IPDrosha tumors than IPRb1 tumors, consistent with a loss of microRNA repression (**Fig. 2D**).

However, observing loss of a microRNA and upregulation of a predicted target transcript cannot fully distinguish between the direct and indirect effects of microRNA targeting. Some genes may be upregulated as a secondary effect of another de-repressed microRNA target gene. Thus, we next used microRNA enhanced cross-linking immunoprecipitation (miR-eCLIP) sequencing to confirm which genes are directly targeted by microRNAs in IPRb1 tumors [43]. This technique directly identifies microRNA-mRNA interactions through immunoprecipitation of Argonaute 2 (AGO2) and ligation of microRNA-mRNA “chimeras”. We performed miR-eCLIP sequencing in two independent IPRb1 tumors, and we examined the non-chimeric peaks and chimeric peaks that were reproduced between the two samples (**Suppl. Fig. S4A**, **Suppl. Data 2**). As expected, reproduced reads clustered primarily in the 3’ untranslated regions (3’UTRs) of coding genes, confirming enrichment for regions captured by microRNA targeting [12] (**Suppl. Fig. S4B**). In non-chimeric transcripts immunoprecipitated with AGO2, two of the three most significantly enriched motifs were let-7/miR-98-5p recognition sequences (**Suppl. Fig. S4C**). Furthermore, let-7/miR-98-5p family members were commonly identified in the microRNA side of chimeric reads (**Suppl. Fig. S4D**). Altogether, the results from our integrated sequencing analyses highlight let-7/miR-98-5p microRNAs as key drivers of the gene expression differences between IPDrosha/IPDicer1 tumors and IPRb1 tumors.

Next, we examined whether the genes identified as direct microRNA targets in miR-eCLIP sequencing of IPRb1 tumors are repressed compared to IPDrosha/IPDicer1 tumors. Genes represented in microRNA-mRNA chimeric reads were significantly upregulated in IPDrosha/IPDicer1 tumors, and they were more likely to be upregulated than downregulated (**Suppl. Fig. S4E,F**). These confirmed microRNA target genes were even more upregulated in IPDrosha/IPDicer1 tumors than the microRNA targets predicted by TargetScan (**Fig. 2D,F**). Similarly, the confirmed top targets of the let-7/miR-98-5p family were more enriched than the set of predicted let-7/miR-98-5p targets (**Fig. 2E,G**). In other words, much of the transcriptomic signature in IPDrosha/IPDicer1 tumors can be directly attributed to loss of let-7/miR-98-5p.

Together, these results demonstrate that de-repression of direct microRNA targets explain many of the differences between IPDrosha and IPRb1 tumors. Nevertheless, genes represented in microRNA-mRNA chimeras only constituted 23.1% of all genes overexpressed in IPDrosha tumors (**Suppl. Fig. S4F**). In other words, while microRNAs directly drive many of the transcriptomic differences between IPDrosha and IPRb1 tumors, up to 77% of the genes overexpressed in IPDrosha tumors may be attributable to indirect effects.

### Loss of microRNAs and loss of Rb1 converge on overlapping tumor-driving effects

To understand how de-repression of microRNA target genes causes cancer, we next examined other gene sets upregulated by loss of microRNAs, whether directly or indirectly. Specifically, we identified “hallmark” gene sets differentially enriched in IPDrosha/IPDicer1 tumors, IPRb1 tumors, and normal adult pineal glands. We noted that four of the top five most enriched hallmark gene sets overlapped between IPDrosha/IPDicer1 and IPRb1 tumors, demonstrating their overlapping paths to tumor formation (**Fig. 3A, Suppl. Data 3**). These similarities in overall transcriptome, despite independent molecular mechanisms, suggests that *Drosha/Dicer1* loss and *Rb1* loss converge to drive cancer formation through overlapping downstream effects.

**Figure 3.**
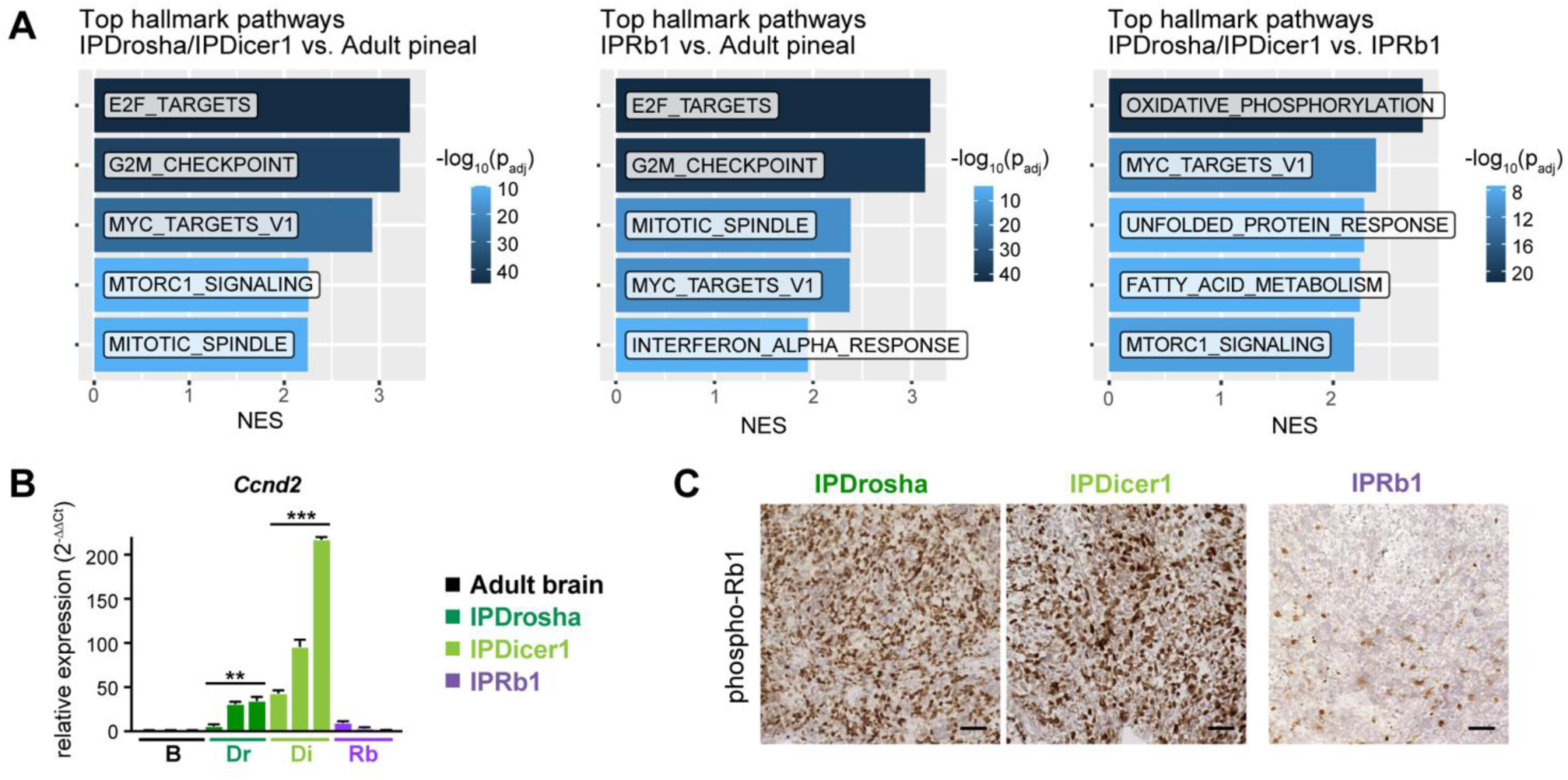
*Drosha/Dicer1*-driven tumors are highly proliferative. **(A)** Hallmark gene sets most enriched in IPDrosha/IPDicer1 tumors vs. age-matched adult pineal gland, IPRb1 tumors vs. age-matched adult pineal gland, and IPDrosha/IPDicer1 tumors vs. IPRb1 tumors. **(B)** qPCR for *Ccnd2* in normal adult brain, IPDrosha, IPDicer1, and IPRb1 tumors. Values shown are mean ± SD (*p< 0.05; vs. normal brain; two-sided t-test). **(C)** Photomicrographs of IPDrosha, IPDicer1, and IPRb1 tumors stained for phospho-Rb1 (Ser807/811) (scale bar 50 µm).

As expected, since Rb1 normally regulates E2F transcription factors, the most enriched “hallmark” gene set in IPRb1 tumors is “E2F target genes” (**Fig. 3A**, **Suppl. Fig. S5B**). However, E2F target genes were also the most enriched gene set in IPDrosha tumors (**Fig. 3A**, **Suppl. Fig. S5A**). In cells with intact Rb1, D-type cyclins (cyclin D1, D2, or D3) partner with cyclin-dependent kinases 4 and 6 (CDK4/6) to phosphorylate Rb1 and release it from inhibiting E2Fs [20, 44]. It had previously been shown that cyclin D1 (*Ccnd1*) overexpression was sufficient to replace Rb1 loss and drive pineal tumor formation [23, 45]. Thus, we examined the expression of D-type cyclins in IPDrosha and IPDicer1 tumors and found that cyclin D2 (*Ccnd2*) was the only one significantly overexpressed (**Fig. 3B**, **Suppl. Fig. S5C**). *Ccnd2* is a known target of several microRNAs; for example, it is thought to cause proliferation of leukemias driven by loss of miR-16 [46, 47]. We confirmed that *Ccnd2* is regulated by several microRNAs in our miR-eCLIP sequencing, including the miR-15/16-5p family (**Suppl. Fig. S5D**). Using immunohistochemistry, we also confirmed that IPDrosha and IPDicer1 tumors exhibit high levels of CDK4/6 activity, as nearly every cell stained positive for phosphorylated Rb1 (**Fig. 3C**). (As expected, the tumor cells in IPRb1 tumors lacked Rb1, though some entrapped non-tumor cells stained positive.) Thus, *Drosha* loss and *Rb1* loss converge on the same proliferation-driving pathway to drive cancer cells through the G1/S cell cycle checkpoint.

Another hallmark gene set upregulated in both IPDrosha/IPDicer1 and IPRb1 tumors compared to normal adult pineal was “Myc targets V1” (**Fig. 3A**). This gene set was also one of the most enriched genesets in IPDrosha/IPDicer1 compared to IPRb1 tumors (**Fig. 3A**). *MYC* drives many cancers by activating transcription of proliferation drivers [21, 48, 49], and gain of *MYC* or *FOXR2* is known to drive a third subset of human pineoblastoma [19]. In our RNA-seq, *Mycn* is the most abundant *Myc* family member in IPDrosha, IPDicer1, and IPRb1 tumors, but it was not significantly overexpressed in IPDrosha/IPDicer1 compared to IPRb1 tumors (**Suppl Fig. S5E**). Nevertheless, *Mycn* was one of the most abundant mRNA transcripts in miR-eCLIP chimeras, and it was targeted by a large number of microRNAs (**Suppl. Fig. S5F**).

### Proliferation restrains differentiation in Drosha/Dicer1-driven tumors

In the developing brain, proliferation maintains progenitors in an undifferentiated state, and differentiation is allowed to occur once proliferation slows down [50]. Accordingly, IPRb1 tumors remain undifferentiated solely because of unrestrained proliferation of these embryonic progenitors. Thus, to understand whether IPDrosha/IPDicer1 tumors with high levels of *Ccnd2* are dependent on CDK4/6 activity to remain undifferentiated, we treated them with the clinically approved CDK4/6 inhibitor palbociclib [51]. Specifically, we implanted tumor cells as subcutaneous allografts and administered either vehicle or palbociclib to recipient mice. Over two weeks, palbociclib suppressed proliferation and extended survival in IPDrosha and IPDicer1 tumors (**Fig. 4A** and **Suppl. Fig. S6A**). Palbociclib-treated tumors exhibited less staining for phosphorylated Rb1 and Ki-67, consistent with a loss of CDK4/6 activity (**Fig. 4B and Suppl. Fig. S6C**). Histologically, the palbociclib-treated IPDrosha and IPDicer1 tumors exhibit increased karyorrhexis, a sign of cell death, and increased background neuropil, a sign of neuronal differentiation (**Fig. 4B**).

**Figure 4.**
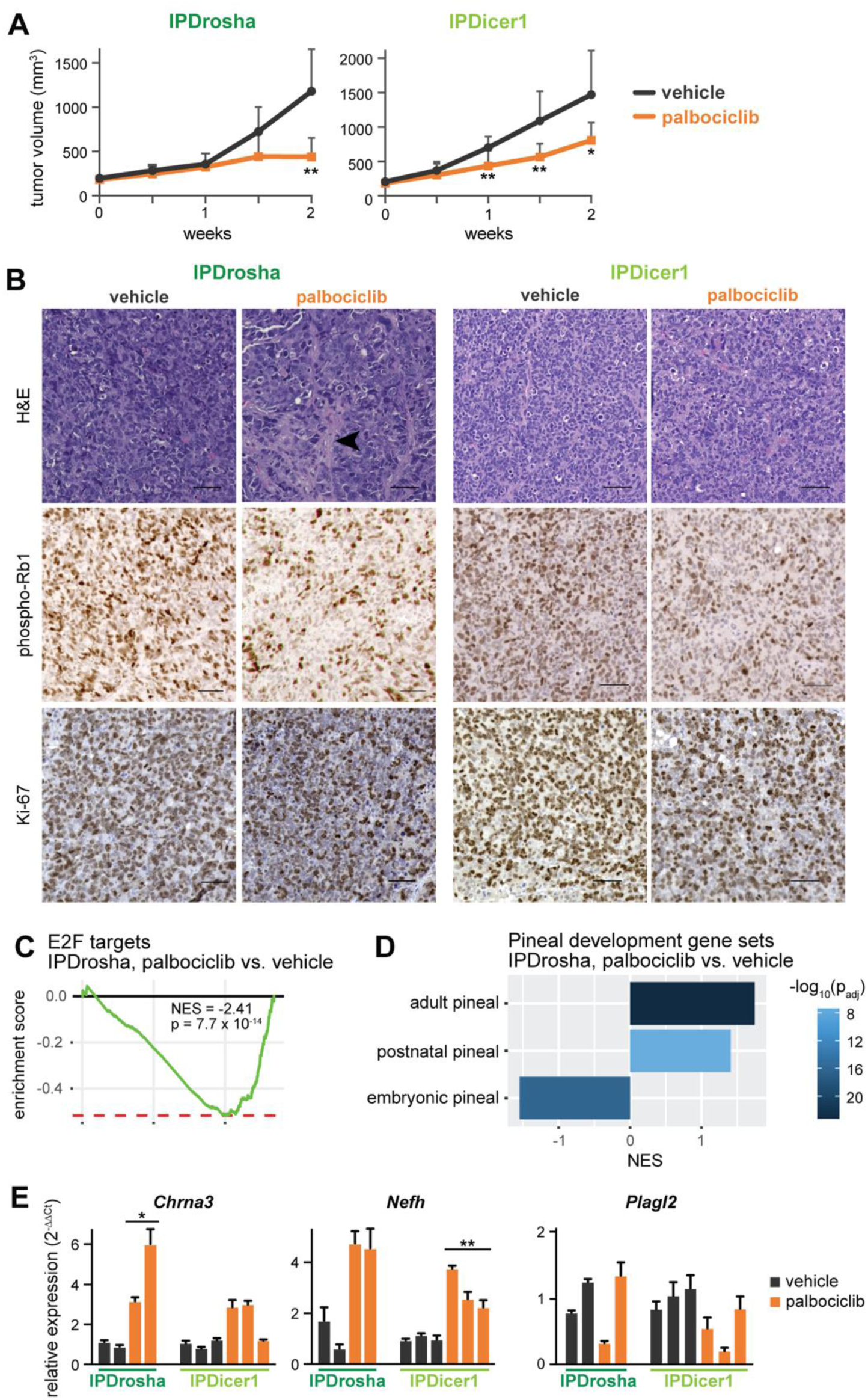
Palbociclib restrains proliferation and induces differentiation. **(A)** Tumor volumes for IPDrosha or IPDicer1 tumors in mice dosed with vehicle or palbociclib (n=6 or 8 for IPDrosha, respectively; n=9 or 7 for IPDicer1, respectively). Palbociclib was given at 150 mg/kg/day by oral gavage, 5 days/week. Values shown are mean ± SD (*p<0.05 and *p<0.01; vs. vehicle-treated; two-sided t-test). **(B)** Photomicrographs of vehicle-or palbociclib-treated IPDrosha and IPDicer1 tumors (scale bar 50 µm). Top row shows H&E, middle row shows immunostaining for phospho-Rb1 (Ser807/811); and bottom row shows staining for Ki-67. Arrowhead denotes area of neuropil accumulation. **(C)** GSEA for E2F target genes in RNA sequencing of IPDrosha tumor allografts treated with palbociclib vs. vehicle. **(D)** GSEA of enrichment for pineal development genes in IPDrosha tumors treated with palbociclib vs. vehicle. **(E)** qPCR for adult pineal marker genes (*Chrna3* and *Nefh*) and an embryonic pineal marker gene (*Plagl2*) in IPDrosha and IPDicer1 tumors treated with palbociclib vs. vehicle. Values shown are mean ± SD (*p<0.05; vs. vehicle-treated; two-sided t-test). See also **Supplementary Figure S6**.

Next, we used RNA-seq to understand how palbociclib affected IPDrosha tumor cells. Consistent with inhibition of CDK4/6, E2F target genes were the most downregulated hallmark gene set in palbociclib-treated tumors (**Fig. 4C**). In contrast, interferon alpha and gamma response pathways were enriched in palbociclib-treated tumors, consistent with reports that CDK4/6 inhibitors can induce antitumor immunity [52–55] (**Suppl. Fig. S6B**). When we examined pineal differentiation markers, we found that palbociclib-treated tumors were enriched for adult pineal markers such as *Chrna3* and *Nefh* (**Fig. 4D,E, Suppl. Fig. S6D**). On the other hand, embryonic markers were mildly depleted as a whole (**Fig. 4D**, **Suppl. Fig. S6E**). Many embryonic genes remain highly expressed upon palbociclib treatment, such as the embryonic transcription factor *Plagl2* (**Fig. 4E**). In other words, blocking the effect of *Ccnd2* partially restores differentiation in IPDrosha tumors, but microRNAs remain absent, allowing some embryonic pineal markers to remain highly expressed.

### The microRNA target gene Plagl2 regulates proliferation drivers Igf2 and Ccnd2

Because CDK4/6 inhibition could induce expression of adult genes but could not suppress the embryonic gene signature, we next identified the proliferation-independent microRNA target genes that contribute to maintaining an embryonic expression program. Specifically, we identified embryonic pineal markers that were overexpressed in IPDrosha tumors, confirmed “top” let-7/miR-98-5p target genes by miR-eCLIP, and independent of palbociclib. Three genes met these criteria: *Bach1*, *Tgfbr1*, and *Plagl2*. We focused on *Plagl2* (*PLAG1*-like gene 2) because of previous evidence that it is an oncogenic, developmental transcription factor, and because its most well-characterized target gene, *Igf2*, was the single most overexpressed gene in IPDrosha tumors [56–58] (**Fig. 5A-B**). Our miR-eCLIP sequencing demonstrated that the 3’ UTR of *Plagl2* interacted with several microRNAs (**Fig. 5C**). In fact, let-7/miR-98-5p microRNAs interacted with *Plagl2* at three sites, including one highly conserved site with a 7-mer perfect match (**Fig. 5C**).

**Figure 5.**
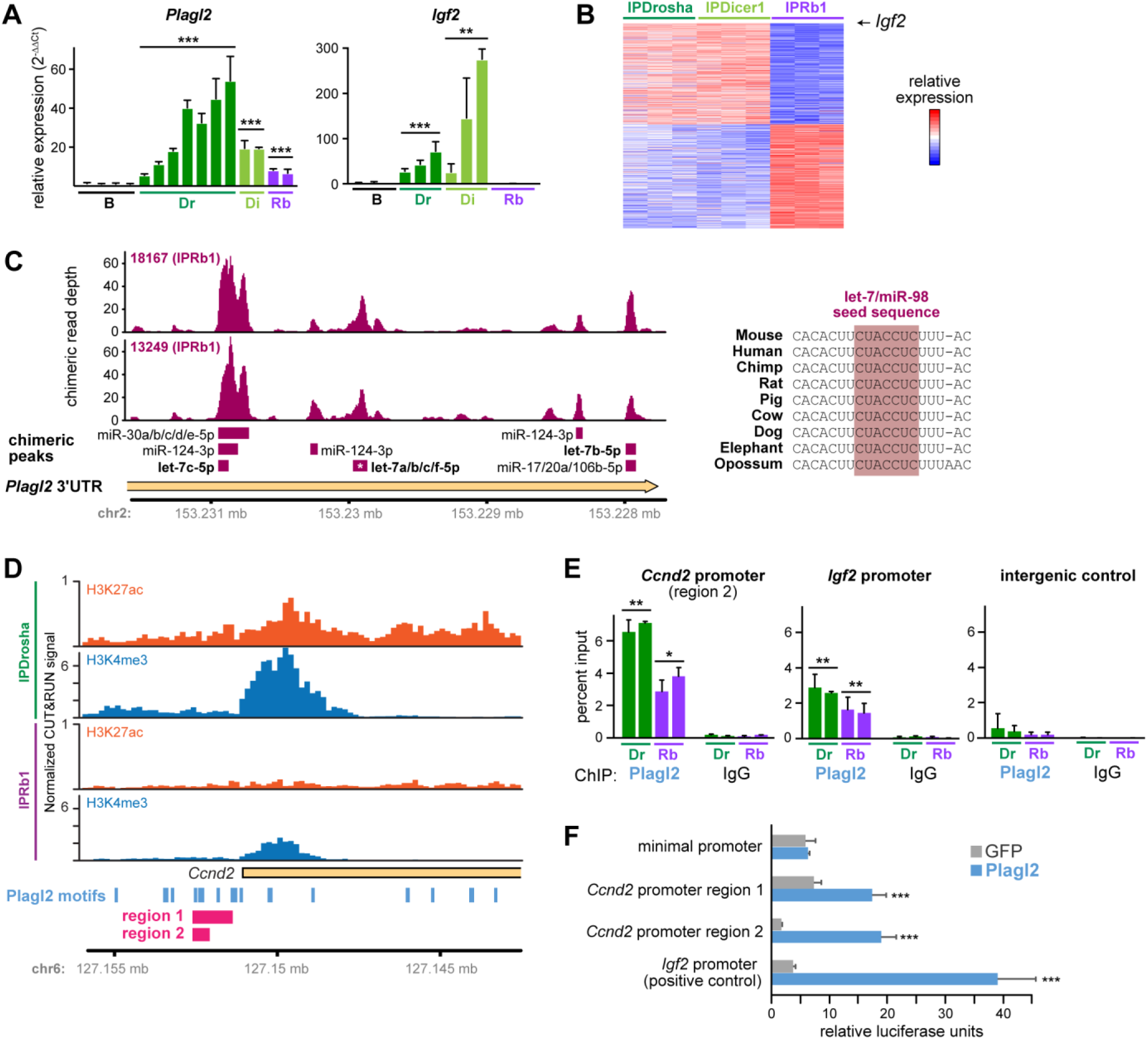
The microRNA target gene *Plagl2* regulates proliferation drivers *Igf2* and *Ccnd2*. **(A)** qPCR for *Igf2* and *Plagl2* in normal adult brain, IPDrosha, IPDicer1, and IPRb1 tumors. Values shown are mean ± SD (*p< 0.05 and ***p<0.001; vs. normal adult brain; two-sided t-tests). **(B)** Heat map from total RNA sequencing of IPDrosha, IPDicer1, and IPRb1 tumors highlighting *Igf2* (n= 3 tumors each). **(C)** miR-eCLIP chimeric reads from IPRb1 tumors of the 3’ UTR of *Plagl2,* with microRNA chimeric read peaks denoted below, including three peaks containing let-7/miR-98-5p family members. The conservation of the second let-7/miR-98-5p binding site (marked with an asterisk) is featured on the right. **(D)** H3K27ac and H3K4me3 signal at the *Ccnd2* promoter in IPDrosha and IPRb1 tumors, with predicted Plagl2 binding sites and regions used for luciferase reporter assay. **(E)** ChIP-qPCR detecting Plagl2 occupancy in IPDrosha and IPRb1 tumors at the mouse *Ccnd2* promoter, *Igf2* promoter, and a negative-control intergenic region. Values shown are mean ± SD (*p<0.05 and **p<0.01; vs. IgG; two-sided t-test). **(F)** Luciferase activity of cells transiently transfected with luciferase reporter (preceded by listed promoter regions) and an expression construct encoding either *Plagl2* or GFP. Values shown are relative ratio of firefly to *Renilla* luciferase from six replicate wells per condition, mean ± SD (***p<0.001; vs. GFP; two-sided t-test). See also **Supplementary Figure S7**.

Histone marks at the *Plagl2* locus were similar in IPDrosha and IPRb1 tumors, consistent with a gene whose expression is primarily regulated by microRNAs post-transcriptionally (**Suppl. Fig. S7A**). At the *Ccnd2* locus, however, IPDrosha tumors had much higher levels of H3K27ac and H3K4me3, suggesting that *Ccnd2* upregulation is due to a combination of transcriptional and post-transcriptional regulation (**Fig. 5D**). Since the *Ccnd2* promoter also contains several predicted Plagl2 binding sites, we examined whether Plagl2 resides at the *Ccnd2* promoter in IPDrosha tumors (**Fig. 5D**). Using ChIP-qPCR, we verified Plagl2 binding at a region of the *Ccnd2* promoter near Plagl2 binding motifs (**Fig. 5E**). Lastly, we noted that levels of *Plagl2*, *Igf2,* and *Ccnd2* are high in embryonic pineal glands and fall in parallel during pineal development, suggesting coordinated regulation (**Suppl. Fig. S7B**).

To verify that Plagl2 can activate this region of the *Ccnd2* promoter, we next cloned two fragments of the *Ccnd2* promoter into a luciferase reporter plasmid (denoted as regions 1 and 2 in **Fig. 5D**). The first region was a 1.2kb fragment that extended nearly to the *Ccnd2* transcriptional start site, and the second region was a 0.5kb fragment centered around the cluster of Plagl2 binding sites we assayed by ChIP-qPCR. We used a minimal promoter as a negative control and a 0.9kb fragment from the *Igf2* promoter as a positive control. We co-transfected these reporters into HEK293 cells with an expression construct encoding either *Plagl2* or GFP, and we found that *Plagl2* activated both Ccnd2 reporters (**Fig. 5F**, **Suppl. Fig. S7C**). Together, these results suggest that *Ccnd2* is upregulated by both increased transcription and decreased microRNA repression, and that the microRNA target gene *Plagl2* contributes to the maintenance of the embryonic expression program, in part by regulating *Igf2* and *Ccnd2*.

### Igf1r inhibition blocks growth of Drosha/Dicer1-driven tumors

*Igf2*, whose overexpression drives many other childhood cancers, signals through the insulin signaling pathway primarily by binding the insulin-like growth factor 1 receptor (Igf1r), leading to activation of mTORC1 and other kinase pathways [59]. Accordingly, we observed that the “insulin signaling” and “mTORC1 signaling” gene sets were significantly enriched in IPDrosha/IPDicer1 tumors (**Fig. 6A** and **Fig. 3A**). Since Igf1r can signal through the Akt/mTOR or mitogen-activated protein kinase (MAPK) pathways, we next used Western blots to examine the relative activities of these signaling pathways in IPDrosha and IPRb1 tumors. These showed somewhat higher Akt/mTOR activity (phosphorylation of Akt and S6) in the IPDrosha tumors but similar levels of MAPK activity (Erk1/2 phosphorylation) (**Suppl. Fig. 8A)**. In both IPDrosha and IPDicer1 tumors, we found that ceritinib significantly impaired tumor growth (**Fig. 6B**). Vehicle– and ceritinib-treated IPDrosha and IPDicer1 tumors are histologically similar, though ceritinib-treated tumors appeared to have slightly more background neuropil (**Fig. 6C**). By immunostains, ceritinib-treated tumors exhibited less phosphorylation of S6, as expected for an Igf1r inhibitor (**Fig. 6C**, **Suppl. Fig. S8B**). Similarly, RNA sequencing revealed depletion of the insulin signaling genes in ceritinib-treated IPDrosha tumors (**Fig. 6D**). Similarly, we also treated IPRb1 tumors with ceritinib (**Suppl. Fig. S8C**). Unlike in IPDrosha/IPDicer1 tumors, treatment with ceritinib did not impair tumor growth, which confirms that the gene expression differences in IPDrosha/IPDicer1 tumors renders them specifically sensitive to ceritinib.

**Figure 6.**
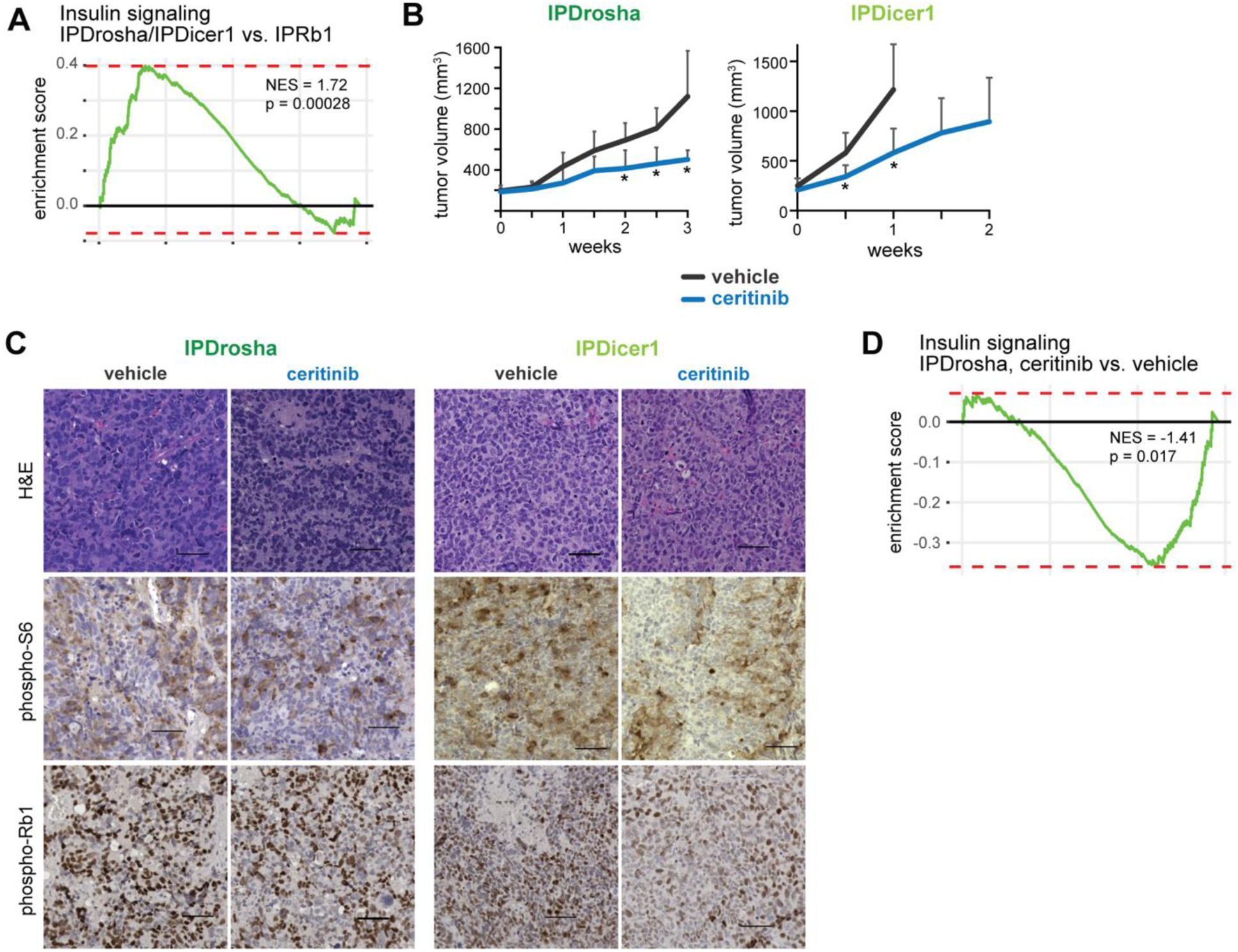
The Igf inhibitor ceritinib suppresses tumor growth. **(A)** GSEA of enrichment for insulin signaling in IPDrosha vs. IPRb1 tumors (Wikipathway WP481). **(B)** Tumor volumes for IPDrosha and IPDicer1 tumors in mice dosed with vehicle or ceritinib (n=5 for IPDrosha, respectively; n=5 or 7 for IPDicer1, respectively). Ceritinib was given at 50 mg/kg/day by oral gavage, 5 days/week. Values shown are mean ± SD (*p< 0.05; vs. vehicle-treated; two-sided t-test). **(C)** Photomicrographs of vehicle-or ceritinib-treated IPDrosha and IPDicer1 tumors (scale 50 µm). Top row shows H&E, the middle row shows phospho-S6 (Ser235/236), and the bottom row shows phospho-Rb1 (Ser807/811). **(D)** GSEA of enrichment for insulin signaling gene set in IPDrosha tumors treated with ceritinib vs. vehicle.

We also observed that ceritinib-treated IPDrosha tumors downregulated Rb1 phosphorylation (**Fig. 6C**, **Suppl. Fig. S8B**). We thus tested whether the combination of ceritinib and palbociclib would be additive in IPDrosha tumors. To achieve tolerability of the combination *in vivo*, we reduced the doses of both drugs. At these lower doses, ceritinib alone did not significantly reduce tumor size, but palbociclib still effectively suppressed tumor growth effectively (**Fig. 7A**). Furthermore, the combination of palbociclib and ceritinib significantly suppressed tumor growth even further compared to the palbociclib-treated tumors (**Fig. 7A**). Histologically, ceritinib-treated tumors resembled vehicle-treated tumors, while palbociclib– and combination-treated tumors exhibited more karyorrhexis and neuropil (**Fig. 7B**). As in the single-drug experiments, ceritinib, palbociclib, and combination treatment reduced Rb1 phosphorylation (**Fig. 7B**, **Suppl. Fig. S8D**). However, we did not observe a difference in phosphorylation of S6, suggesting that Igf1r inhibition was less effective at this dose.

**Figure 7.**
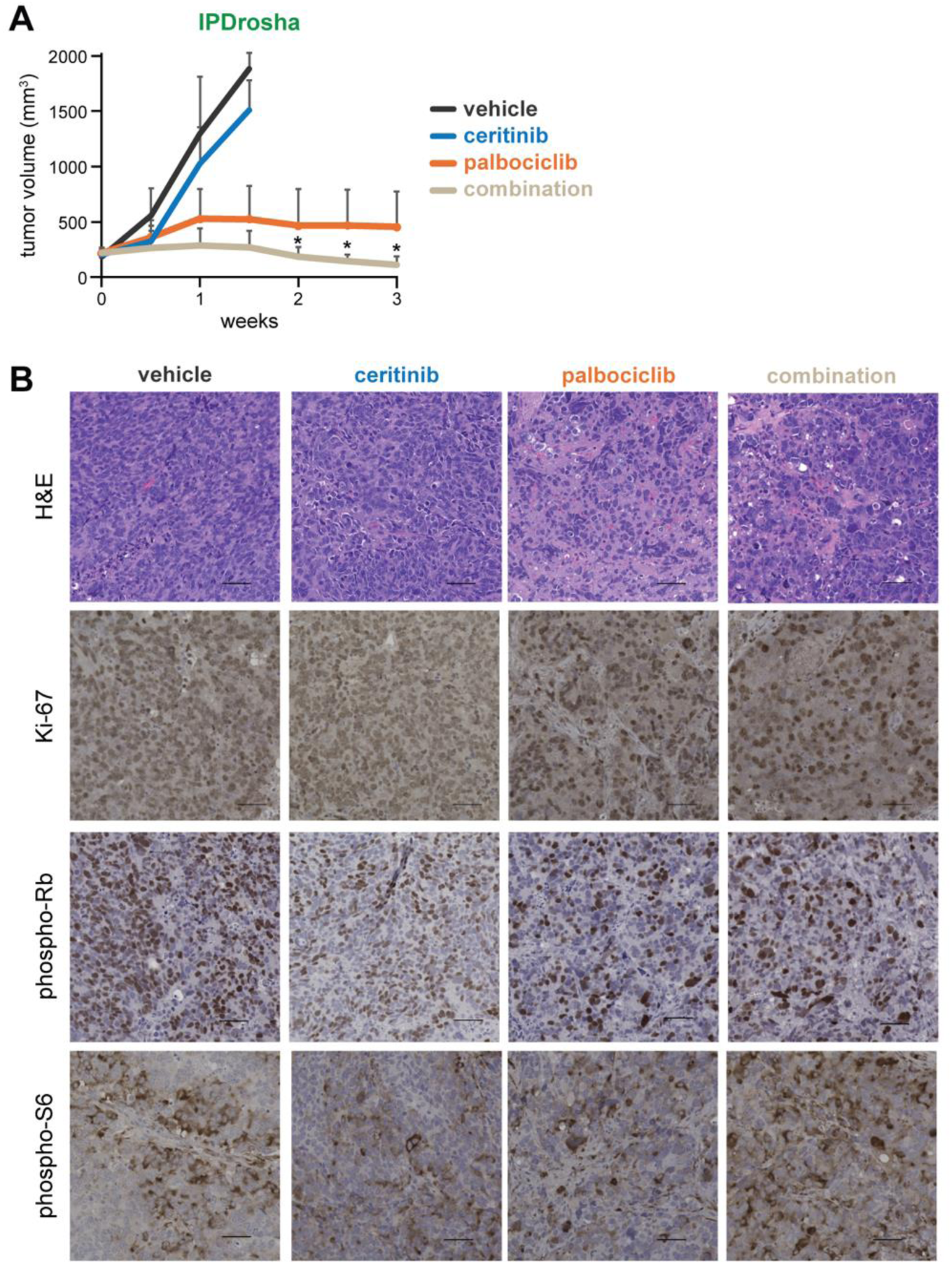
The combination of palbociclib and ceritinib is more effective than either drug alone. **(A)** Tumor volumes for IPDrosha tumors in mice dosed with vehicle, ceritinib, palbociclib, or both drugs (n=8, 7, 7 or 7, respectively). Ceritinib and palbociclib were given at 25 mg/kg/day and 100 mg/kg/day, respectively, by oral gavage, 5 days/week. Values shown are mean ± SD (*p< 0.05; vs. palbociclib; two-sided t-test). **(B)** Photomicrographs of vehicle-, ceritinib-, palbociclib-, or combination-treated tumors (scale 50 µm). Top row shows H&E, the second row shows Ki-67, the third row shows phospho-Rb1 (Ser807/811), and the bottom row shows phospho-S6 (Ser235/236).

### E2F targets are enriched in human pineoblastoma tumors with low DROSHA

Lastly, we examined RNA-seq data from human pineoblastoma tumors to assess whether IPDrosha/IPDicer1 pineal tumors resemble the human disease. Among these tumors, we noted that *DROSHA* expression was essentially undetectable in two of nine tumors (**Fig. 8A**). Other pineoblastoma driver genes (*DICER1*, *DGCR8*, *RB1*, *MYC*) were expressed at comparable levels across these nine tumors (**Suppl. Fig. S9A**). We then compared expression patterns in these two “*DROSHA*-low” tumors to the other seven “*DROSHA*-high” tumors.

**Figure 8.**
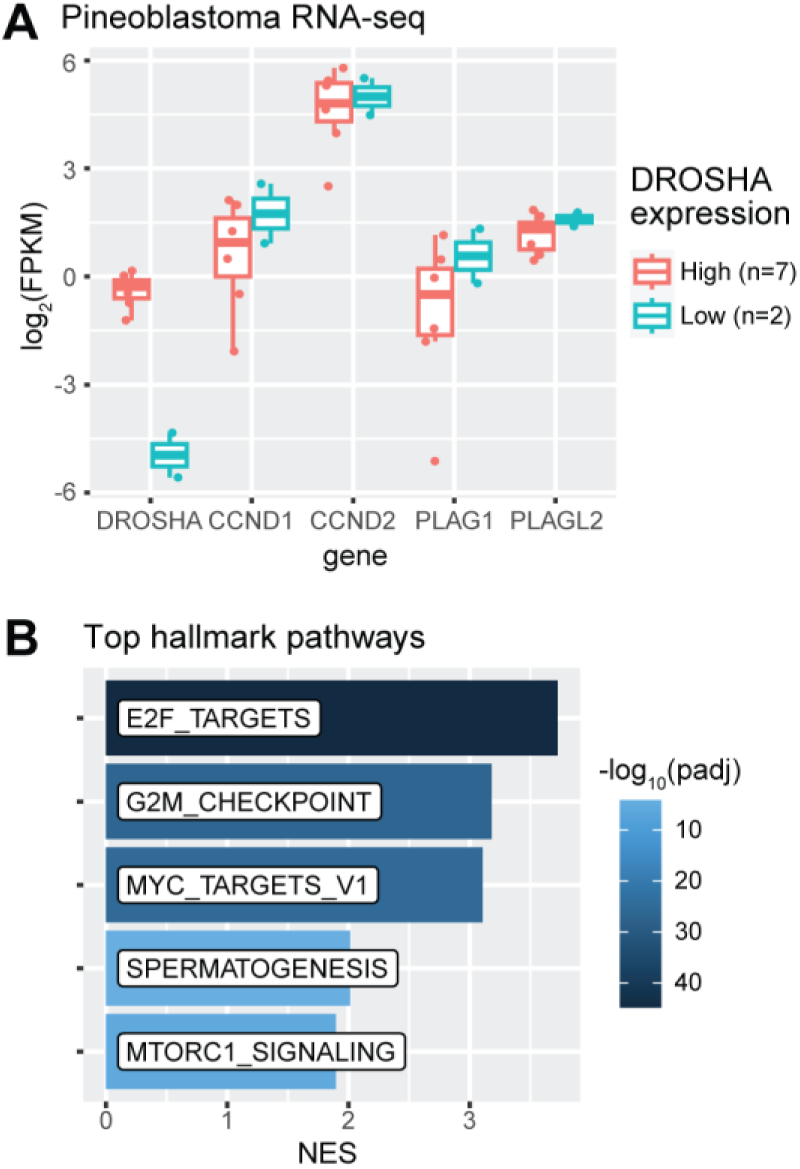
Low *DROSHA* expression is correlated with enrichment for E2F target genes in human tumors. **(A)** Expression of *DROSHA*, *CCND*s, and *PLAG* family genes in *DROSHA*-low vs. *DROSHA*-high human pineoblastomas from St. Jude Cloud RNA-seq (center line, median; box limits, upper and lower quartiles; whiskers, 1.5x interquartile range). FPKM, fragments per kilobase per million reads. **(B)** Top “hallmark” gene sets enriched in *DROSHA*-low pineoblastoma tumors.

Consistent with impairment in microRNA processing, we found that *DROSHA*-low tumors are enriched for genes predicted to be targeted by let-7/miR-98-5p and other microRNAs (**Suppl. Fig. S9B,C**). Furthermore, four of the top five most enriched “hallmark pathways” in these *DROSHA*-low tumors overlapped with the most enriched hallmark pathways in IPDrosha/IPDicer1 tumors (**Fig. 8B**, see **Fig. 3A**). Specifically, the single most enriched hallmark gene set in *DROSHA*-low tumors is “E2F targets” (**Fig. 8B**, **Suppl. Fig. S9D**, **Suppl. Data 4**). Lastly, although they did not reach statistical significance, the two *DROSHA*-low tumors expressed slightly higher levels of *CCND1, CCND2*, *PLAG1*, and *PLAGL2* (**Fig. 8A**). (*IGF2* was not consistently detected in these tumors). In sum, these data in human tumors are consistent with findings from our mouse model that impaired microRNA processing dysregulates the G1/S checkpoint.

## DISCUSSION

*Drosha* loss leads to de-repression of microRNA target genes, which has cascading effects on the expression of other genes. In this study, we dissect *Drosha*-dependent gene networks with a combination of microRNA sequencing, RNA-sequencing, miR-eCLIP sequencing and histone profiling. Our study shows that loss of *Drosha* drives pineal tumors by mimicking *Rb1* loss. Together, these studies revealed that de-repression of direct microRNA targets (such as *Plagl2* and *Ccnd2*) produces secondary effects that drive tumor proliferation (overexpression of *Igf2* and E2F target genes). Small-molecule inhibitors of these indirect effects can block tumor growth (**Suppl. Fig. S9E**). In other words, while mutations in *DROSHA* and *DICER1* cannot be therapeutically targeted currently, the activation of oncogenic pathways driven by microRNA target genes could be blocked with available drugs.

Our results shed light on the cells from which pineoblastoma can arise. IPDrosha, IPDicer1, and IPRb1 tumors express high levels of *Pax6*, *Otx2*, *Crx*, and *Rax* – the homeobox transcription factors that cooperate to regulate pinealocyte development. In particular, *Pax6* and *Otx2* are expressed in rapidly dividing cells at the earliest stages of pineal development [30, 60]. *Otx2* regulates the expression of *Crx*, a specific marker for retina and the pineal gland that may even be used as a diagnostic marker for retinoblastoma and pineoblastoma [19, 61–63]. *Otx2* and *Crx* recognize the same binding motif and thus activate the same target genes, including *Irbp* and *Rax* [30, 64, 65]. In fact, *OTX2* and *CRX* are key oncogenic drivers in group 3 medulloblastoma and are likely to play similar roles in pineoblastoma [66, 67]. Together, these results suggest that pineoblastomas driven by loss of *Drosha*, *Dicer1*, or *Rb1* arise from an embryonic pineal progenitor marked by expression of *Pax6*, *Otx2*, *Crx*, *Rax*, and *Irbp*.

Using a different Cre driver initially developed for the mammary gland, Chung et al. previously showed that *Dicer1* ablation also drives development of pineal tumors; to our knowledge, our study is the first report of pineal tumors arising in mice through loss of *Drosha* [26, 68]. Our findings recapitulate many of their findings, including incomplete penetrance and delayed onset compared with *Rb1*-driven tumors, suggesting that tumors from the two different Cre drivers represent the same cell of origin. Incomplete penetrance is also observed in humans with germline pathogenic variants in *DICER1*. Only ∼19% of these patients develop any kind of cancer, and pineoblastoma is rare even in this population [69]. It is thought that the rate-limiting step in development of *DICER1*-related cancers in this population is the somatic “second-hit” in certain cell types during a vulnerable stage of development [9]. In our model, based on expression of embryonic pineal markers, we surmise the window of vulnerability to be a restricted period of prenatal development, and the number of cells at risk in a developing mouse pineal gland is small. Loss of *Rb1* may cause unrestrained proliferation in the next cell division, but loss of *Drosha* or *Dicer1* may require multiple steps may need to occur within a narrow window before they can impact cell proliferation: depletion of residual microRNAs, de-repression of microRNA target genes, and deployment of secondary effects. Future studies will be needed to identify whether overexpression of any individual microRNA target genes such as *Plagl2* is necessary for pineal tumor development.

Like previous studies of murine pineoblastoma, p53 loss was required for tumor development in our model [23, 25, 27, 45]. These previous studies showed that when pineal cells are forced to proliferate, p53-induced senescence restrained progression into a pineal tumor [23, 45]. Loss of p53 allowed tumor formation by circumventing this senescence. However, while *TP53* is commonly mutated in other *DICER1*-driven human cancers [70, 71], the role of such mechanisms is unclear in pineoblastoma, which rarely exhibits loss or mutation of *TP53.* Human pineoblastoma may use other mechanisms to bypass oncogene-induced senescence, such as loss or silencing of *CDKN2A* (which encodes p16^INK4A^ and p14^ARF^) or amplification of *MDM2* [72]. Larger genomic studies in patient biospecimens, combined with existing conditional mouse models, could be used to study the role of other senescence mechanisms in pineoblastoma formation.

Previous studies showed that pineal tumors in mice can arise through dysregulation of the G1/S checkpoint, either by ablating *Rb1* or upregulating *Ccnd1* [23, 45]. By demonstrating that IPDrosha/IPDicer1 tumors drive proliferation by upregulating D-type cyclins, we build upon these studies and show similarities between loss of *Drosha/Dicer1* and loss of *Rb1*. CDK4/6 inhibitors are clinically approved for use in hormone receptor-positive breast cancer and under investigation for treatment of other undifferentiated pediatric and adult tumors that retain intact RB1-E2F responses [20, 44, 73, 74]. Specifically, CDK4/6 inhibitors have been found to induce differentiation in neuroblastoma [75], rhabdomyosarcoma [22], medulloblastoma [76], and prostate cancer [77]. To date, the only known genomic predictors of sensitivity to CDK4/6 inhibition directly drive overexpression of *CCND1*, *CCND2*, or *CCND3* [78]. These may arise through translocation, amplification, or variants that remove or alter microRNA binding sites in the 3’ UTR. Our results suggest that removal of the microRNAs themselves could also contribute to overexpression of D-type cyclins and thus sensitivity to CDK4/6 inhibitors.

By identifying direct microRNA targets with miR-eCLIP, we showed that proliferation of IPDrosha/IPDicer1 tumors depends on a combination of the direct and indirect effects of microRNA loss. Our study also shows the limitations of microRNA target prediction. We used TargetScan, which has been shown to outperform other microRNA prediction algorithms [79, 80]. Nevertheless, identifying direct targets by miR-eCLIP was more predictive of gene expression changes than using computational predictions alone. Further refinements to these prediction algorithms may be needed to account for long-term *in vivo* effects.

In this study, we focus on *Plagl2* and *Ccnd2*, but the roles of other microRNA target genes should also be considered. For instance, we also found that MYC target genes were enriched in the IPDrosha/IPDicer and IPRb1 tumors, and many microRNAs regulate *Mycn*. Human pineoblastomas not driven by loss of *RB1*, *DROSHA*, or *DICER1* are commonly driven by *MYC* activation [19]. Similarly, a recently published mouse model driven by oncogenic *Dicer1* mutations led to upregulation of E2F and Myc target genes [81]. In other words, *MYC* activation may be a common downstream effect of these mutations.

We have also previously reported overlapping results in Wilms tumor, a pediatric cancer of embryonic renal progenitors frequently driven by mutations in microRNA processing genes [3–7]. Unlike the complete loss of microRNA production seen in pineoblastoma, Wilms tumors most commonly exhibit point mutations in *DROSHA* that lead to incomplete impairment of microRNA processing. In two separate studies, we previously found that *DROSHA/DICER1* regulate both *PLAG1/IGF2* and *CCND2* in Wilms tumor [82, 83]. In the first study, we showed that *PLAG1* was overexpressed in Wilms tumors with mutations in *DROSHA* or *DICER1*, and that *DICER1*-knockout Wilms tumor cells overexpress *PLAG1* and *IGF2* in a microRNA-dependent manner. In the second study, we found that *CCND2* was the single most overexpressed gene in Wilms tumor cells when *DROSHA* was silenced, and that mutated microRNA processing enzymes led to upregulation of E2F target genes. Together, our observations suggest that despite arising from different cell types and through different effects on microRNA processing, *DROSHA*-mutant Wilms tumor and pineoblastoma harness similar pathways to drive proliferation of embryonic progenitors.

It remains unclear why pineoblastoma tolerates complete loss of *DROSHA* or *DICER1*, while other tumors, including pleuropulmonary blastoma, Sertoli-Leydig cell tumor, and rhabdomyosarcoma, arises through point mutations in *DICER1* [9, 10]. These point mutations affect the metal-binding pocket of the RNase IIIb domain, which cleaves the 5p arm of pre-microRNA hairpins, resulting in loss of 5p-derived microRNAs but retention of 3p-derived microRNAs. Many of these *DICER1*-related tumors also resemble highly proliferative, undifferentiated progenitors. Furthermore, many of the microRNAs we identified as key regulators of gene expression in pineoblastoma are 5p-derived and thus lost in tumors with *DICER1* missense variants, including members of the let-7/miR-98-5p, miR-99/100-5p, and miR-125/352-5p families. Our results suggest that loss of let-7/miR-98-5p microRNAs is a key driving force in *Drosha/Dicer1*-driven pineoblastoma formation, and it is unclear whether loss of 3p-derived microRNAs is.

Recently, a subset of embryonal brain tumors was found to be defined by amplification and overexpression of *PLAGL2*, and our study expands on its oncogenic importance [84]. *PLAGL2* was initially described as a homolog of the *PLAG1* transcription factor, whose translocation-driven overexpression defines pleomorphic adenoma of the salivary glands [85]. In the embryonic brain and in brain tumors, *Plagl2* regulates the balance between differentiation and proliferation, and its overexpression shifts neural progenitors towards proliferation [86, 87]. Outside the brain, *PLAGL2* also contributes to oncogenesis in lung cancer [88, 89], colorectal cancer [90–93], gastric cancer [94], breast cancer [95, 96], and acute myeloid leukemia [97]. In these cancers, *PLAGL2* has been shown to contribute to proliferation, epithelial-mesenchymal transition, and “stemness” by regulating target genes such as *Igf2* and regulators of Wnt signaling [84, 87, 90, 91, 93, 96]. To our knowledge, however, this is the first description of *PLAGL2* upregulation in *DROSHA*-or *DICER1*-related pediatric cancers.

Our studies used a subcutaneous allograft model over an orthotopic model for practical reasons. Hundreds of subcutaneous patient-derived xenografts of pediatric cancer have been used to test dozens of drugs in preclinical studies, and response rates in these studies commonly correlates with clinical response [98, 99]. In our study, using a subcutaneous model allowed us to test the principle that these small molecules are capable of blocking growth of *Drosha/Dicer1*-driven cancers. CDK4/6 inhibitors and Igf1r inhibitors have been shown to cross the blood-brain barrier [100–103]. Before such drugs are tested in a clinical setting, response in an orthotopic model and further validation of these correlations in human tumors may be required.

In this study, we identify two small molecules that impaired tumor proliferation by blocking signaling pathways normally regulated by microRNAs. Although they are both clinically approved, it remains to be seen whether they will be effective in children with these cancers. In our models, neither drug was cytotoxic on its own, and they may need to be combined with other therapies to harness clinical benefit. For instance, previous studies have found synergy between CDK4/6 inhibitors and immune checkpoint blockade, and we also found that palbociclib upregulated immune gene signatures [52–55] (**Suppl. Fig. S4E**). Alternatively, human tumors may harness other oncogenic signaling pathways or growth factors, such as RAS/MAP kinase; indeed, RAS pathway genes are the most frequently mutated genes in *DICER1*-related tumors after *TP53* [71, 104, 105]. Future studies will be needed to identify the specific mitogenic signaling pathways used in human pineoblastomas. Nevertheless, our results demonstrate the potential for the signals upregulated by microRNA loss to be therapeutic targets in cancers driven by impairment of *DROSHA* or *DICER1*.

## METHODS

### RNA-seq

Flash frozen mouse tissue was disrupted using an electric pestle and homogenized using a Qiashredder column (Qiagen, cat. no. 79654). Total RNA was extracted using the miRNeasy mini kit (Qiagen, cat. no. 217004). For IPDrosha, IPDicer1, and IPRb1 pineal tumors, single-end whole transcriptome sequencing was performed at the UT Southwestern McDermott Next Generation Sequencing (NGS) Core. For palbociclib and ceritinib treated tumor allografts, single-end strand-specific mRNA sequencing was performed at the UT Southwestern McDermott Next Generation Sequencing (NGS) Core. For IPDrosha tumors, matched normal brain, and age-matched normal adult pineal glands, paired-end whole transcriptome sequencing was performed at DNA Link Inc. For control samples from normal adult pineal glands, pineal glands were pooled from two age– and genotype-matched adults to ensure sufficient RNA sample for sequencing.

Reads from each sample were mapped to the reference genome using STAR (v2.5.3a) [106], and read counts were generated using featureCounts [107]. Differential gene expression calculations were then computed from raw counts using the DESeq2 package (v1.42.0) [108]. The Wald statistic output for protein-coding genes from DESeq2 was used as input for gene set enrichment analysis (GSEA) using the fgsea package (v1.28.0) [109]. Hallmark gene sets were derived from MSigDB v2023.1.Mm or v2023.1.Hs for mouse or human, respectively [32]. Predicted microRNA target genes were defined as genes with Aggregate P_CT_ ≥ 0.9 and Total context++ score < –0.4 based on TargetScan (v8.0) [42].

For external data, RNA-seq FPKM tables from rat pineal glands across different stages of development were downloaded from GEO dataset GSE46127 [110]. Differential expression was calculated using t-tests after log_2_ transformation. Marker genes at each timepoint were defined as the 300 most overexpressed protein-coding genes with mouse homologs, compared to other timepoints.

For comparing our data to external mouse tumor RNA-seq, counts tables were downloaded from the following GEO datasets, and differential expression analysis was performed as described above: glioblastoma (GSE151414) [111], medulloblastoma (GSE235625) [112], MYCN-driven glioma (GSE227289) [113], H3K27M glioma (GSE95168) [114], and neuroblastoma (GSE230264) [115]. For these analyses, proliferation markers were defined as the 587 unique genes in the following MSigDB gene sets: HALLMARK_E2F_TARGETS, HALLMARK_G2M_CHECKPOINT, HALLMARK_MITOTIC_SPINDLE, and HALLMARK_DNA_REPAIR. Proliferation-independent embryonic pineal markers were defined by taking the top 200 embryonic pineal markers from the rat RNA-seq dataset and removing the 24 genes that overlapped with the proliferation markers. Similarly, proliferation-independent adult pineal markers were defined by taking the top 200 adult pineal markers from the rat RNA-seq dataset and removing the 2 genes that overlapped with proliferation markers.

For correlating with human pineoblastoma RNA-seq, counts tables from pineoblastoma patient samples were obtained from the St. Jude Cloud data repository (https://www.stjude.cloud) [116] and processed as above.

### Quantitative PCR

Flash-frozen mouse tissue was disrupted using an electric pestle and homogenized using a Qiashredder column (Qiagen, cat. no. 79654). RNA was extracted using the miRNeasy mini kit (Qiagen, cat. no. 217004), and cDNA was prepared using RT^2^ HT First Strand kit (Qiagen, cat. no. 330411) or iScript Reverse Transcription Supermix for RT-qPCR (Bio-Rad, cat. no. 1708840). iTaq Universal SYBR Green Supermix (Bio-Rad, cat. no. 1725125) was used for qPCR in four replicates per condition. The oligos used for qPCR are referenced in **Supplementary Table 1**.

For miRNA TaqMan qPCR, total RNA was extracted using the miRNeasy mini kit (Qiagen, cat. no. 217004). cDNA synthesis was performed with the TaqMan MicroRNA Reverse Transcription kit (Applied Biosystems, cat. no. 4366596). The following assays were used for cDNA synthesis and qPCR detection (ThermoFisher cat. no. PN4427975): hsa-let-7a (Thermo Fisher, assay id 000377), hsa-let-7b (Thermo Fisher, assay id 002619), hsa-let-7i (Thermo Fisher, assay id 002221), hsa-miR-100 (Thermo Fisher, assay id 000437), mmu-miR-183 (Thermo Fisher, assay id 002269), and U6 snRNA (Thermo Fisher, assay id 001973). TaqMan Universal PCR Master Mix (Applied Biosystems, cat. no. 4304437) was used for qPCR, and each sample had three replicates per condition. Each miRNA assay was normalized to U6 and displayed as 2^-ΔΔCt^, and p values were calculated using two-tailed *t* tests.

### Small RNA sequencing

For IPDrosha and IPRb1 tumors, total RNA was submitted for small RNA sequencing to the UT Southwestern McDermott NGS Core using the NEBNext Small RNA kit (New England Biolabs, cat. no. E7330), with size selection between 15bp and 30bp. Reads were aligned and counted using miRge3.0 [117]. Filtered reads aligning to microRNAs were normalized to reads aligning to tRNAs and snoRNAs as an internal control.

For small RNA sequencing and analysis from IPDrosha tumors, pineal gland, and brain samples, total RNA was submitted to DNA Link. QIAseq miRNA Library QC Spike-ins (Qiagen cat. no. 331535) were used as a spike-in control. Analysis was performed using mirDeep2 [118] for alignment, quantification, and annotation. Counts for each microRNA were normalized to spike-in control, and microRNAs of the same family were grouped per TargetScan [42]. Known mirtrons were annotated from mirtronDB [119]. Differential expression for individual microRNAs was performed using t-tests of log_2_-transformed, normalized read counts.

### CUT&RUN

Pineal tumors from IPDrosha/IPDicer1 mice were dissociated into single-cell suspension using the Brain Tumor Dissociation Kit (P) (Miltenyi Biotec, cat. no. 130-095-942), and then “Cleavage Under Targets and Release Using Nuclease” (CUT&RUN) [120, 121] libraries were prepared from dissociated tumor cells using the ChIC/CUT&RUN Kit (EpiCypher, cat. no. 14-1048) according to manufacturer recommendations. Briefly, single-cell suspensions were immobilized onto beads, permeabilized, and incubated in Epicypher antibody buffer at 1:50 for H3K4me3, H3K27ac, or Rabbit IgG. (See **Supplementary Table 2** for more details.) Target-DNA complexes were cleaved using pAG-MNase, and libraries were prepared using NEBNext Ultra II DNA Library Kit from Illumina (New England Biolabs, cat. no. E7645S).

Library quality checks and sequencing were performed at the UT Southwestern McDermott NGS Core. Sequencing reads were analyzed using the nf-core/cutandrun bioinformatic pipeline v2.0 [122] on the UT Southwestern BioHPC Astrocyte platform. Briefly, after trimmed reads are aligned to the target genome and the spike-in using Bowtie 2 (v2.4.4) [123], peaks are called using the SEACR (v1.3) [124] and MACS2 (v2.2.7) [125] pipelines. Downstream visualization is performed using the Integrative Genomics Viewer (IGV, v2.17.1) [126], deepTools (v3.5.1) [127], or Gviz (v1.46.1) [128] toolkits.

### miR-eCLIP sequencing

Two frozen IPRb1 tumor fragments underwent miR-eCLIP sequencing at EclipseBio, as previously described [129]. Briefly, Ago2 was immunoprecipitated from crosslinked RNA. After ligation of miRNA-mRNA chimeras, input and immunoprecipitated RNA were analyzed by sequencing on the Illumina NovaSeq 6000 platform. Chimeric and non-chimeric eCLIP sequences were processed using the pipeline described [129] and compared to input in each sample using the CLIPper algorithm (v2.0) [130]. Chimeric and non-chimeric peaks were compared between the two samples to identify reproducible peaks. Reproducible non-chimeric peaks were those enriched in each IP with a log_2_-fold-change ≥ 3 and p value ≤ 0.001. HOMER *de novo* motif analysis [131] was used to identify enriched motifs in these peaks. Reproducible microRNA-miRNA chimeras were those with ≥3 chimeric reads and a reproducibility index ≥ 0.5. Seed matches were then limited to perfect-match, canonical 7-or 8-mers (based on TargetScan definitions [42]) in the 3’UTRs of their target genes.

### Immunohistochemistry

Formalin-fixed tissues were processed at the UT Southwestern Tissue Management Shared Resource or Mouse Histology core, and fixed sections were deparaffinized using Histo-Clear II (VWR, cat. no. 101412-884) or xylene (Sigma, cat. no. 534056) with gradient ethanol washes. Heat antigen retrieval was performed using Trilogy (Sigma, cat. no. 920P) or 1M citrate buffer pH 6.0. Tissue slides were stained with Ki-67, phospho-Rb, synaptophysin, phospho-S6, and GFAP (see **Supplementary Table 2** for dilutions and catalog numbers). Signal was obtained using diaminobenzadine (DAB) (Millipore, cat. no. 281751), and hematoxylin (Sigma Aldrich, cat. no. GHS 132-1L) was used for counterstaining.

For immunohistochemical quantification, biological replicates (n=2) were imaged for each condition, and representative images of each slide were captured at 20x. Adjacent cells (n=150 cells per slide) in a localized region were counted using the manual cell counter function in the QuPath software package [132]. The results from both slides were combined to calculate the proportion of positively-stained cells, and chi-square analysis was used to calculate p values.

### ChIP-qPCR

ChIP-qPCR was performed using the ChIP-IT High Sensitivity kit (Active Motif, cat. no. 53040). Flash-frozen tumors were fixed with 1% formaldehyde, and 6.75 µg chromatin was sonicated using a Diagenode Bioruptor and then immunoprecipitated with 2.5 µg of primary antibodies recognizing Plagl2 (Sigma Aldrich, cat. no. SAB3500815) or Rabbit IgG (Cell Signalling Technology, cat. no. 2729). Control input chromatin was also prepared with the same kit. Quantitative PCR was performed with iTaq Universal SYBR Green Supermix (Bio-Rad, cat. no. 1725125), and each sample had three replicates per condition. The oligos used for qPCR are shown in **Supplementary Table 1**. Results were calculated as percent input, and p values were calculated using two-tailed *t* tests.

### Luciferase reporter assay

Starting from the pcDNA3.1-acGFP plasmid (Addgene 128047), we replaced the GFP coding sequence with the *Plagl2* cDNA using the XbaI and NotI restriction enzymes. We obtained the pGL4.26[luc2/minP/Hygro] minimal promoter firefly luciferase and pGL4.75[hRluc/CMV] *Renilla* luciferase plasmids from Promega (cats. E8441 and E6931). We amplified regions 1 and 2 of the *Ccnd2* promoter from mouse genomic DNA using the primers listed in **Suppl. Table 1** and cloned them into the pGL4.26 plasmid using the XhoI and KpnI restriction enzymes. The *Igf2* promoter was synthesized as a dsDNA gene fragment by Twist Bioscience (sequence given in **Suppl. Table 1**) and cloned into the pGL4.26 plasmid using the XhoI and HindIII restriction enzymes.

To perform the luciferase reporter assay, we seeded HEK293 cells in 96-well plates. The next day, we co-transfected each well with 50 ng of pGL4.26 (with varying promoters), 50 ng of pcDNA3.1 (with or without Plagl2 cDNA), and 1 ng of pGL4.75 (as transfection control) using Lipofectamine 3000 (Thermo Fisher, cat. L3000015). We assessed luciferase activity 72 hours post-transfection using Dual-Glo Luciferase Assay System (Sigma, cat. no. E2920). We performed six replicates per condition and normalized the firefly signal to the *Renilla* signal within each well, and we compared conditions using two-sided t-tests.

### Western blots

Whole cell lysates were obtained from flash-frozen tumor tissue using RIPA lysis buffer (Sigma, cat. no. R0278) with protease and phosphatase inhibitor (Thermo Scientific, cat. no. A32961). The tissue was disrupted with an electric pestle, and lysate was quantified using Pierce BCA Protein Assay (Thermo Scientific, cat. no. 23225). Samples were separated on polyacrylamide gels and transferred to nitrocellulose membranes prior to blocking and probing. Antibodies and dilutions are listed in **Supplementary Table 2**.

### Animal breeding and husbandry

Animal experiments were performed under oversight from the University of Texas Southwestern Institutional Animal Care and Use Committee protocols 2019-102689 and 2019-102680.

Mice of the following strains were obtained from Jackson laboratories: Irbp-Cre (also known as Rbp3-Cre, cat. no. 003967, RRID IMSR_JAX:003967), *Rb1*^flox^ (cat. no. 026563, RRID IMSR_JAX:026563), *Drosha*^flox^ (cat. no. 008592, RRID IMSR_JAX:008592), *Dicer1*^flox^ (cat. no. 006366, IMSR_JAX:006366) and *p53*^flox^ (cat. no. 008462, RRID IMSR_JAX:008462). These were maintained on a C57BL6/J background. Nod-scid gamma (NSG) mice were also purchased from Jackson laboratories (cat. no. 005557, RRID IMSR_JAX:005557) and maintained under the UT Southwestern Immunocompromised Breeding Program. Mice were monitored visually twice weekly for a domed head and given moist chow upon tumor development. Tumors were harvested up to 5 days later. To compare tumor-free survival of mice with different genotypes, p values were calculated using the log-rank test.

### Allografts and in vivo drug treatments

Pineal tumors from IPDrosha/IPDicer1 mice were dissociated using the Brain Tumor Dissociation Kit (P) (Miltenyi Biotec, cat. no. 130-095-942). Male and female NSG mice at 6-8 weeks of age were subcutaneously injected with 0.5-1.5 million tumor cells, suspended in 50% cold PBS and 50% matrigel (Corning, cat. no. CB-40234). Tumor volume was measured by calipers twice a week. When tumors reached 150 mm^3^, mice were administered palbociclib (150 mg/kg, LC Laboratories, cat. no. P-7788) or an equivalent volume of vehicle (50 mM sodium L-lactate, pH 6, Sigma cat. no. 71718) by oral gavage, five days a week for up to 3 weeks. Tumors were harvested when they reached 1,000 mm^3^, 4 hours after the last dose of drug, and rinsed in PBS before flash-freezing or fixation. Tumor sizes were compared using two-tailed t-tests at each timepoint, and survival was compared using the log-rank test.

For ceritinib treatment, pineal tumor cells were injected into NSG mice as described above. Once tumors reached 150 mm^3^, mice were administered ceritinib (50 mg/kg, Sellekchem, cat. no. S4967) or an equivalent volume of vehicle (0.5% methylcellulose and 0.5% Tween-80, Fisher Scientific cat. no. AC428430500 & Sigma cat. no. P8192) by oral gavage, five days a week for up to 3 weeks. Tumors were harvested when they reached 1,000 mm^3^, 3 hours after the last dose of drug, and rinsed in PBS before flash-freezing or fixation.

For combination treatment of palbociclib and ceritinib, pineal tumor cells were injected into NSG mice as described above. Once tumors reached 150 mm^3^, mice were administered palbociclib (100 mg/kg, LC Laboratories, cat. no. P-7788) or an equivalent volume of vehicle (50 mM sodium L-lactate pH 6, Sigma cat. no. 71718) by oral gavage, five days a week for up to 3 weeks. Five hours after each dose of palbociclib or vehicle, mice were administered ceritinib (25 mg/kg, Sellekchem cat. no. S4967) or an equivalent volume of vehicle (0.5% methylcellulose and 0.5% Tween-80, Fisher Scientific cat. no. AC428430500 & Sigma cat. no. P8192) by oral gavage, five days a week for up to 3 weeks. Tumors were harvested when they reached 1,000 mm^3^, 3 hours after the last dose of drug for the single agent treated tumors and 1 hour post the second dose of the combination treated tumors.

### Data availability

Raw and processed sequencing data are available at the GEO SuperSeries GSE255914.

## ACKNOWLEDGMENTS

This work was supported by funding from the V Foundation for Cancer Research (V2022-010 to K.S.C.); National Cancer Institute (K08CA207849 and 1R01CA289259 to K.S.C. and Cancer Center Support Grant P30CA142543); and Cancer Prevention and Research Institute of Texas (RR180071 to K.S.C., RP210041 to C.F.). This research used computational resources provided by the BioHPC supercomputing facility in the Lyda Hill Department of Bioinformatics at UT Southwestern, which is supported by Cancer Prevention and Research Institute of Texas (RP150596), and tissue processing at the UT Southwestern Simmons Comprehensive Cancer Center Tissue Management Shared Resource, which is supported by the National Cancer Institute (P30CA142543). We appreciate the past, current, and future mentoring from James Amatruda, Joshua Mendell, and many others.

## AUTHOR CONTRIBUTIONS

C.R.F., K.D., L.K.M., U.O., T.S., K.O., and K.S.C. conceived and performed experiments. C.R.F., Y.W., and K.S.C. wrote the manuscript. V.R. provided expertise, feedback, and analysis. J.L. and K.S.C. performed data analysis and secured funding.

