## Supplemental Figures for "An imbalance between proliferation and differentiation underlies the development of microRNA-defective pineoblastoma"

### Fig S1. IPDrosha, IPDicer1, and IPRb1 pineal tumors retain embryonic pineal expression pathways.

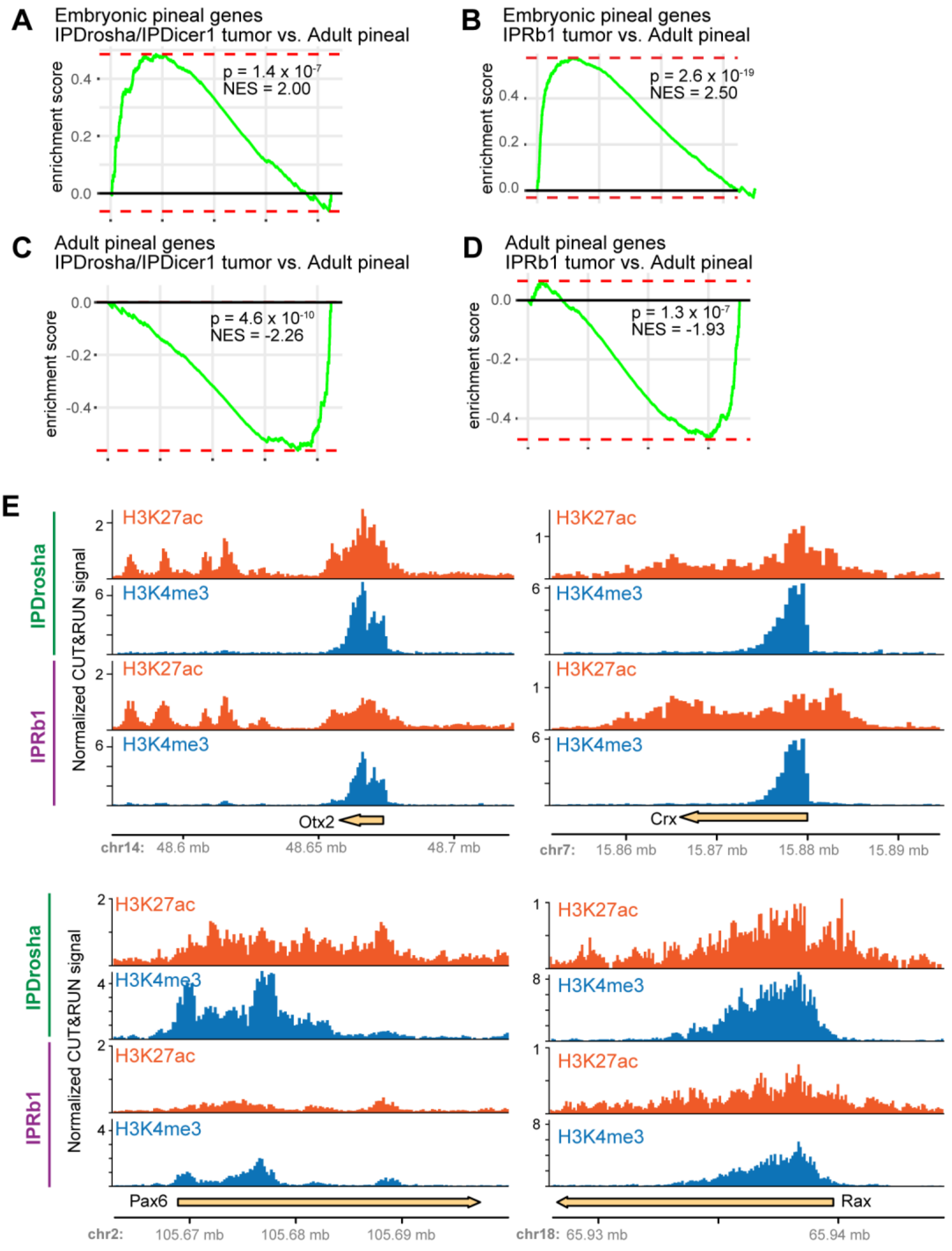

### Fig S2. Comparison of IPDrosha, IPDicer1 and IPRb1 pineal tumors to other murine tumor models.

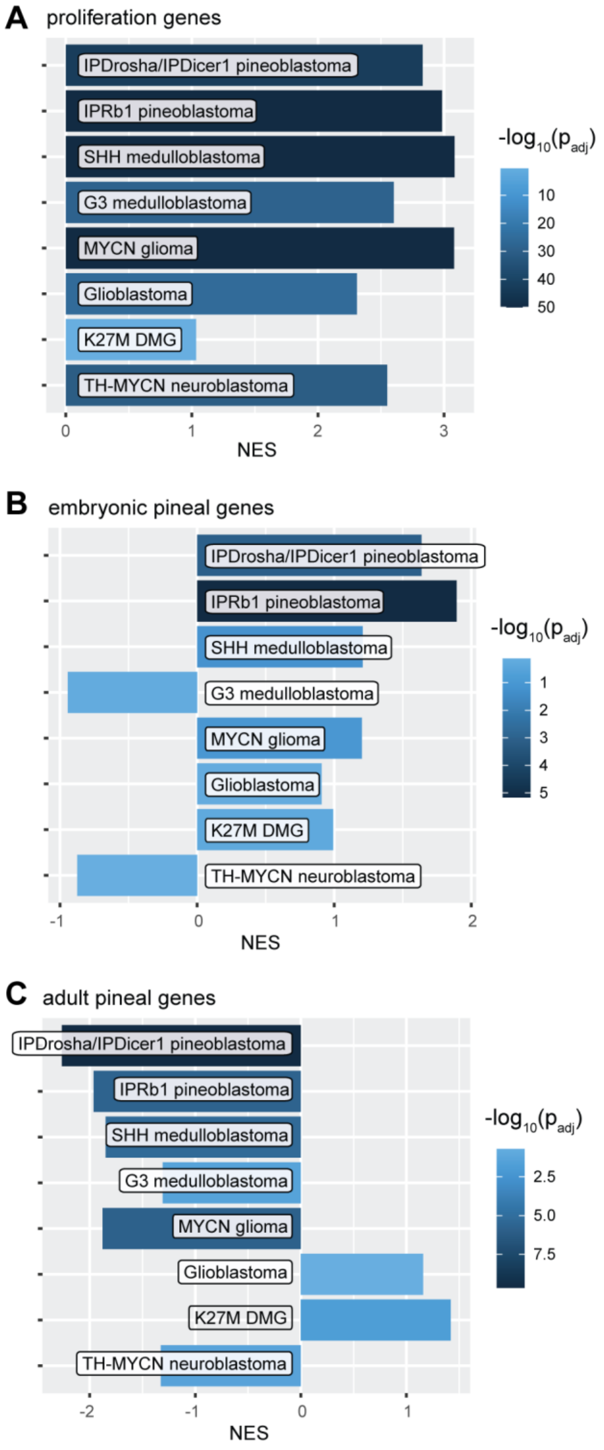

### Fig S3. *Drosha*-driven tumors exhibit loss of canonical microRNAs.

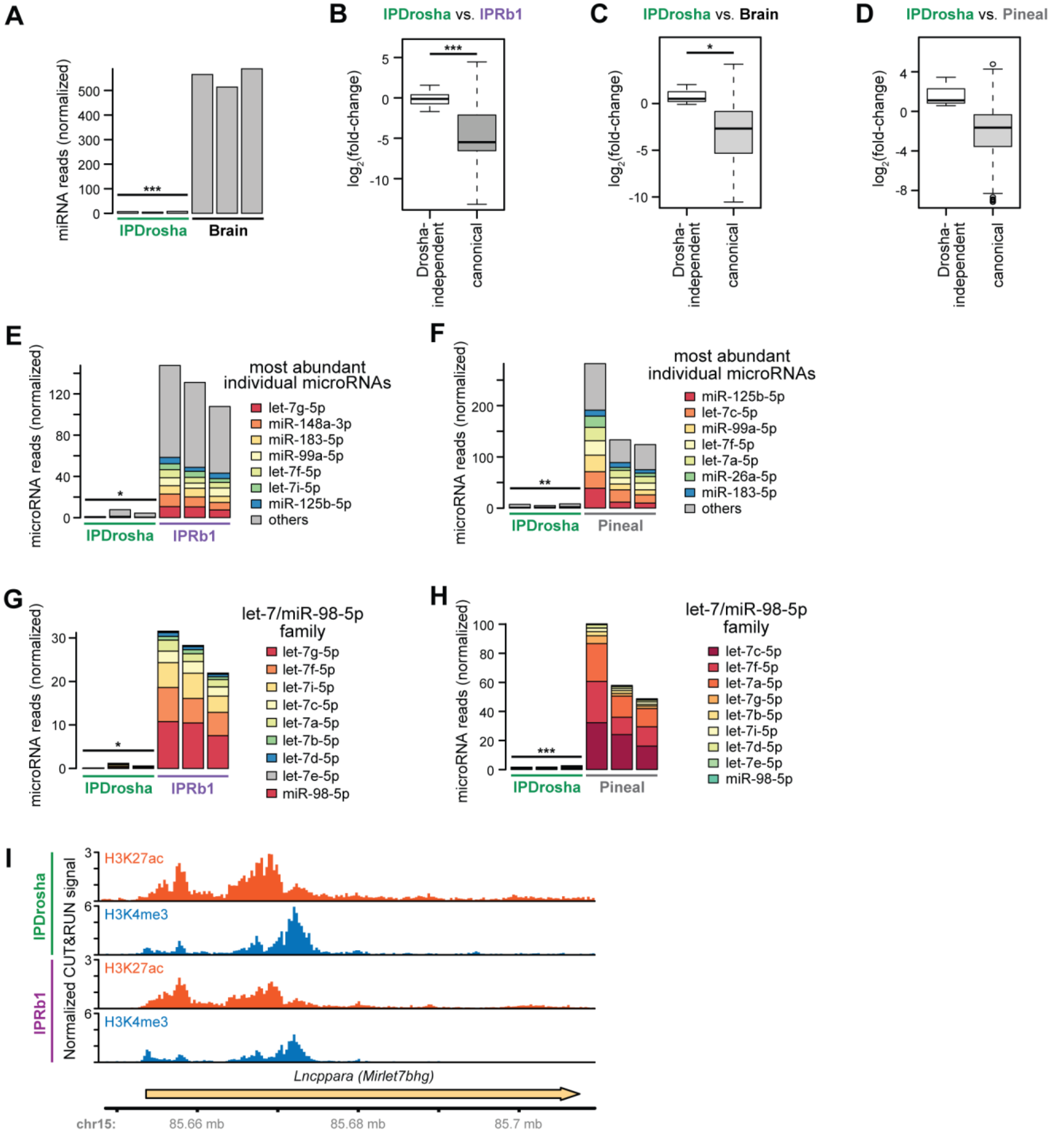

### Fig S4. Patterns of microRNA regulation in miR-eCLIP sequencing of IPRb1 tumors.

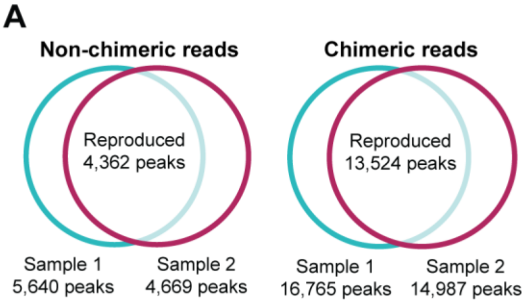


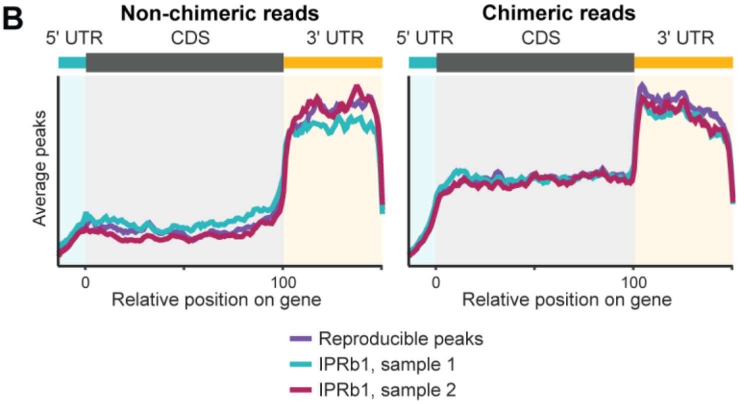


**C** **Top motifs in non-chimeric reads**

| Motif | Seed match | % of peaks with motif | $-log_{10}(p)$ |
|---|---|---|---|
| HWGTGCCTA | miR-124-3p | 8.85% | 748.9 |
| NHHCTACCTCAD | let-7/miR-98-5p | 8.57% | 745.6 |
| ATCTACCTCA | let-7/miR-98-5p | 6.54% | 715.9 |
| CTAAGTGCCTA | miR-124-3p | 8.47% | 709.2 |

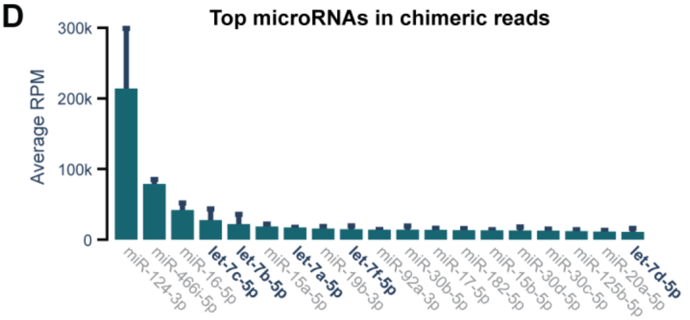


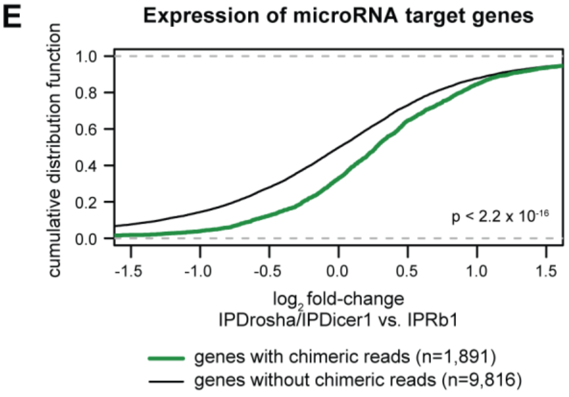


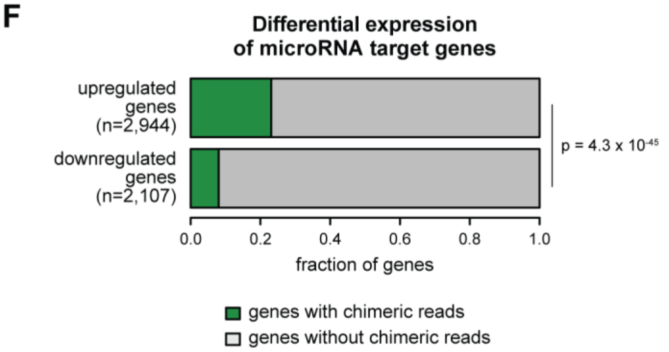

Fig S5. Regulation of *Ccnd2* and *Mycn*.

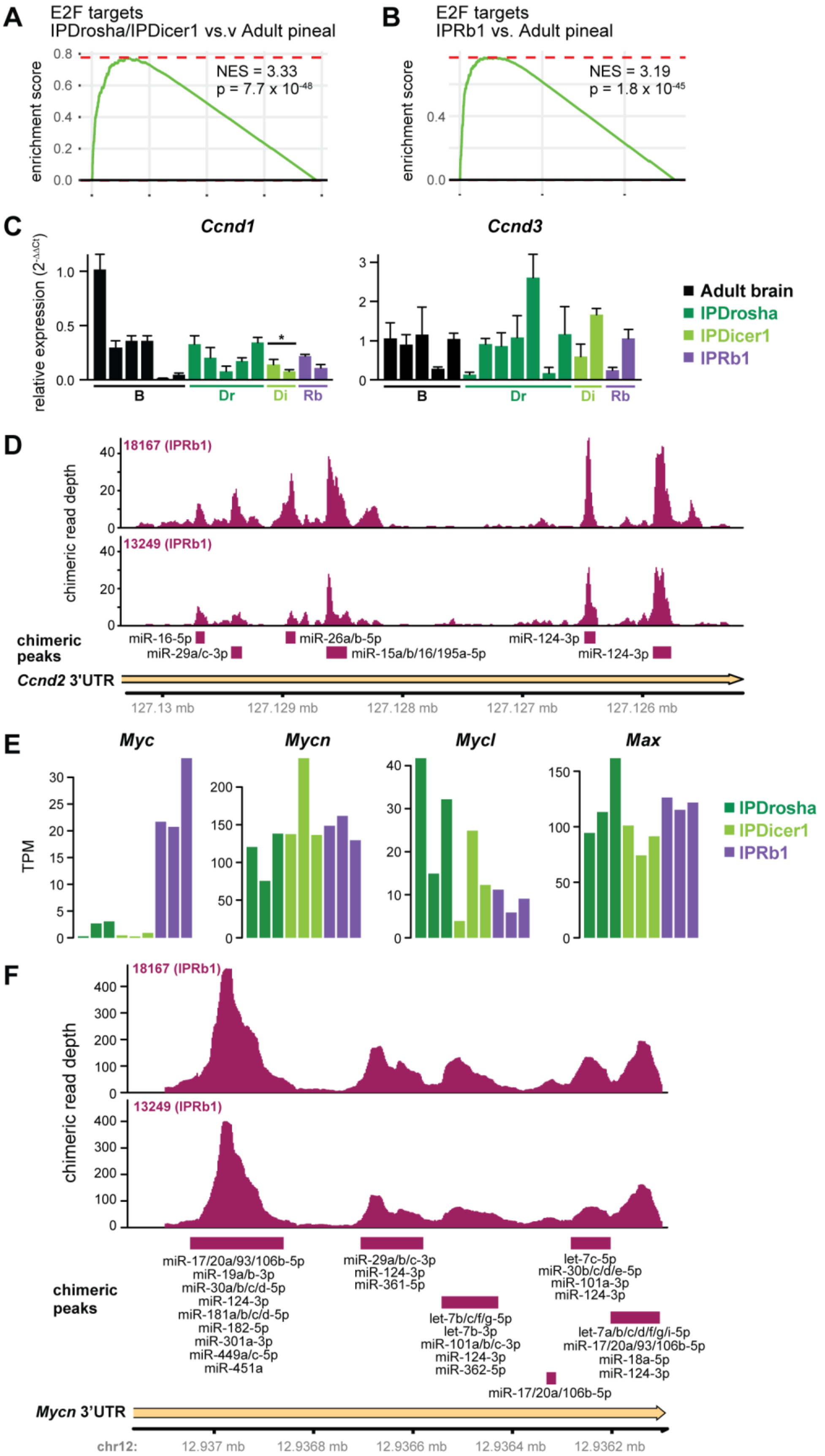

### Fig S6. Palbociclib in IPDrosha/IPDicer1 tumors.

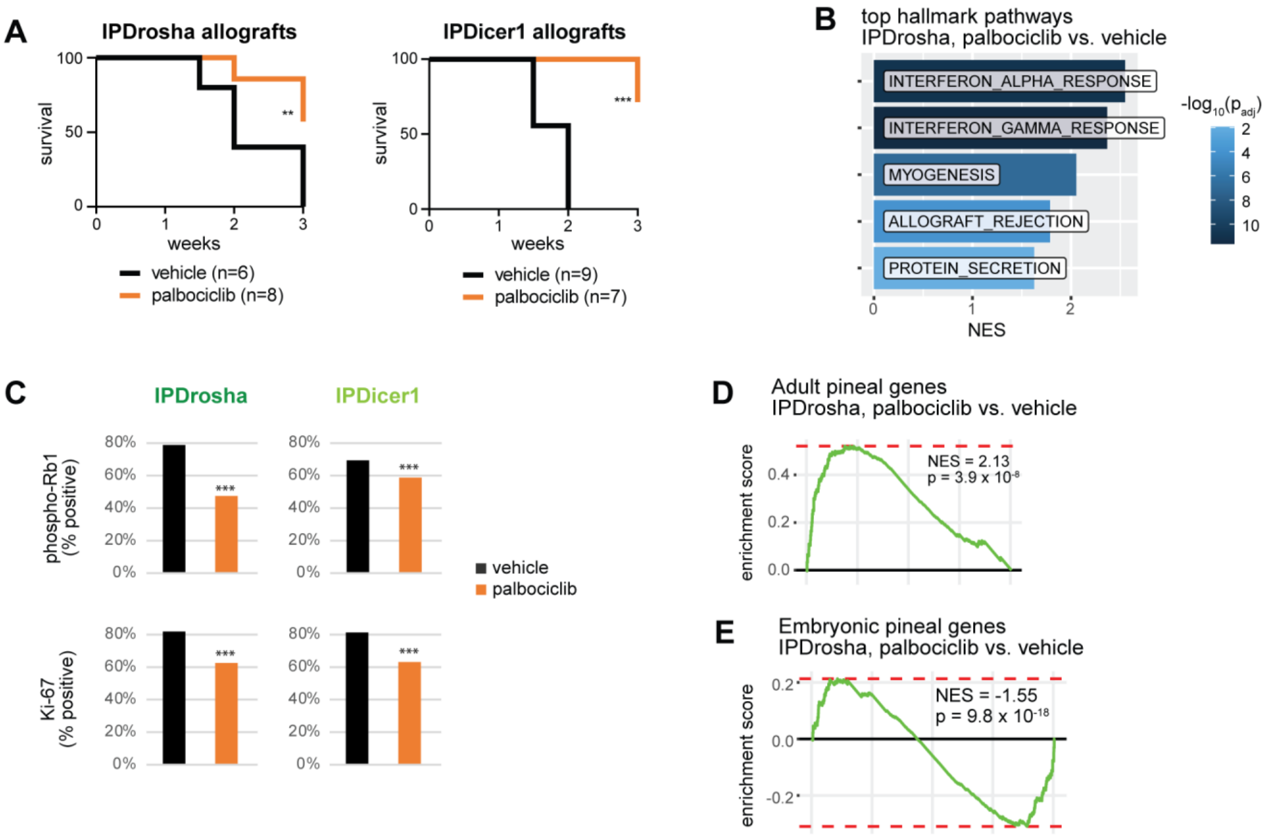

### Fig S7. Regulation of *Plagl2.*

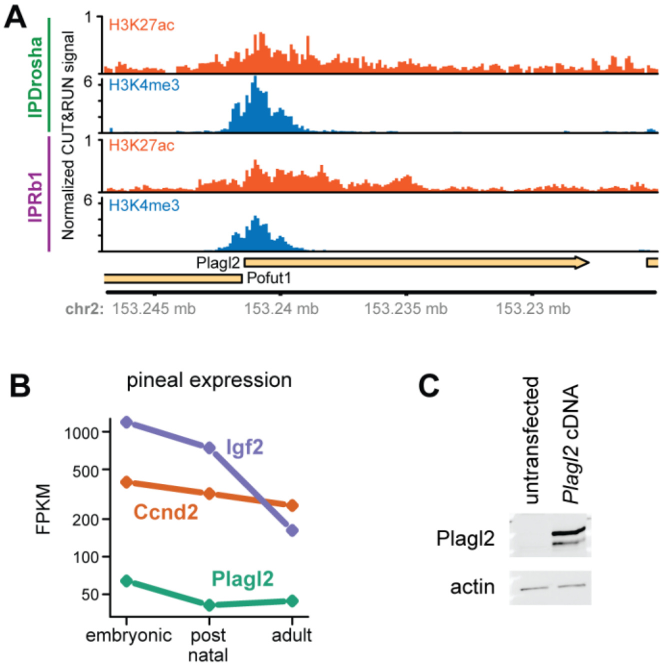

### Fig S8. Ceritinib treatment.

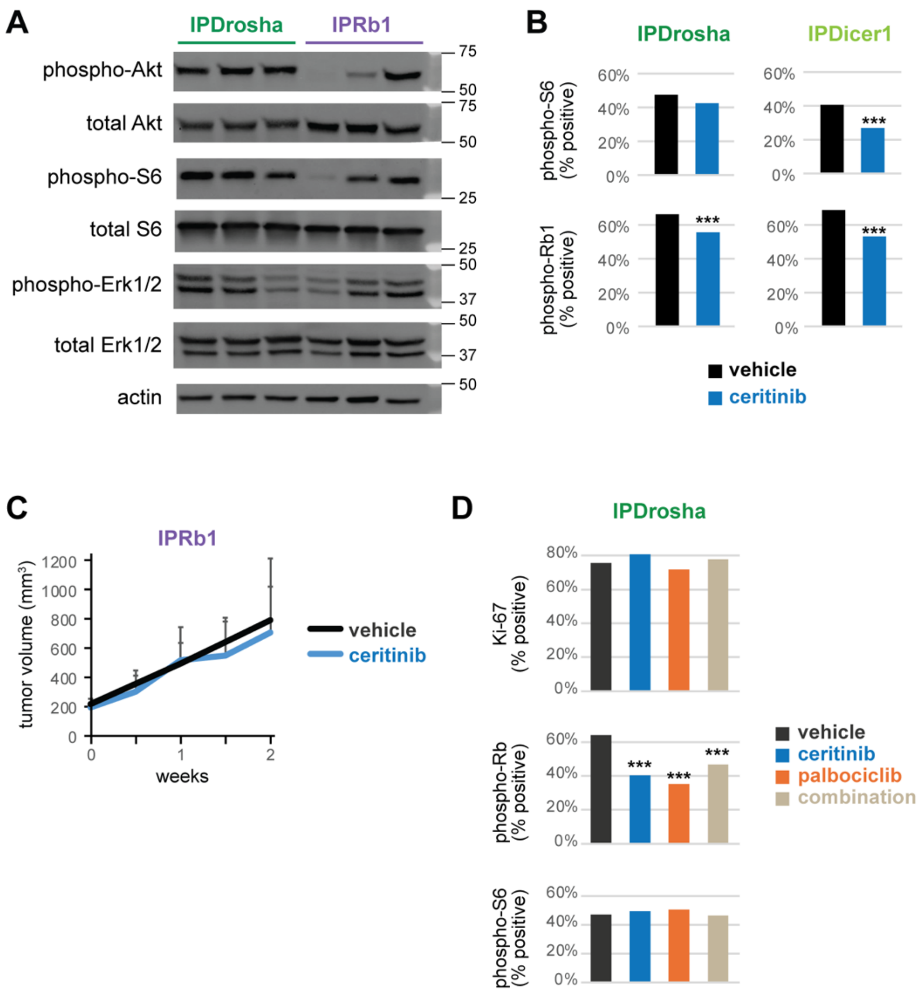

Fig S9. *DROSHA* expression is negatively correlated with let-7/miR-98-5p targets and E2F targets in human tumors.

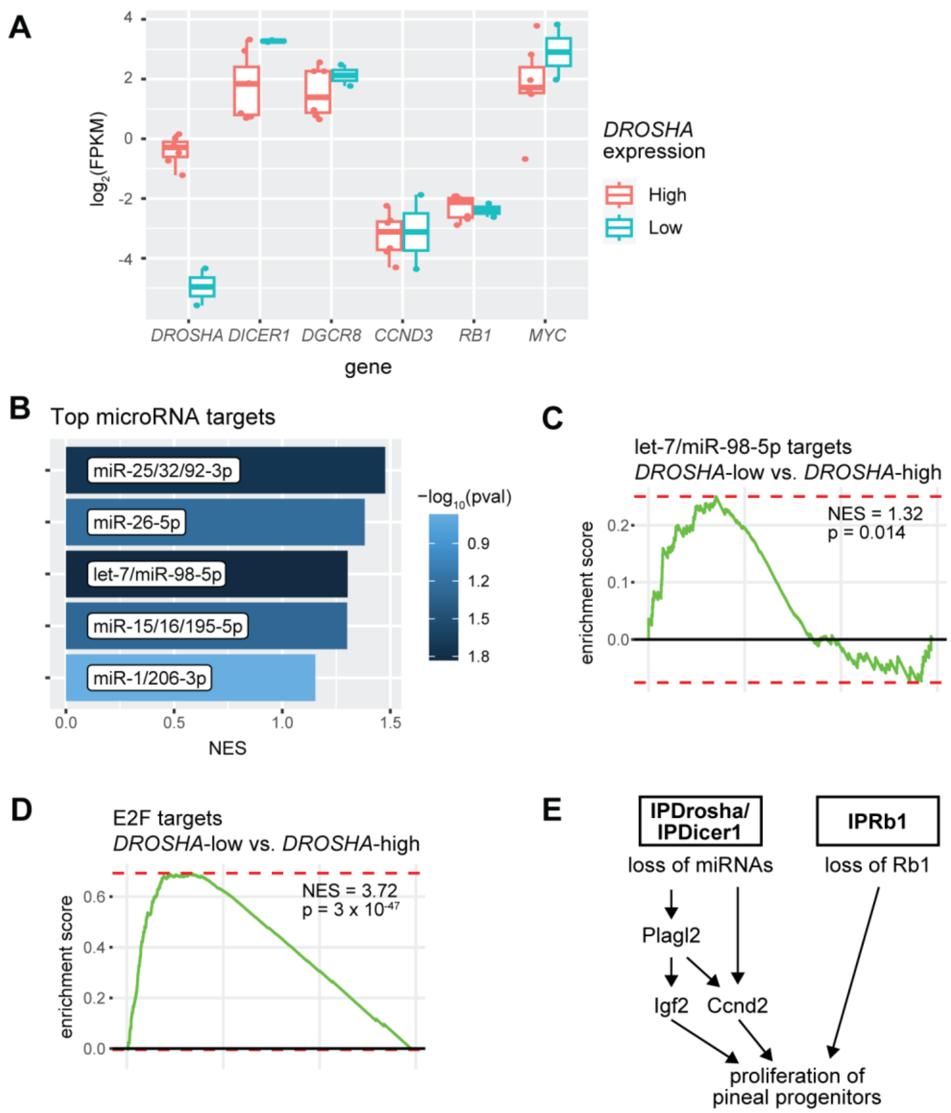
