## Supplementary figures and images for "An imbalance between proliferation and differentiation underlies the development of microRNA-defective pineoblastoma"

### Dicer GFAP tumor 20x.jpeg

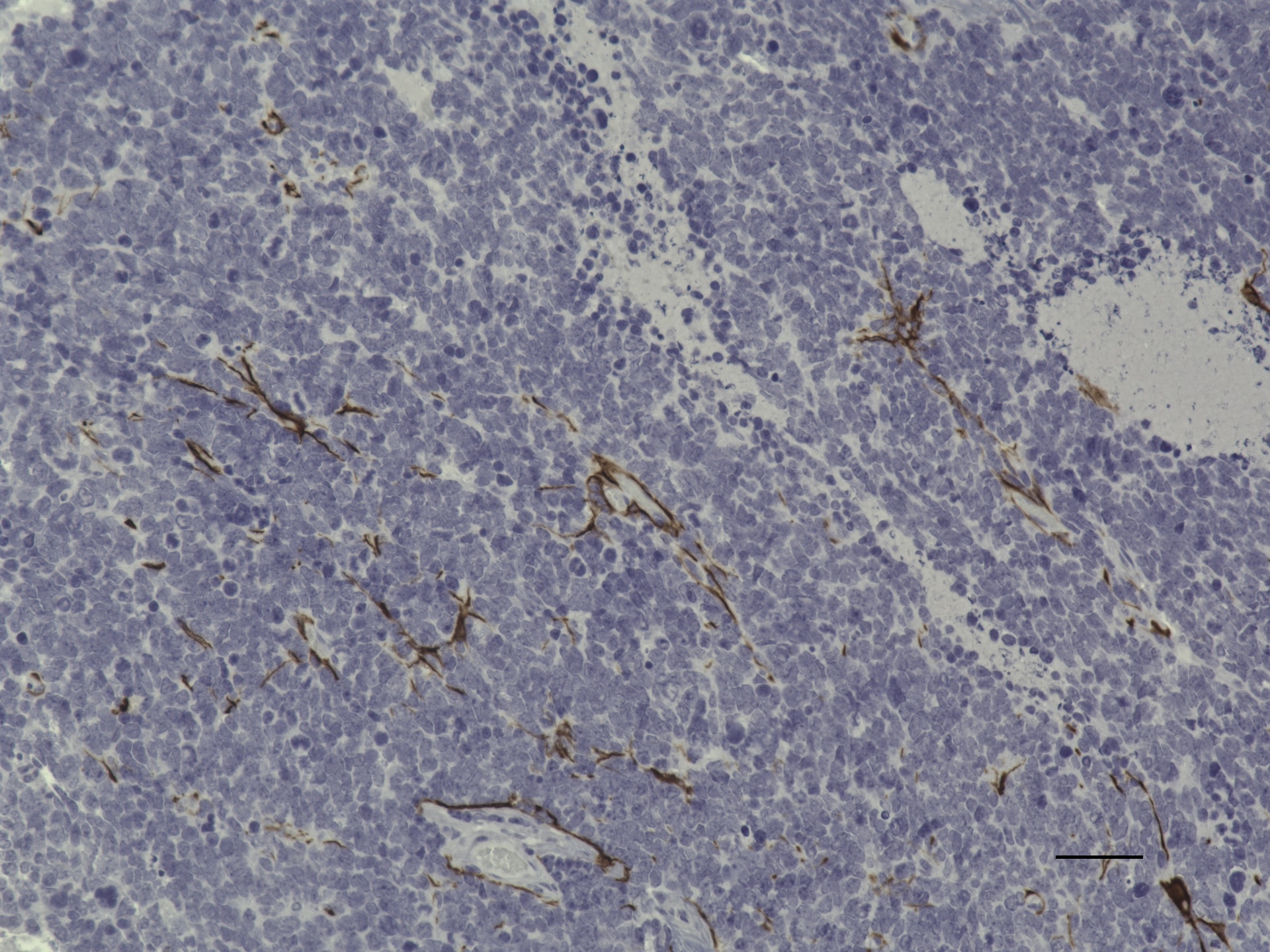

### IPDicer1 2X 1000 um.jpeg

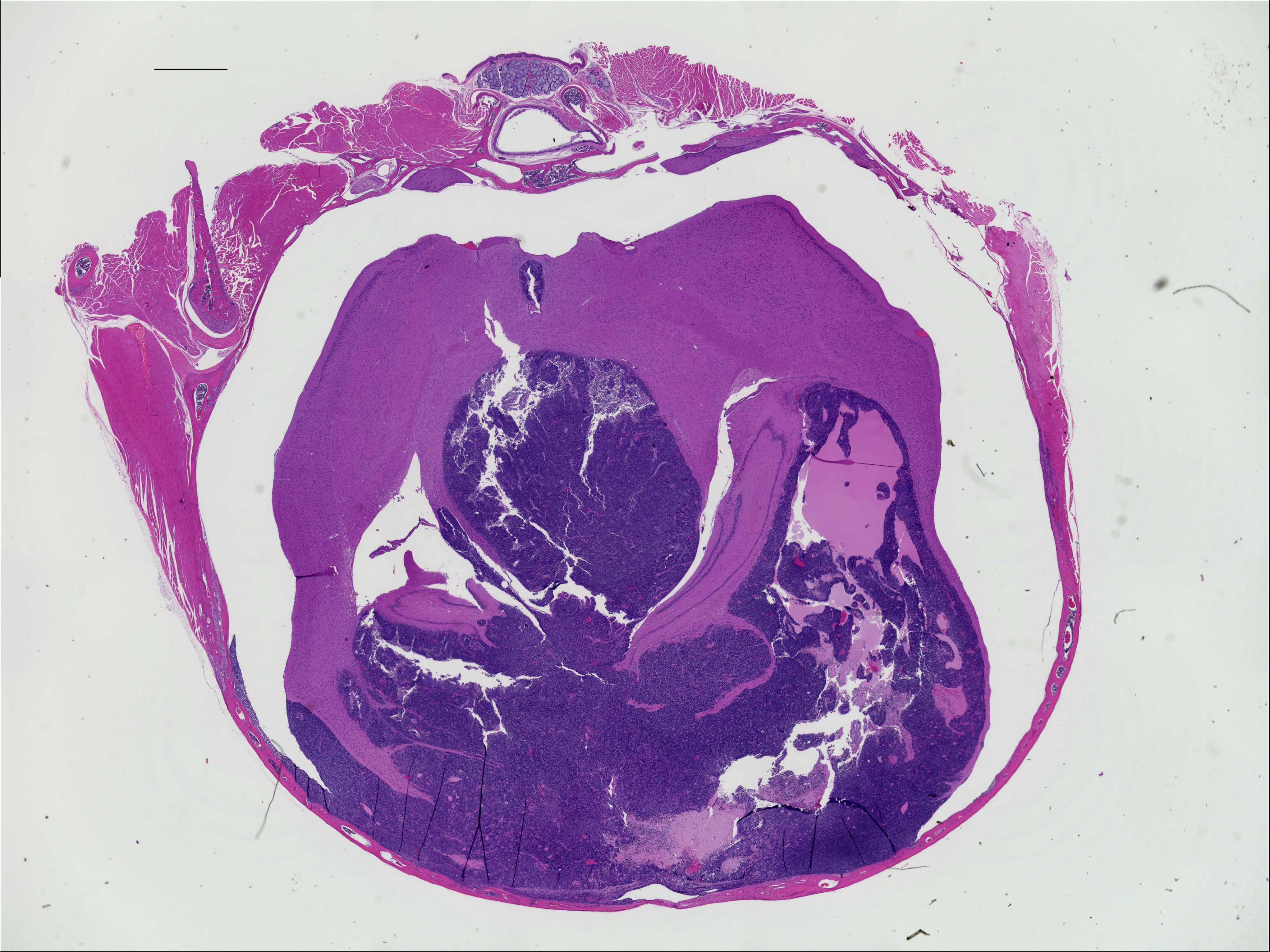

### IPDicer1 20X 50 um Ki67.jpeg

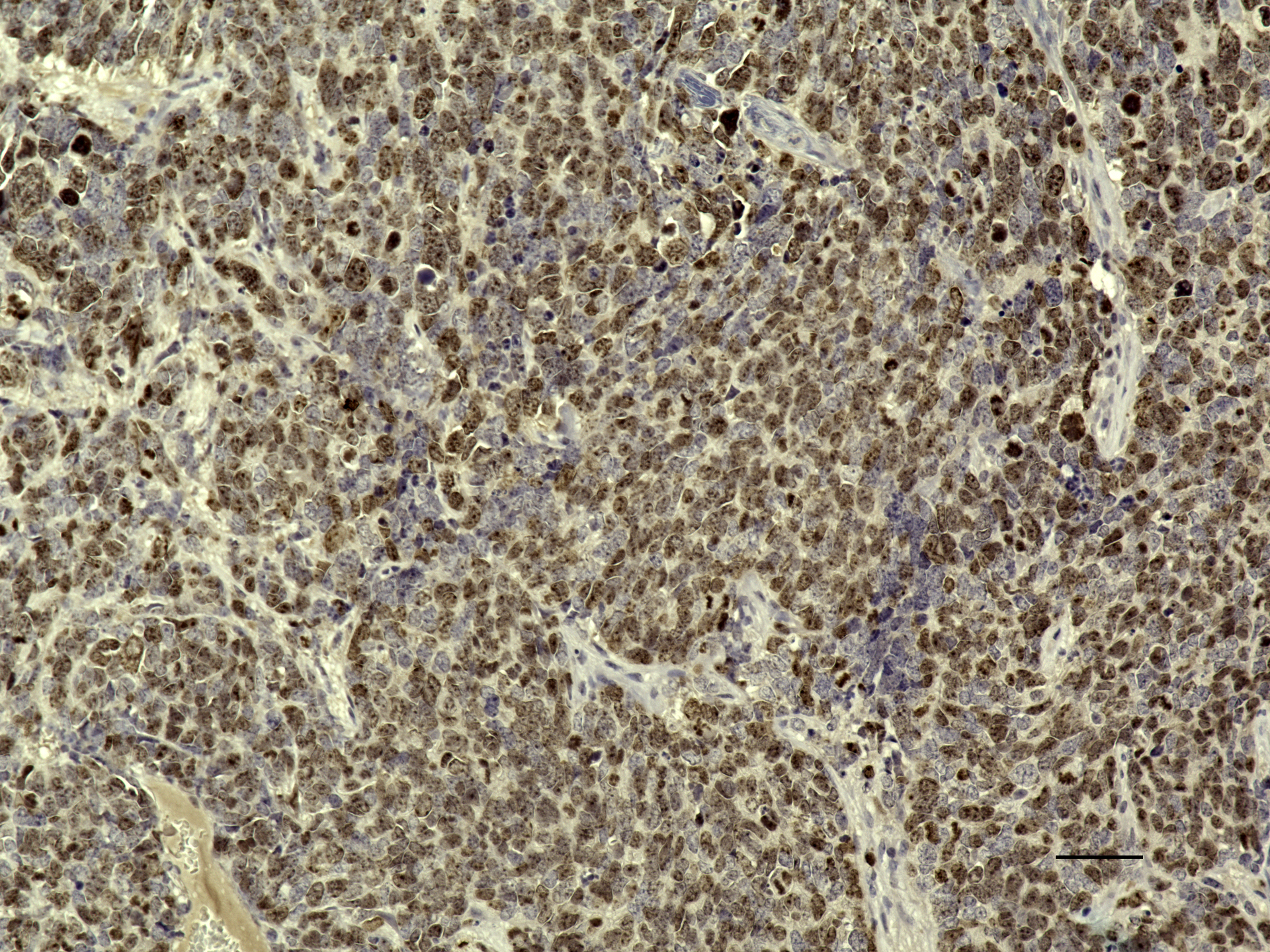

### IPDicer1 20X 50um.jpeg

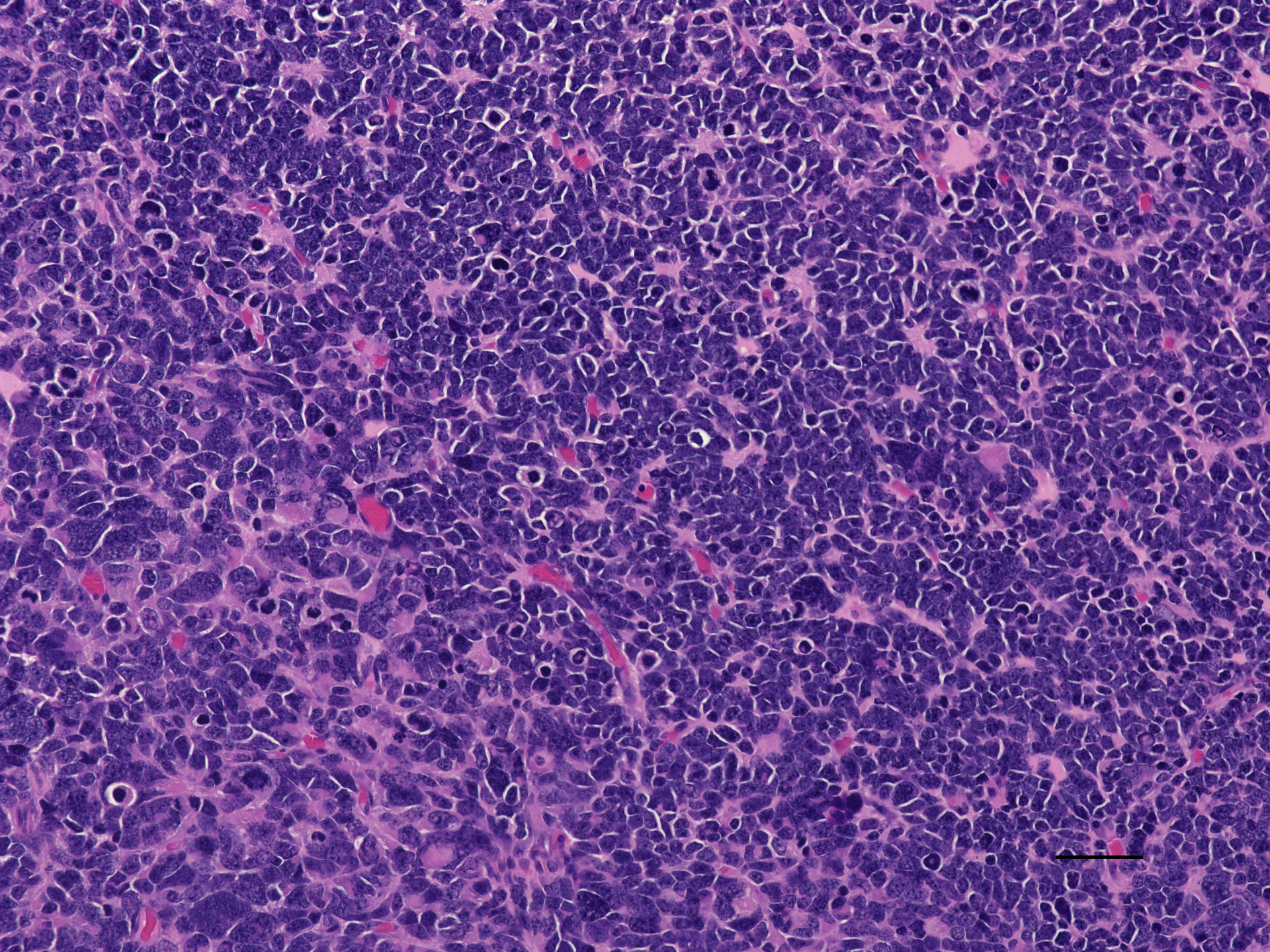

### IPDicer1 palbo 20x H_E.jpeg

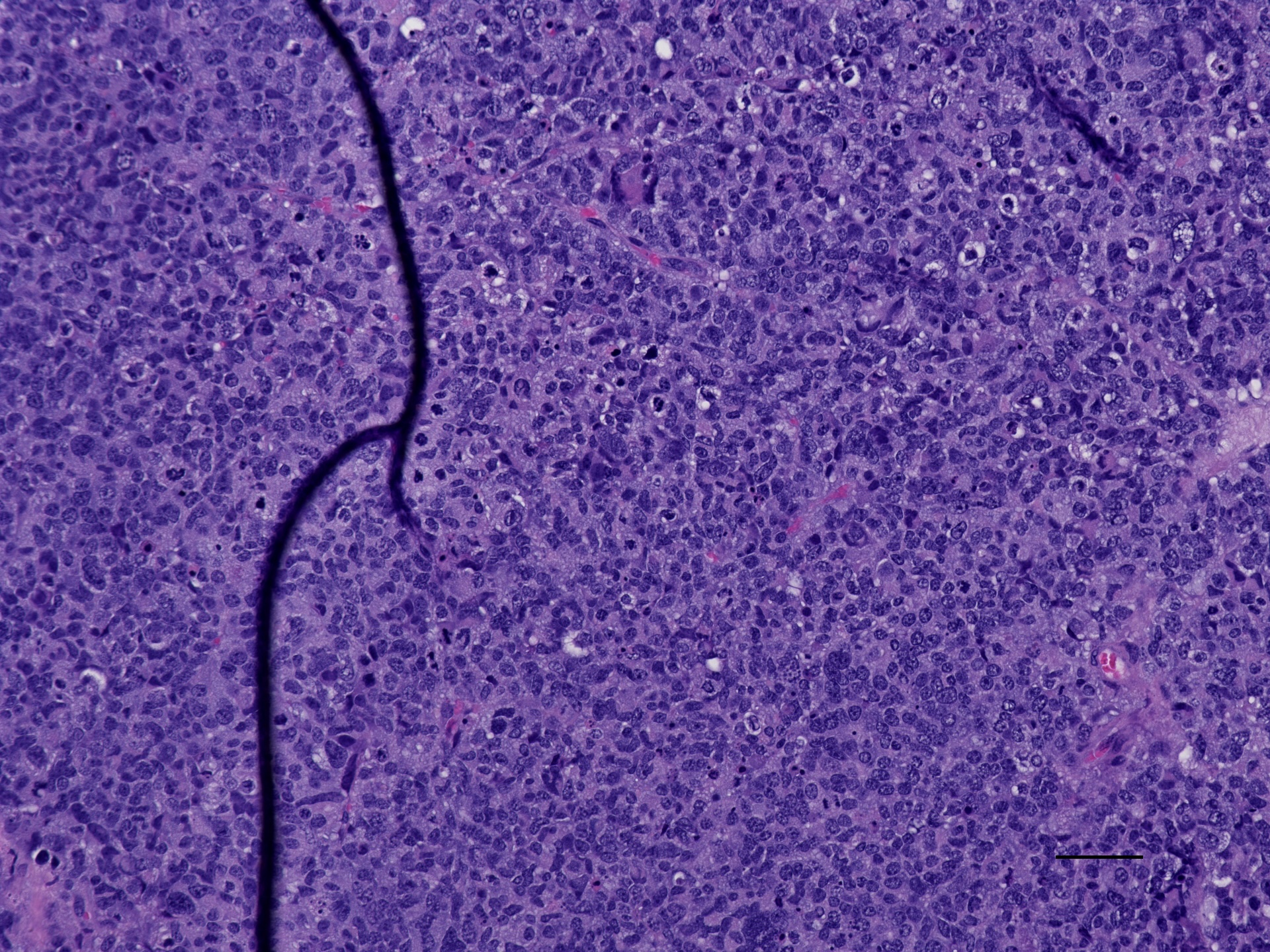

### IPDicer1 palbo 20x pRb.jpeg

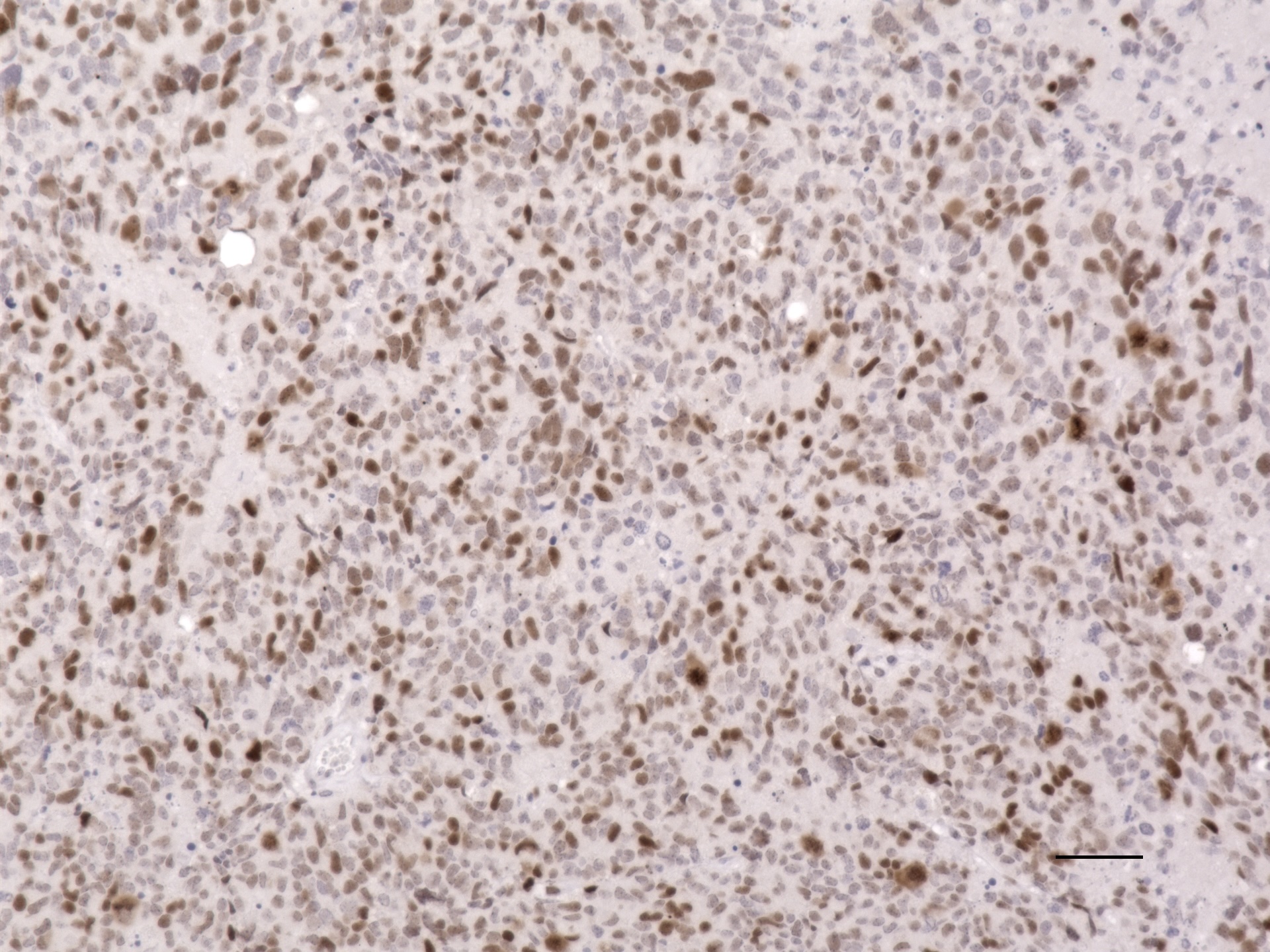

### IPDicer1 palbociclib Ki67 20x_01 50um scale.jpeg

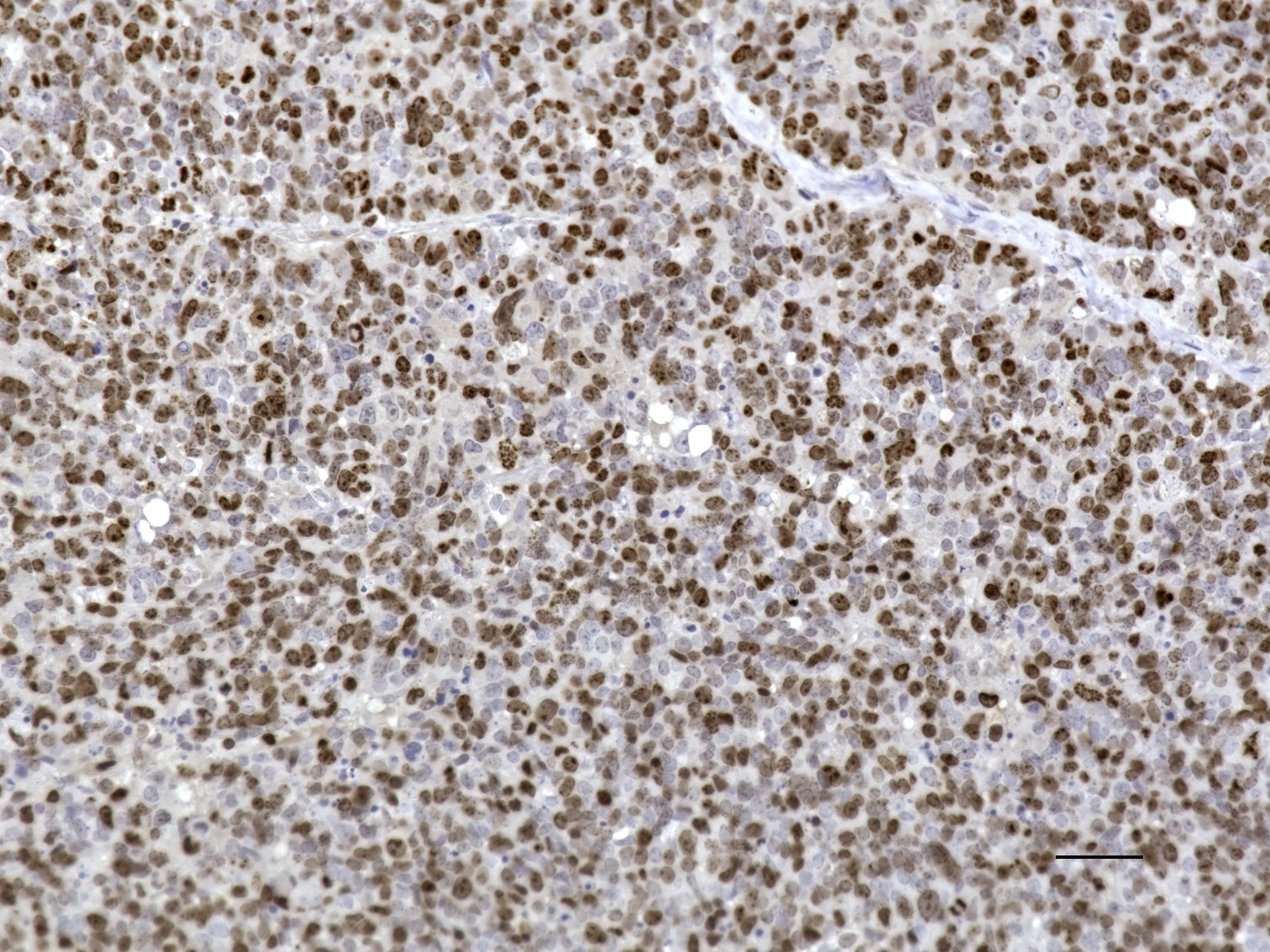

### IPDicer1 synapto 20x 50 um.jpeg

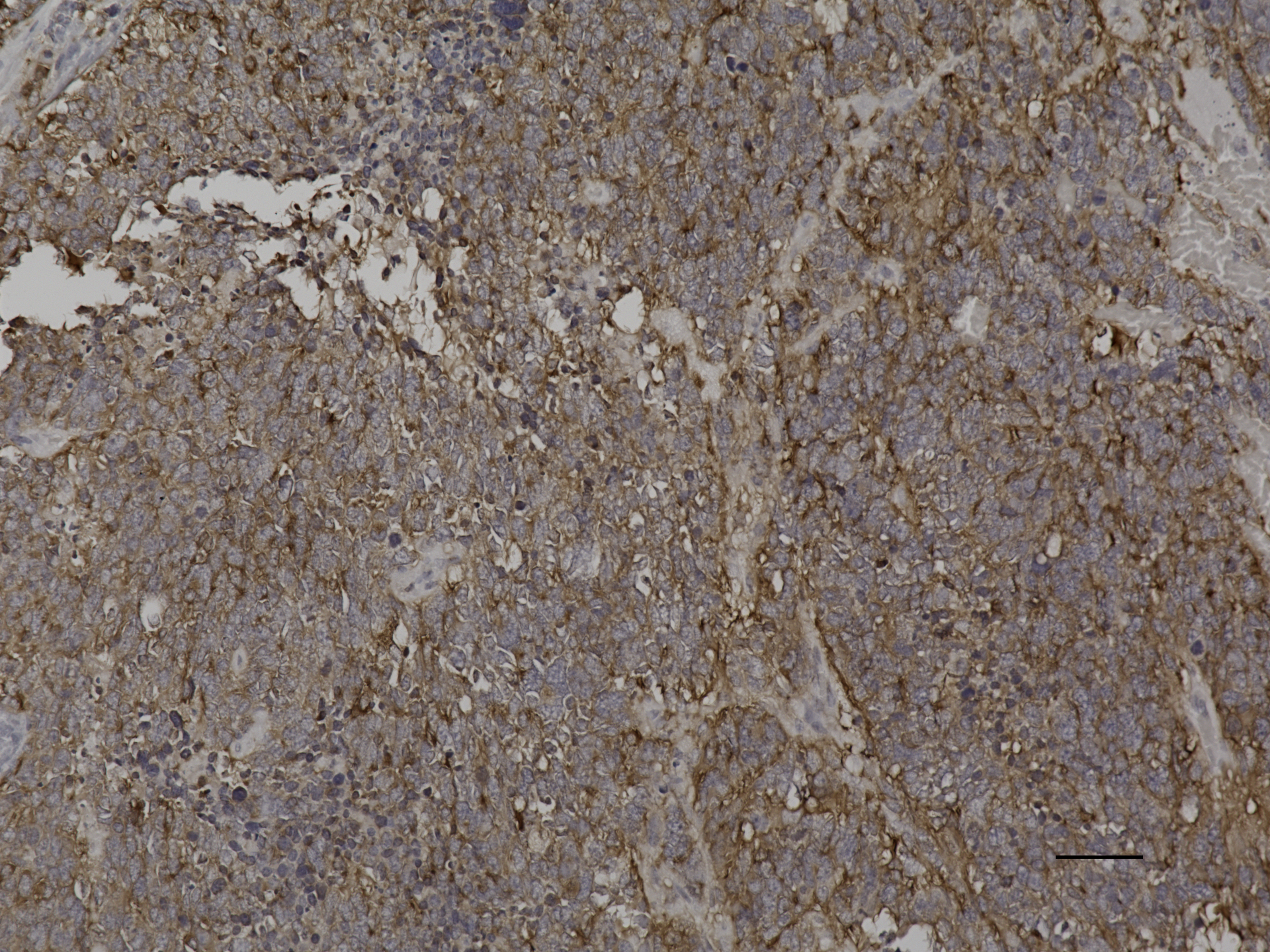

### IPDicer1 vehicle 20x H_E.jpeg

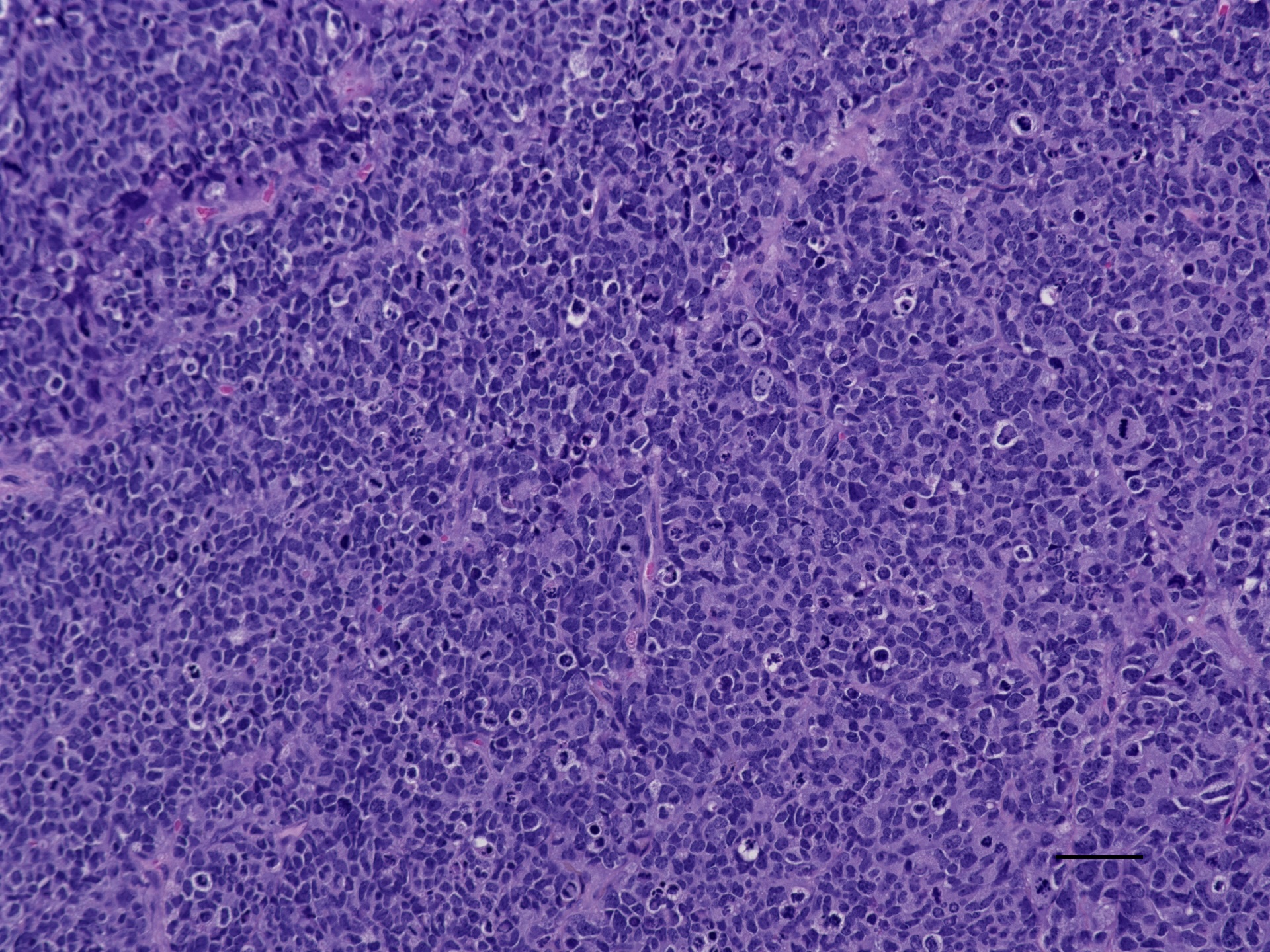

### IPDicer1 vehicle Ki67 20x_01 50 um scale.jpeg

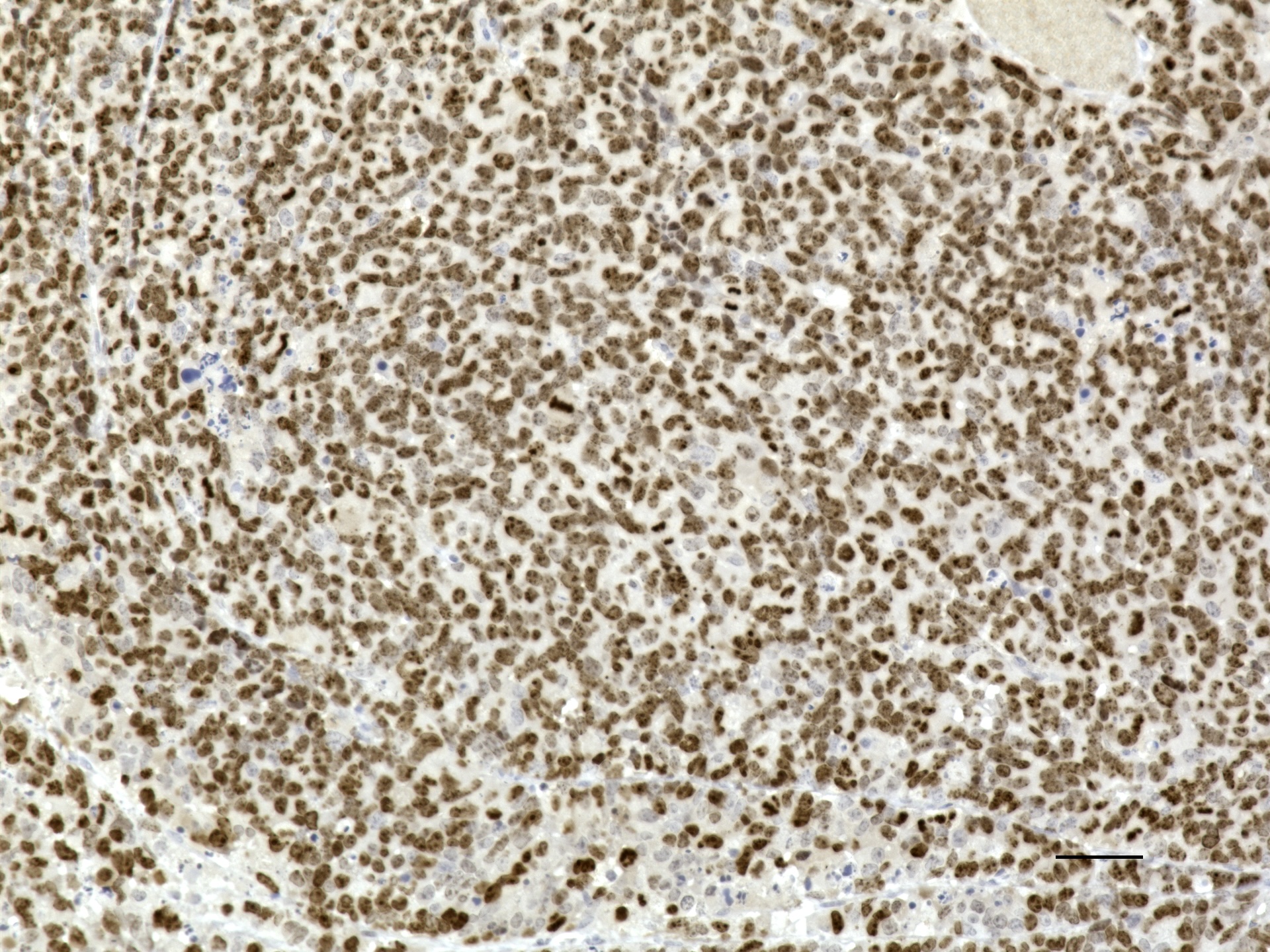

### IPDicer 20X 50 um Ki67.jpeg

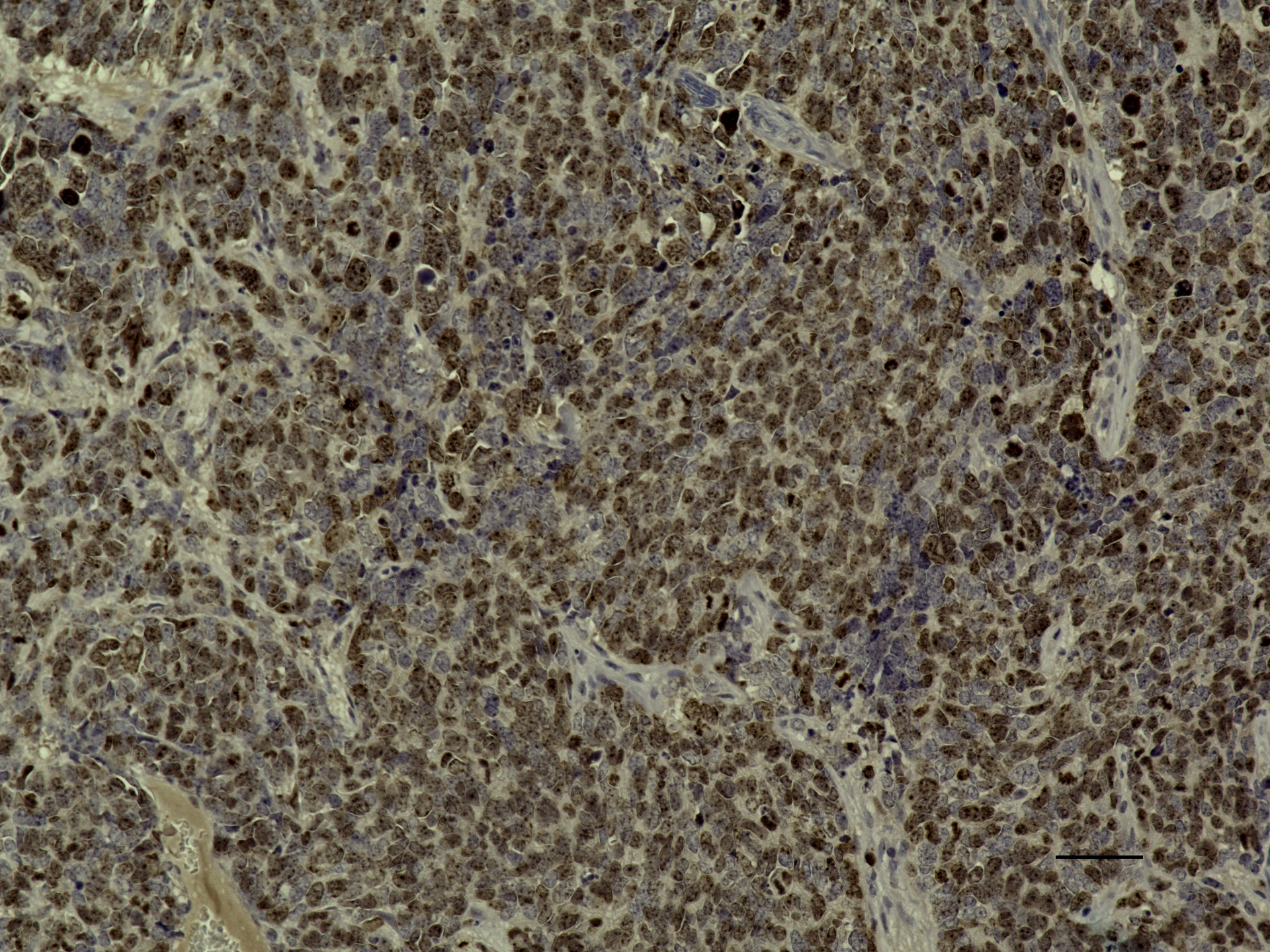

### IPDicer cerit H_E 20X.jpeg

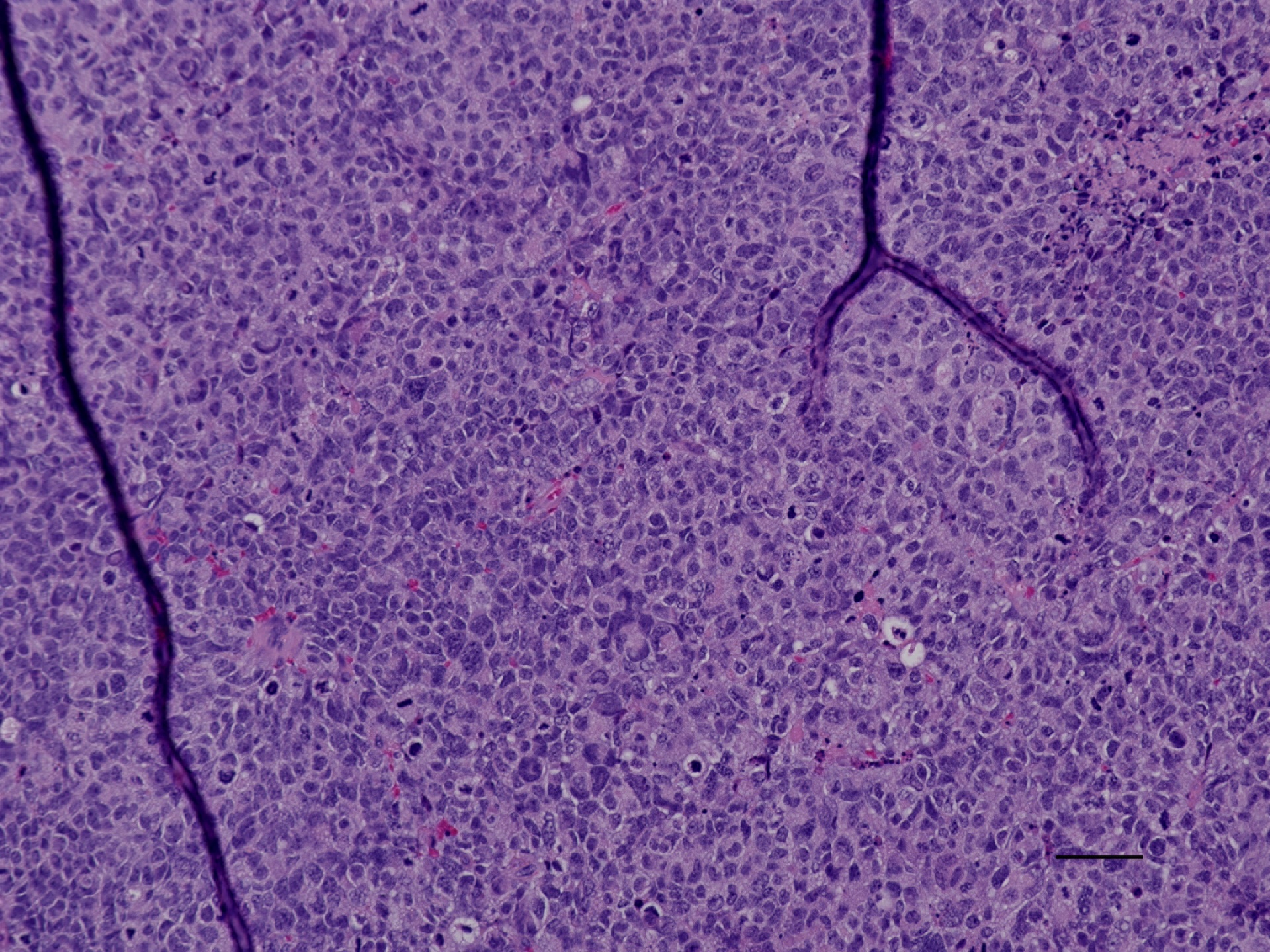

### IPDicer cerit pRb 20x.jpeg

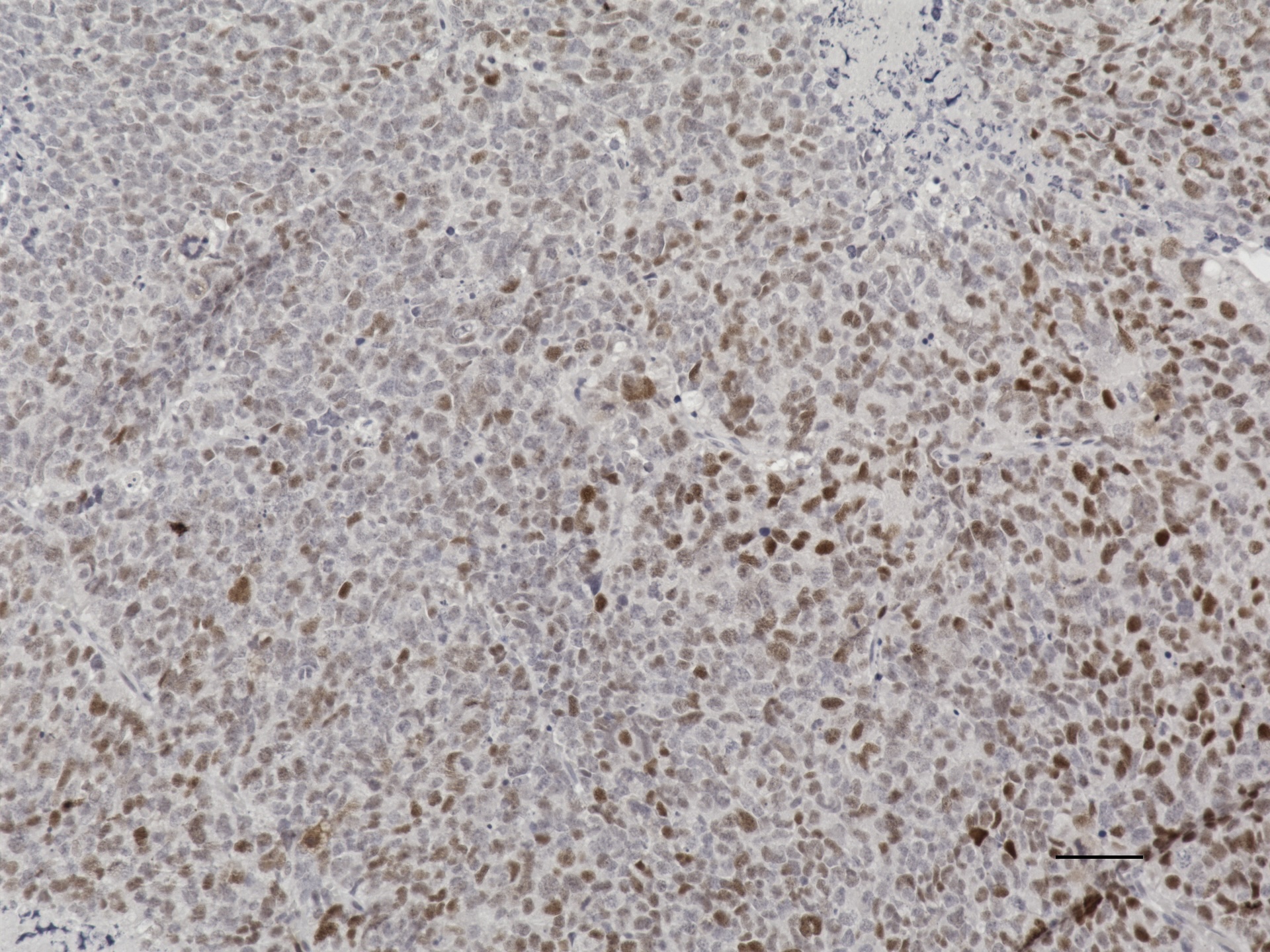

### IPDicer cerit pS6 20X 50um.jpeg

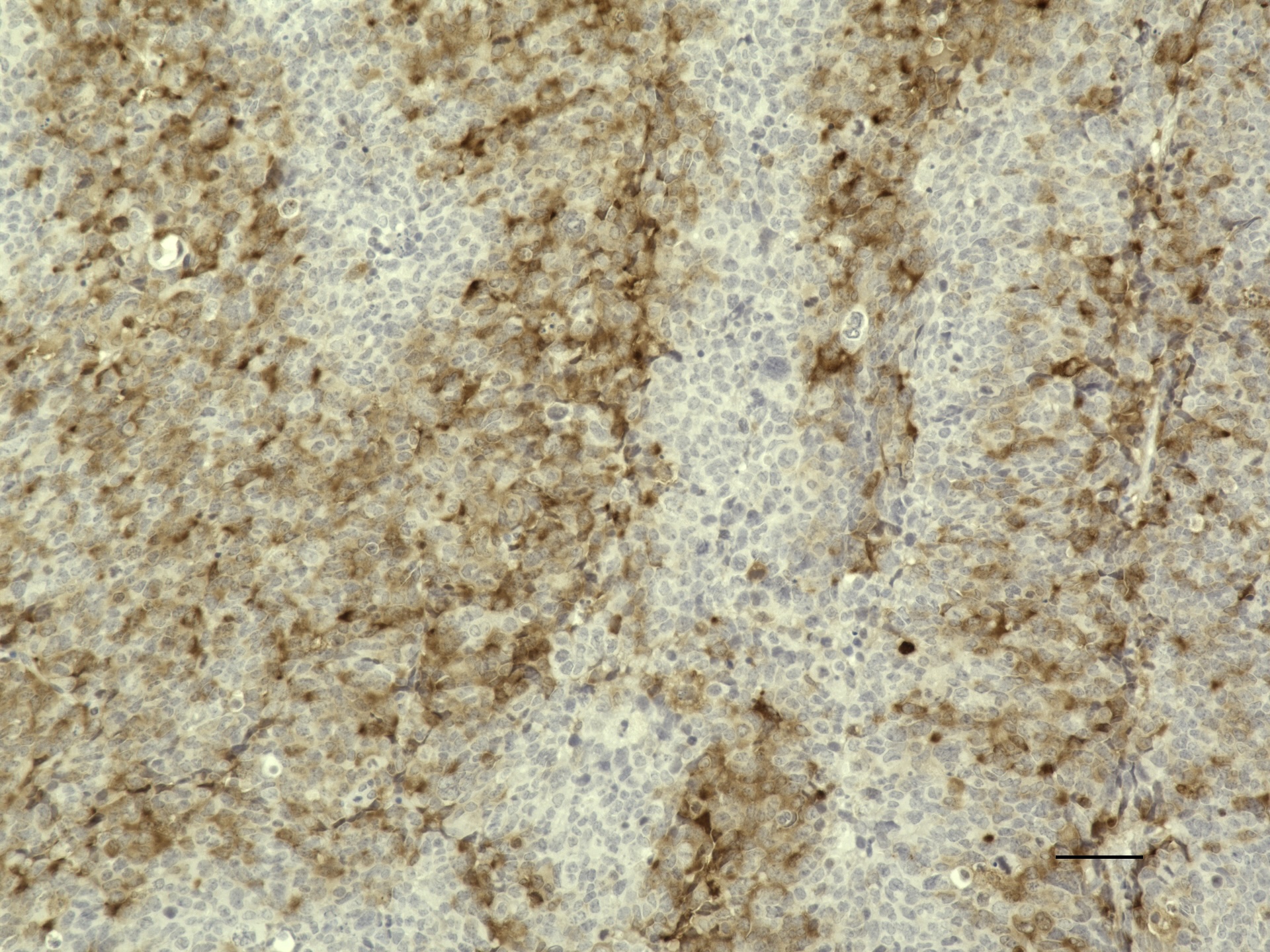

### IPDicer pRb_20X.jpg

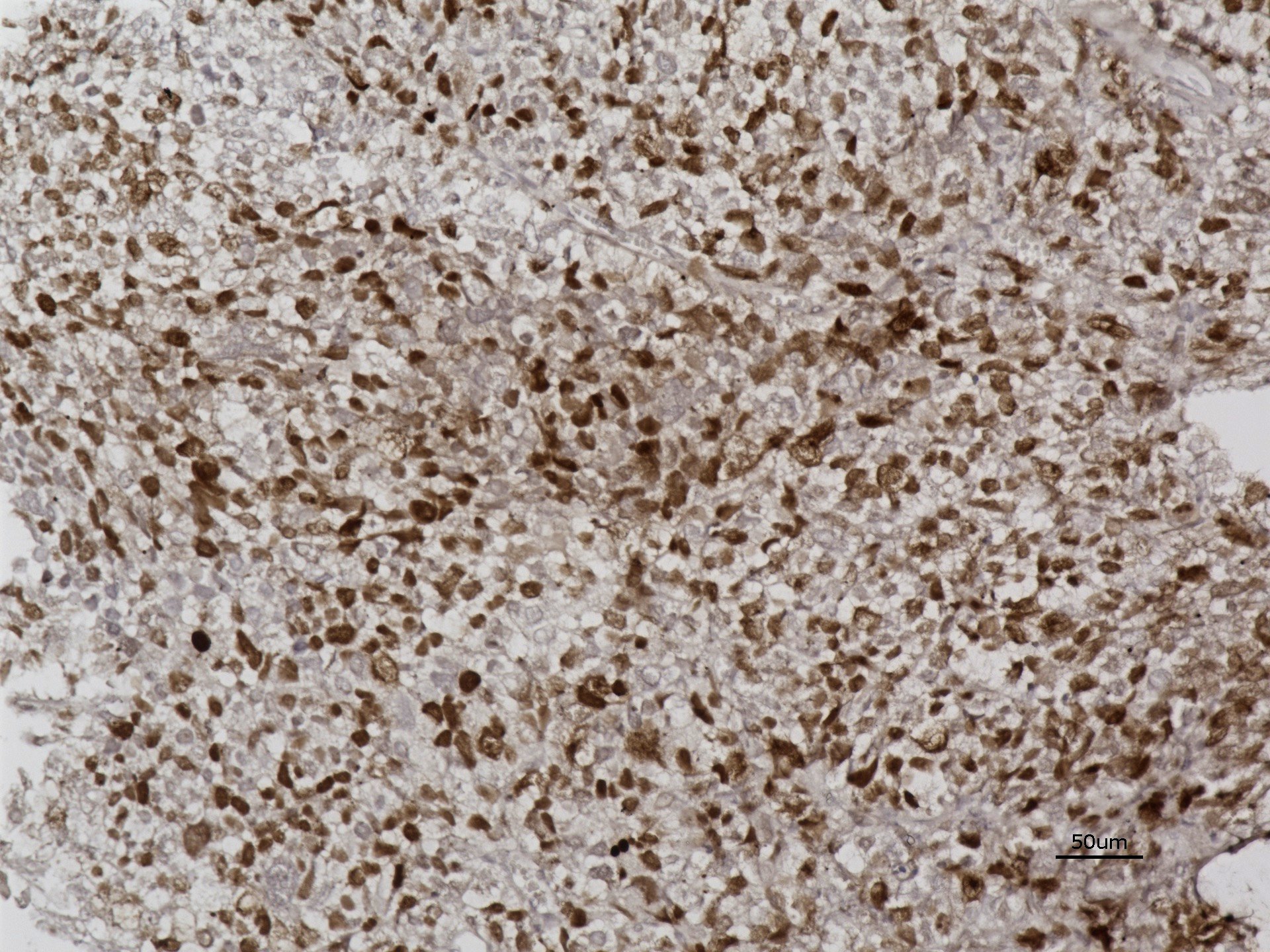

### IPDicer vehicle 20x pRb.jpeg

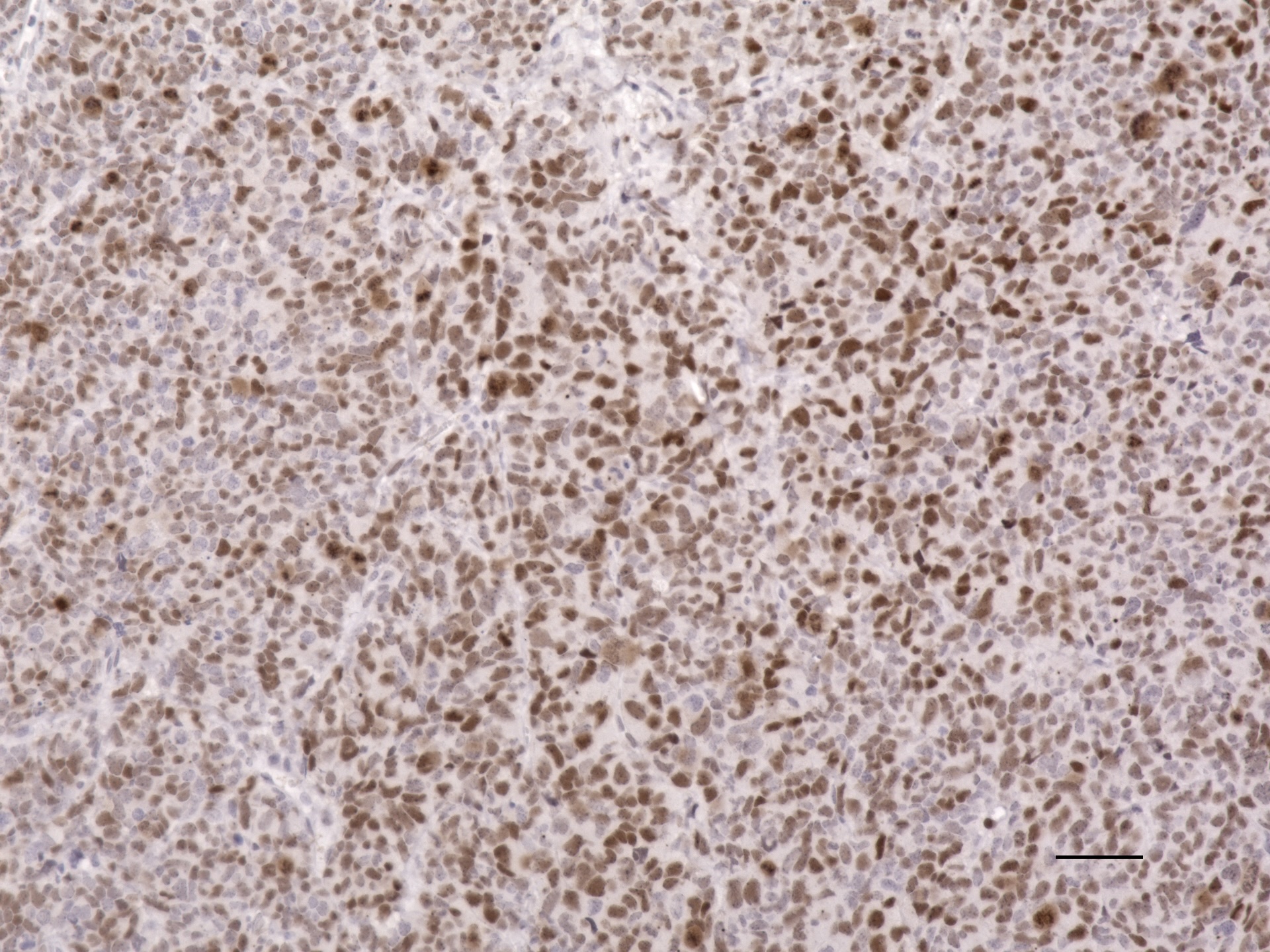

### IPDicer vehicle H_E 20x.jpeg

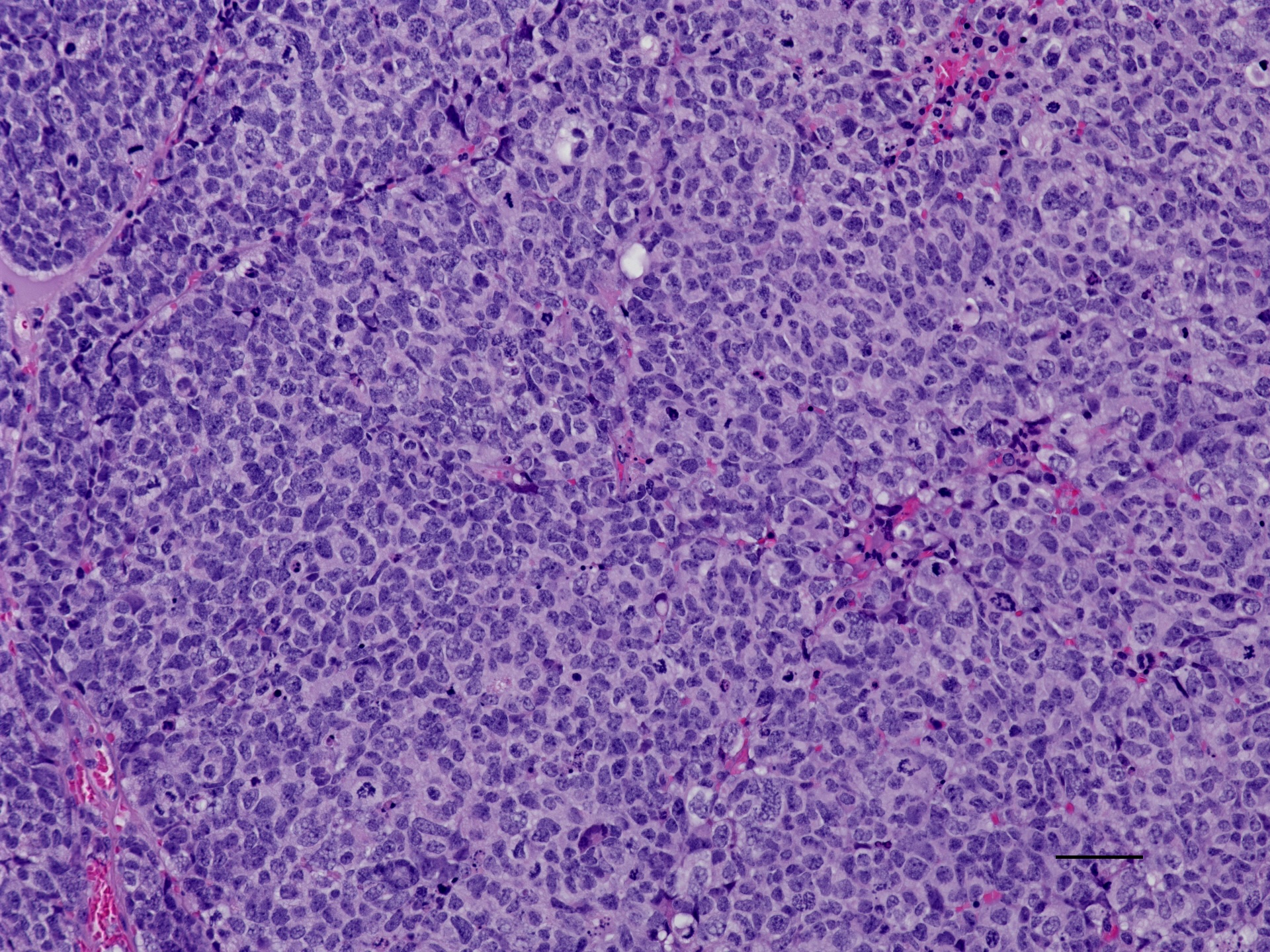

### IPDicer vehicle pRb 20x.jpeg

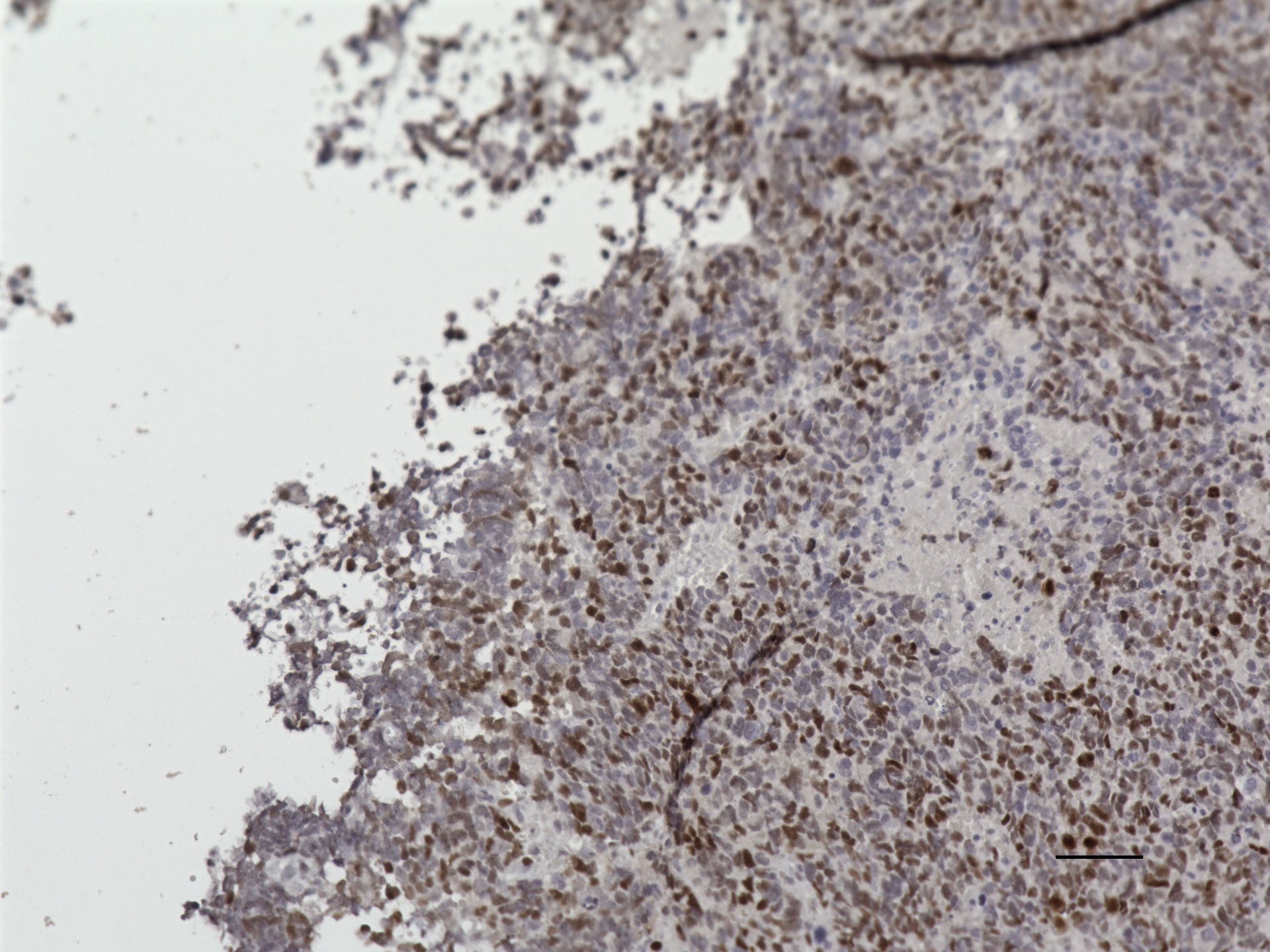

### IPDicer vehicle pS6 20X 50um.jpeg

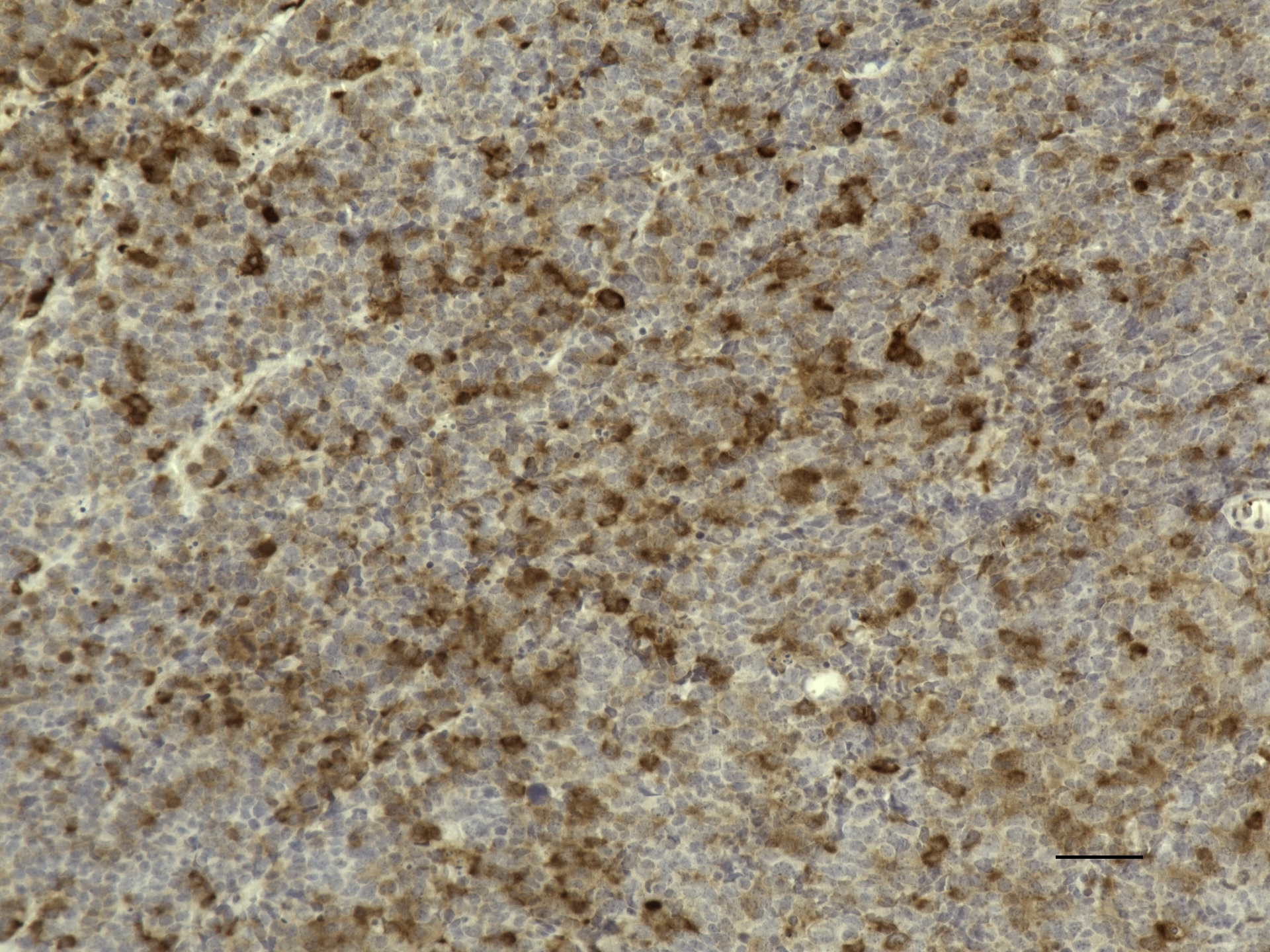

### IPDrosha 2x 1000 um.jpeg

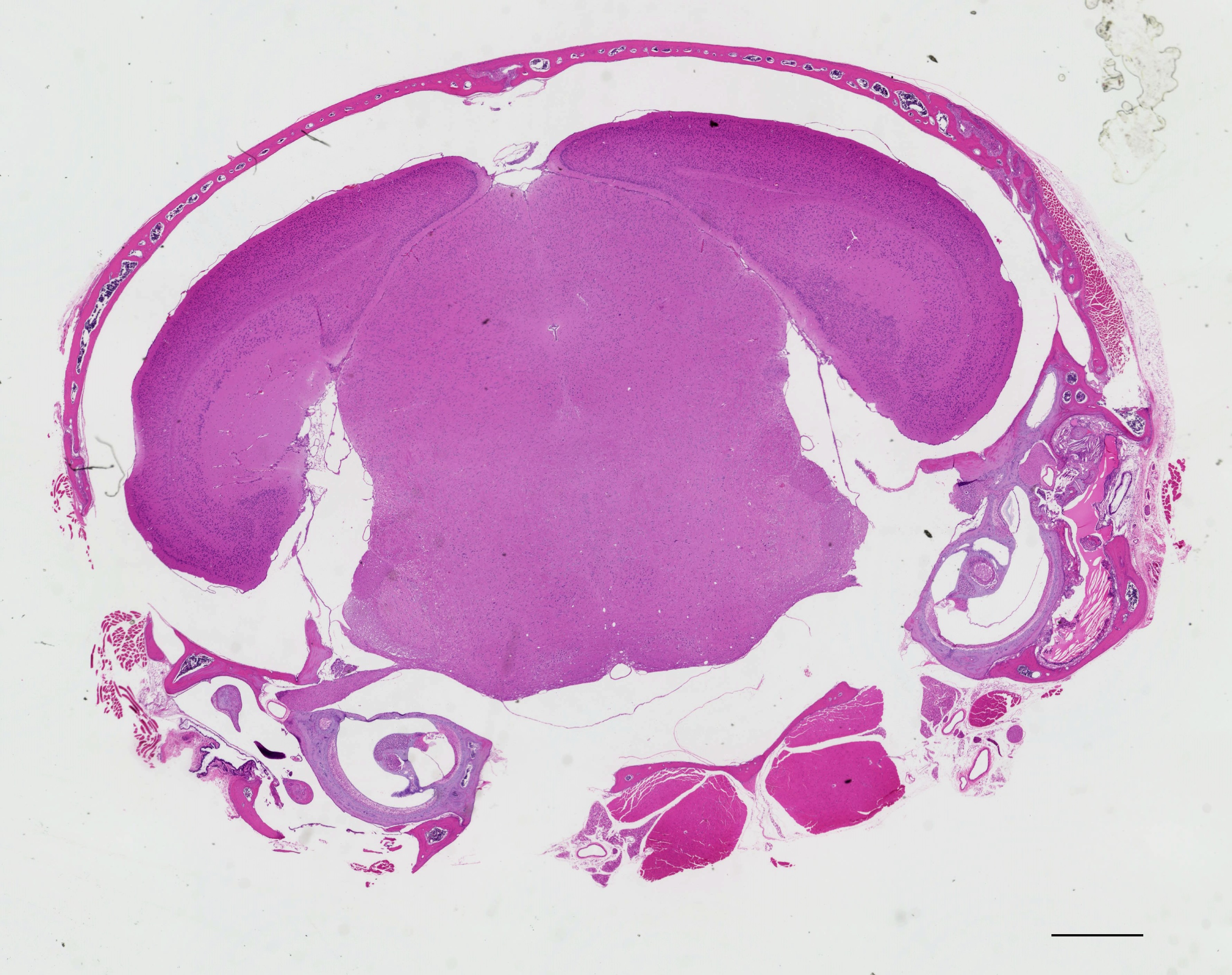

### IPDrosha 2X_1000 um.jpeg

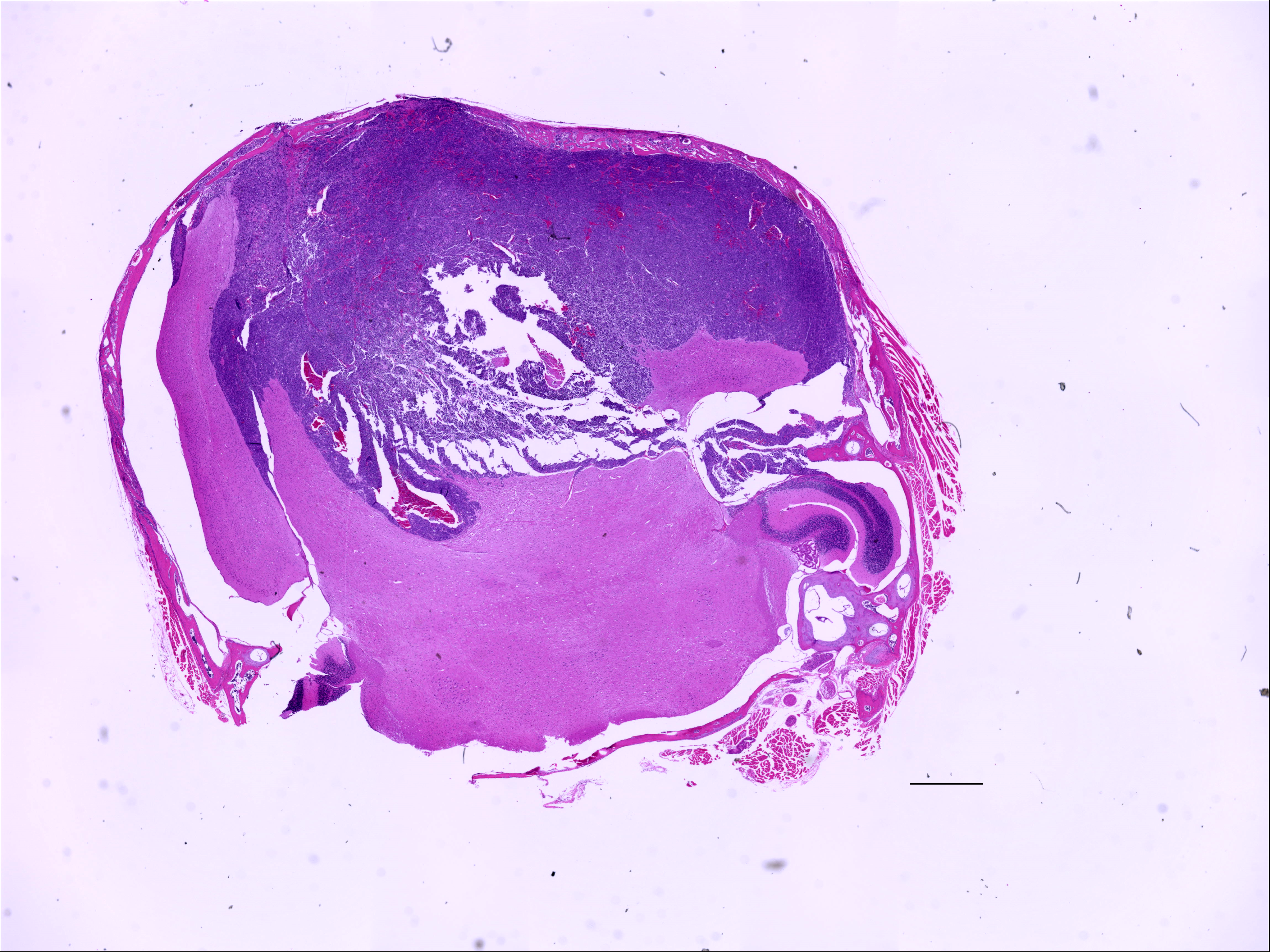

### IPDrosha 20X H_E.jpg

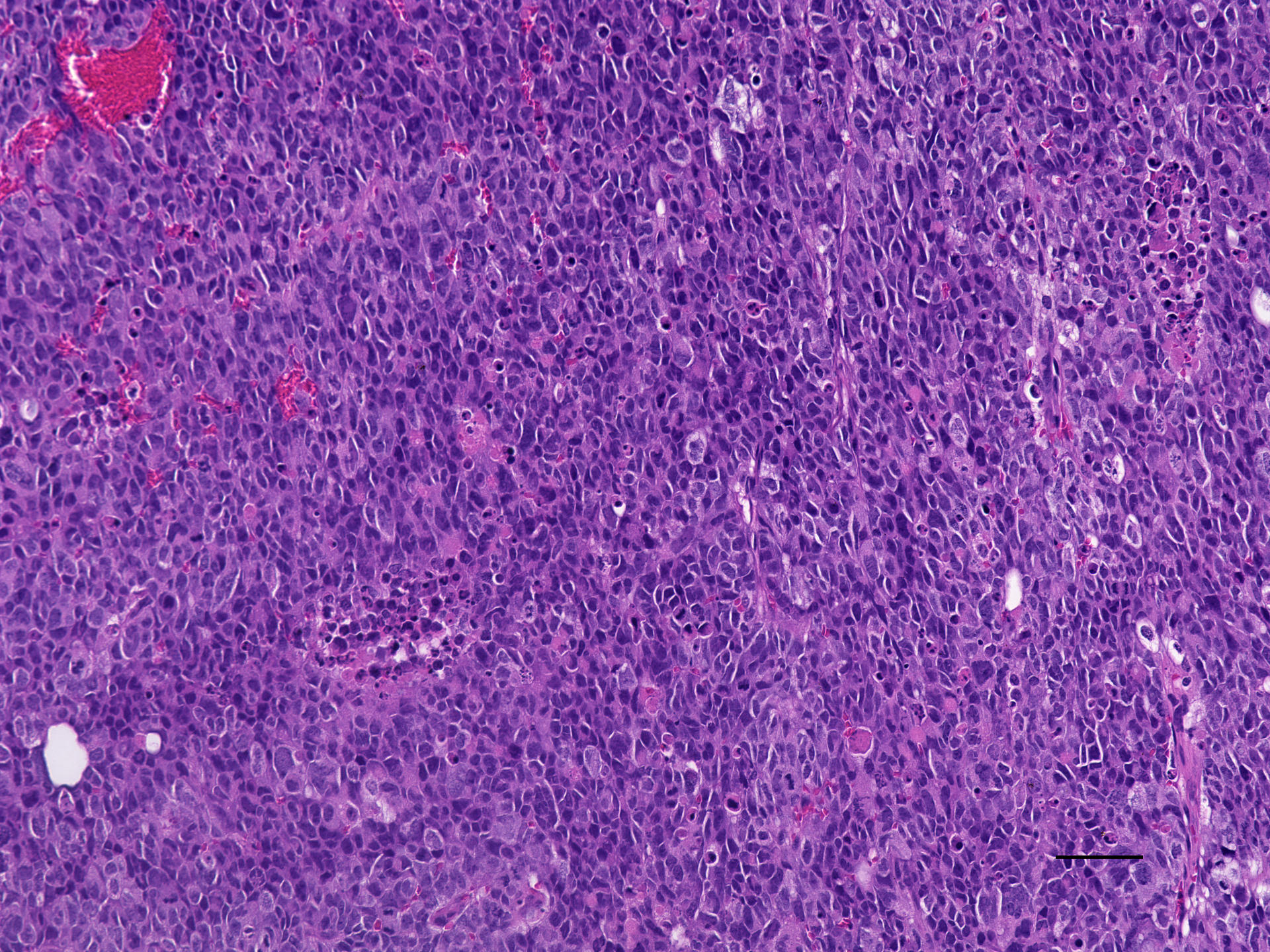

### IPDrosha cerit 20X H_E.jpeg

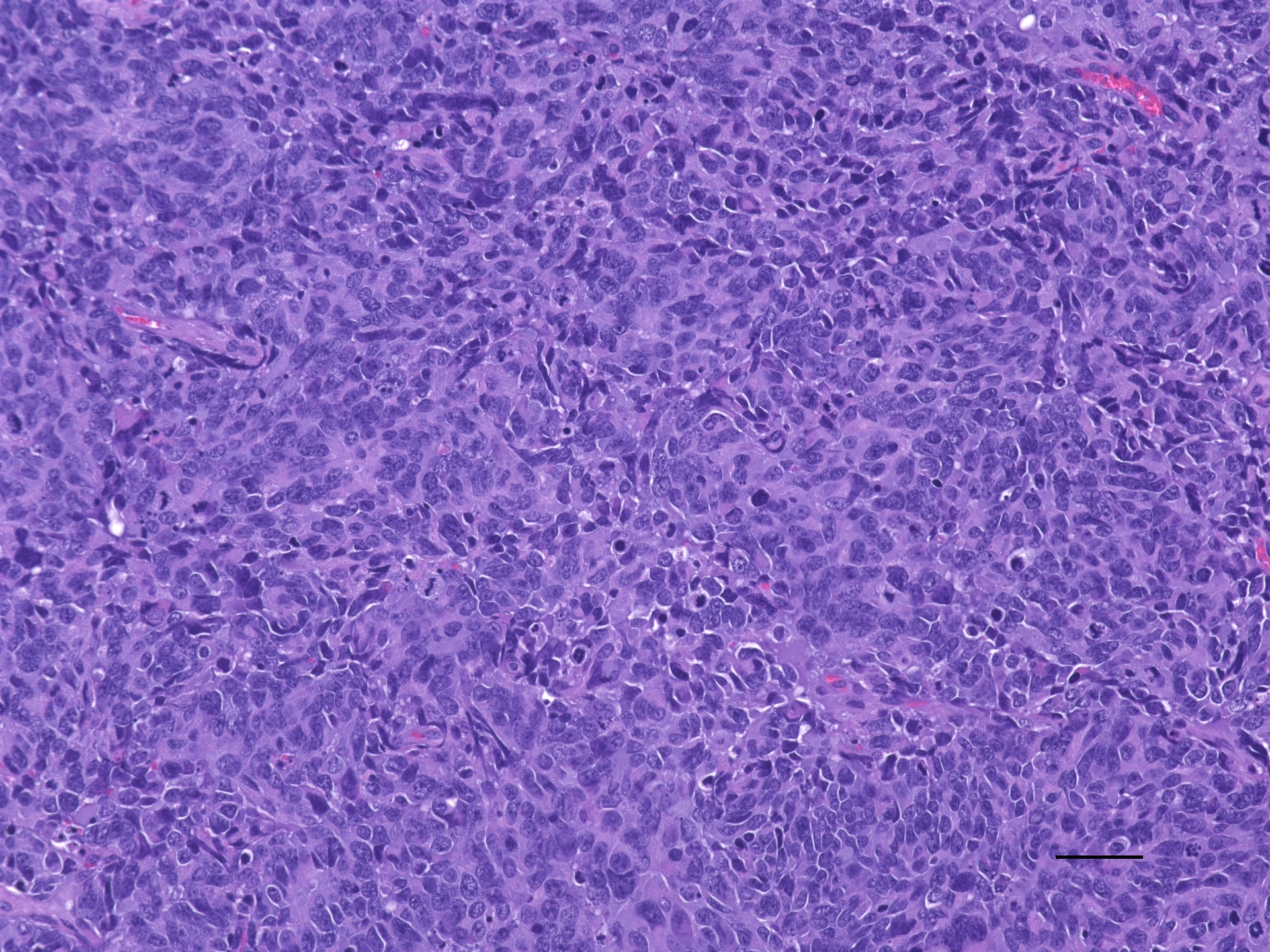

### IPDrosha cerit H_E 20X.jpeg

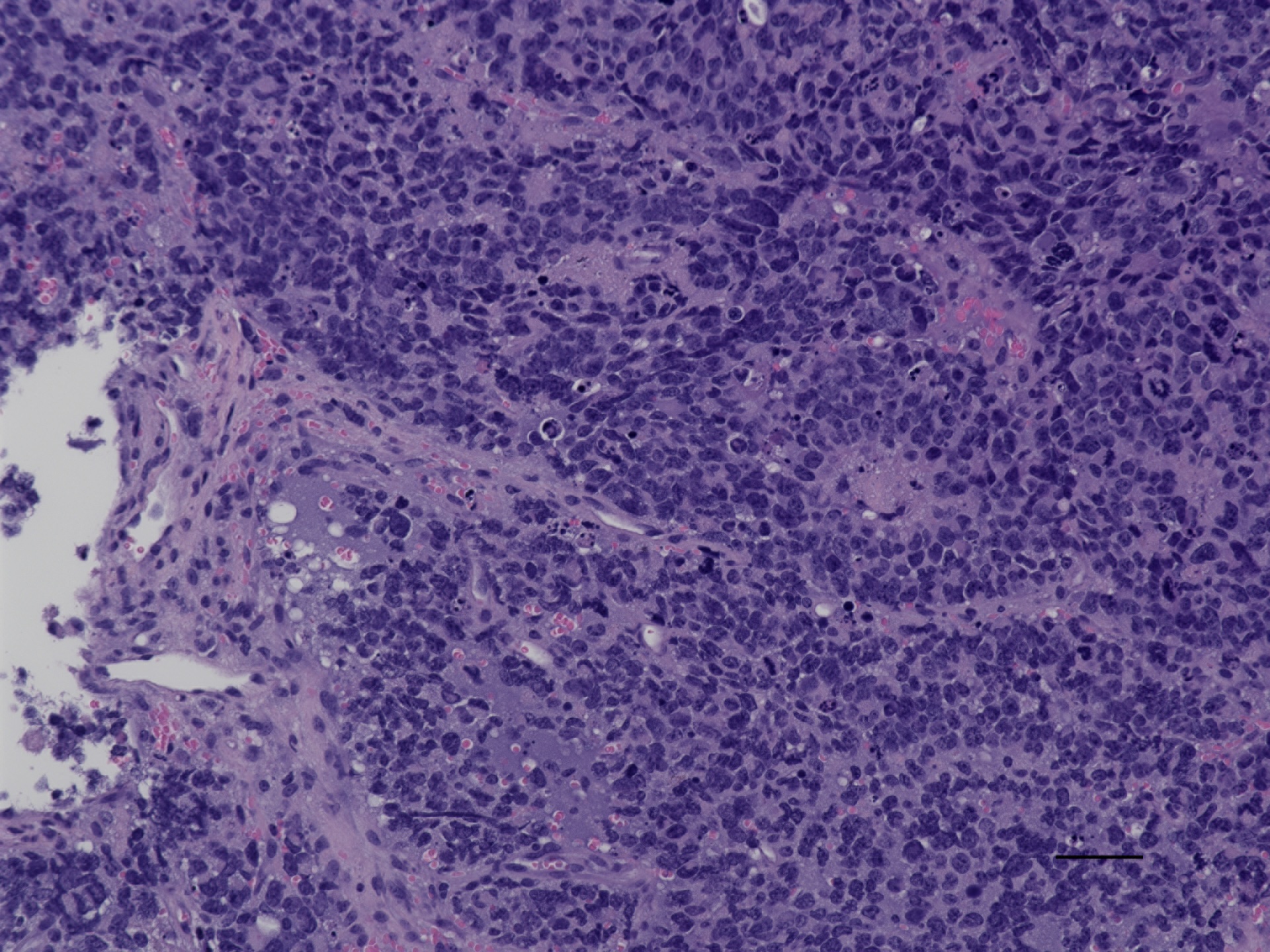

### IPDrosha cerit Ki67 20x.jpeg

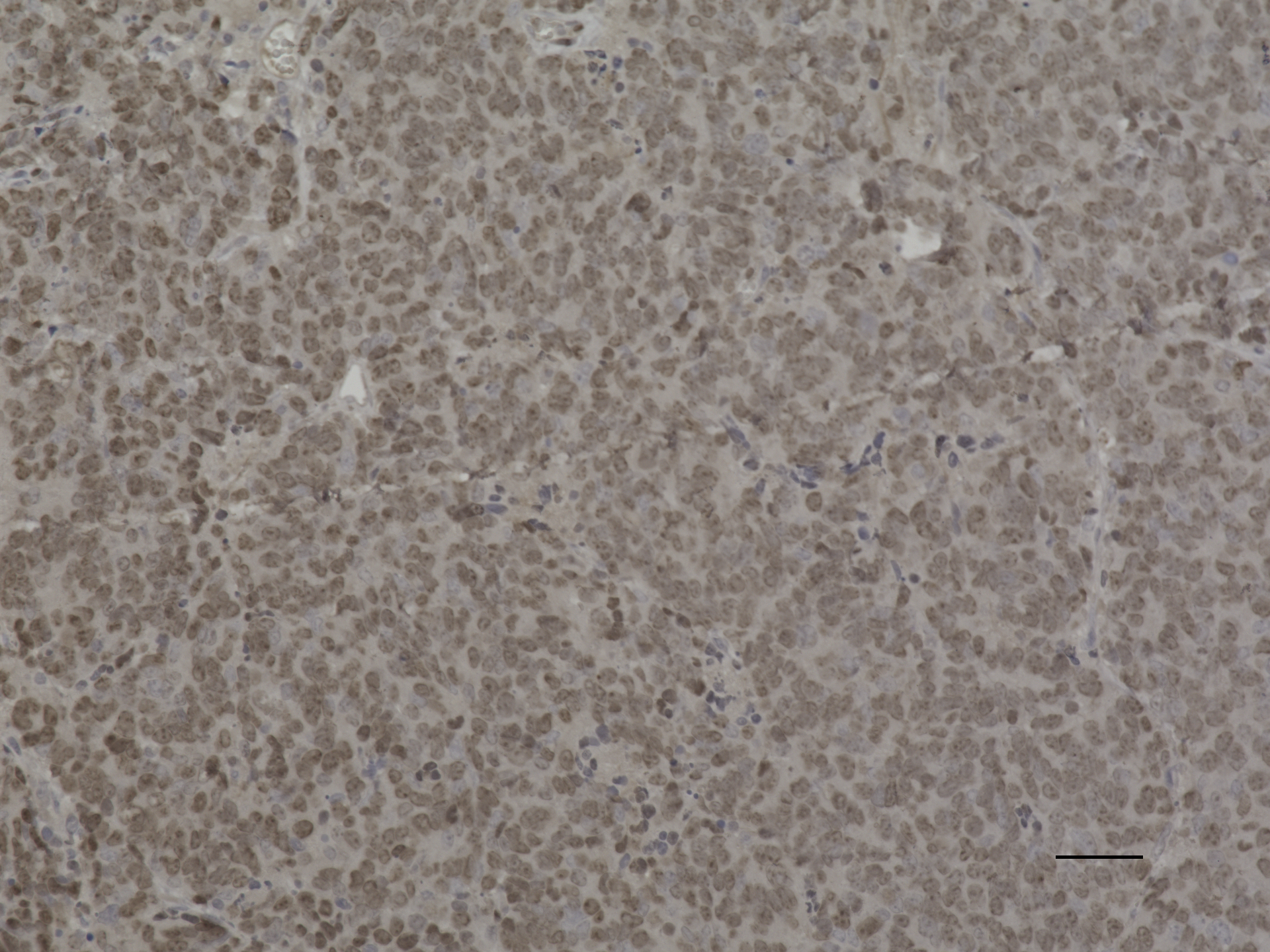

### IPDrosha cerit pRb 20X 50um.jpeg

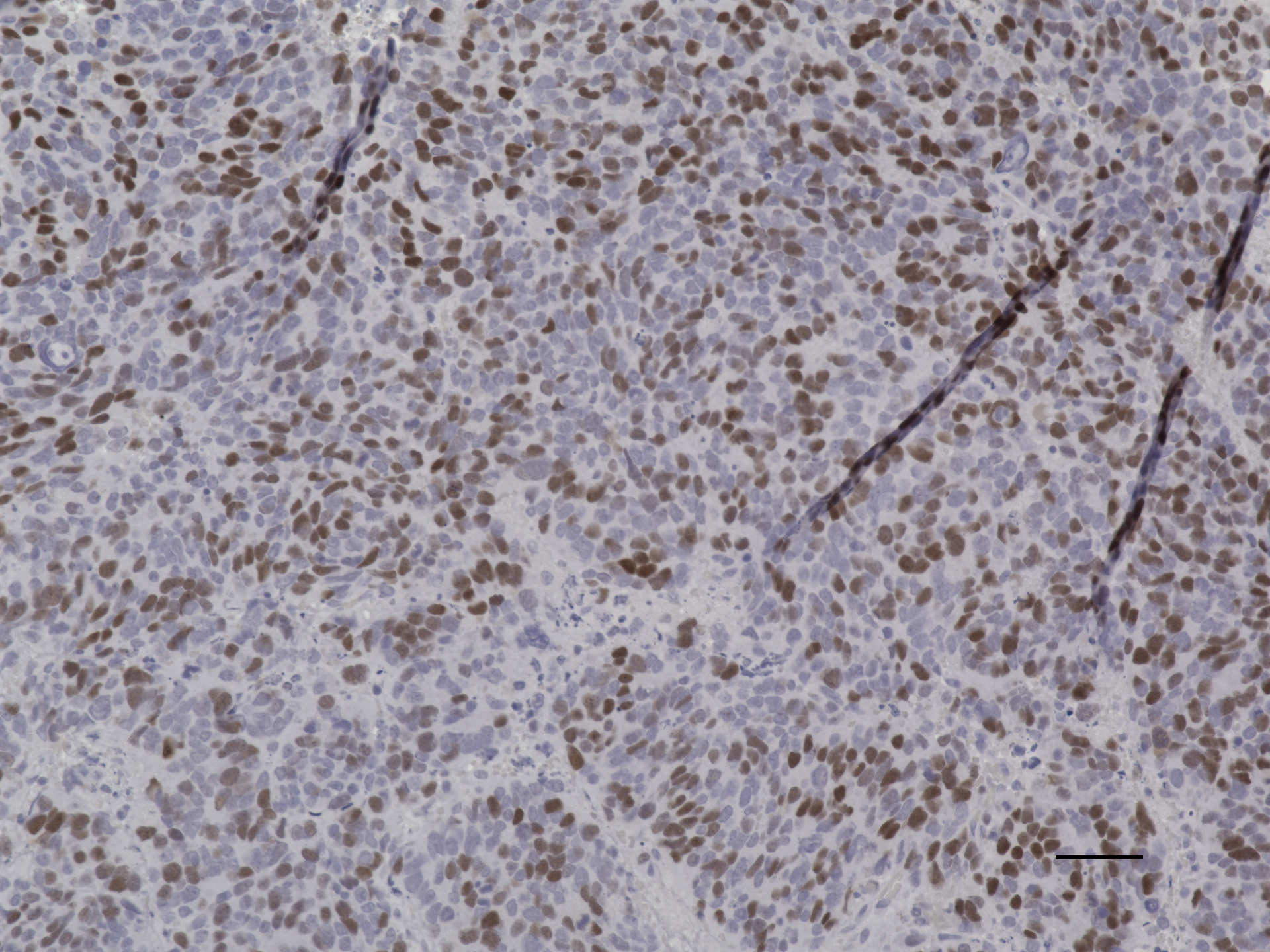

### IPDrosha cerit pRb 20x.jpeg

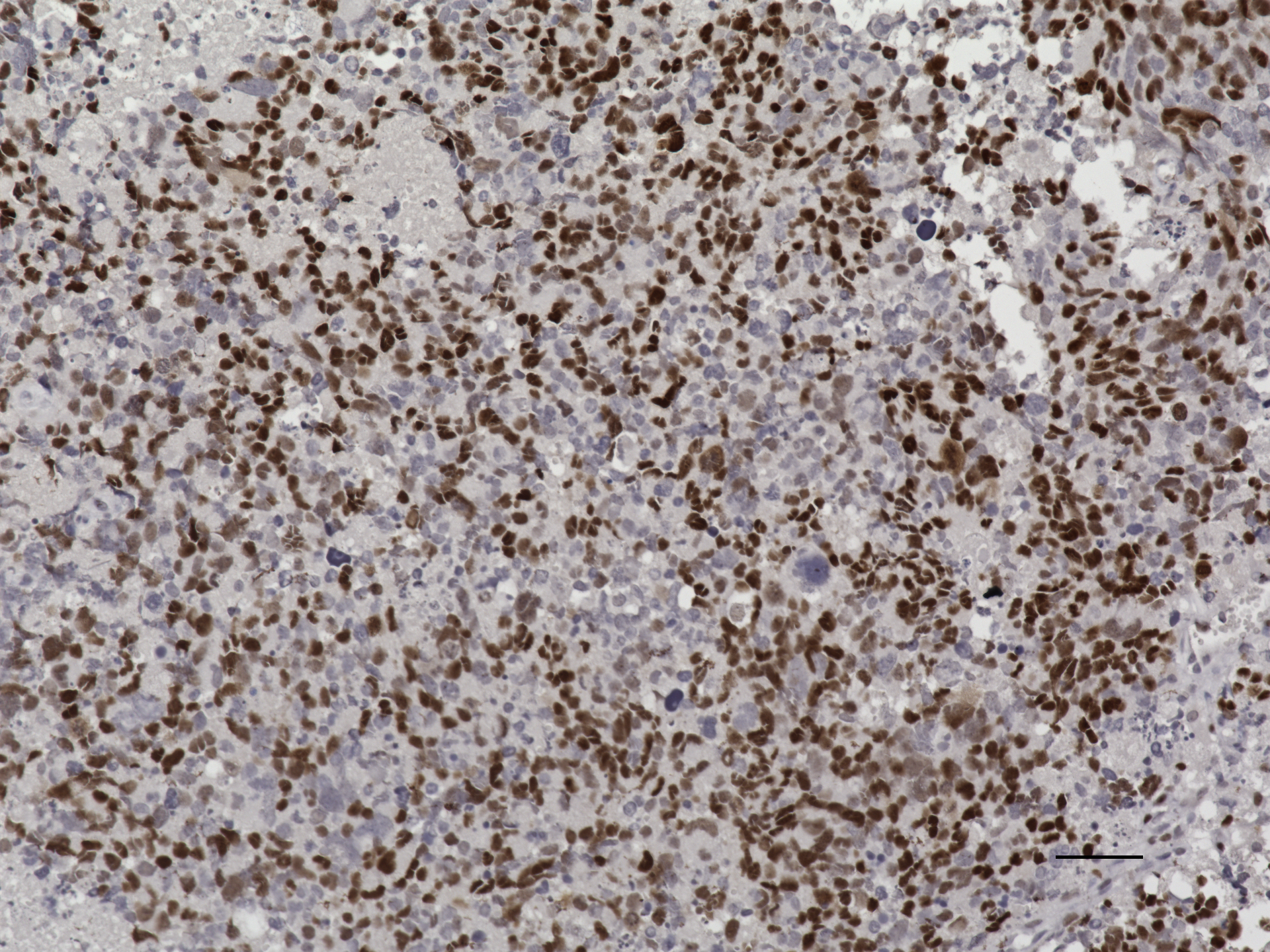

### IPDrosha cerit pS6 20X 50um.jpeg

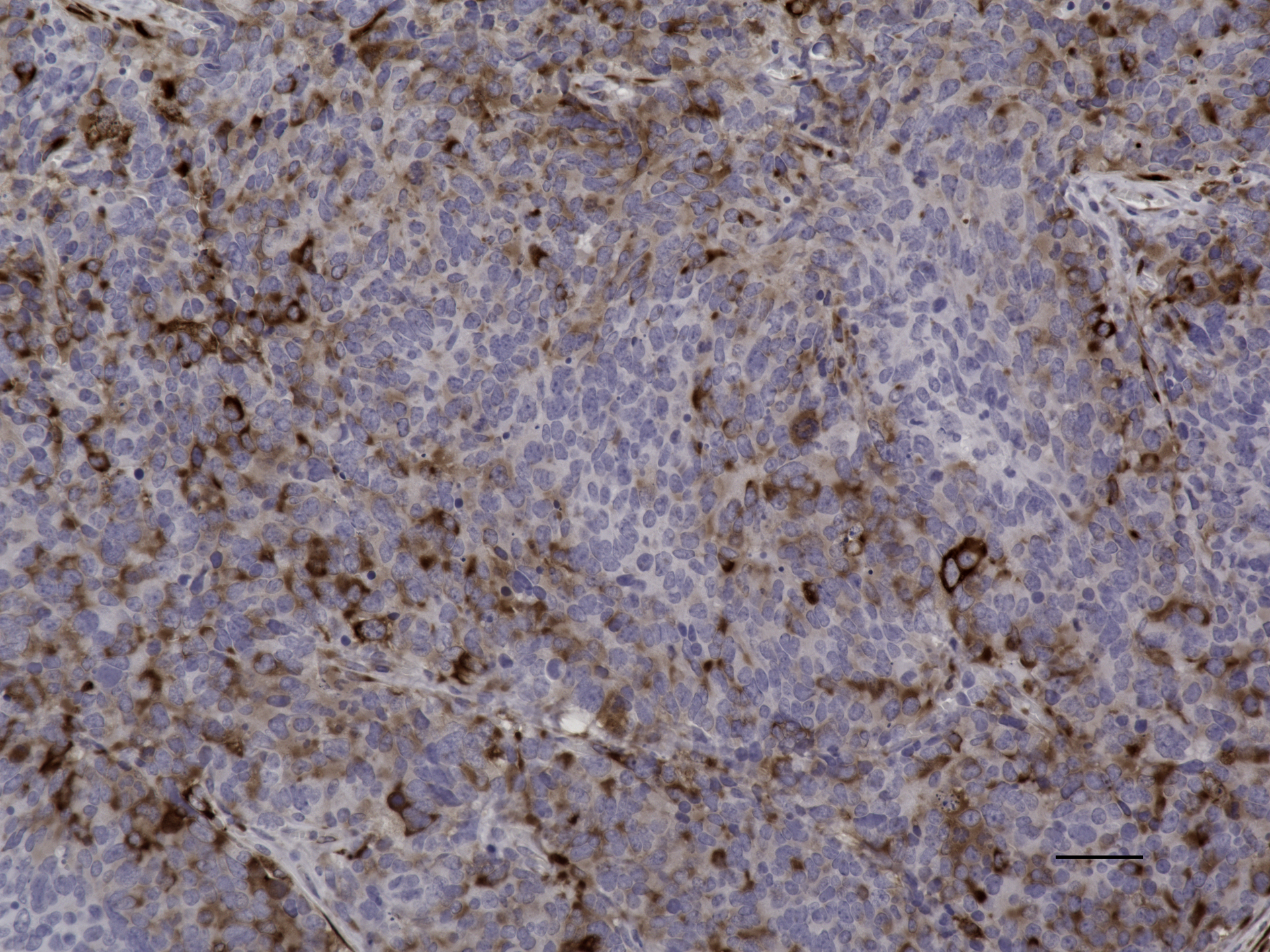

### IPDrosha cerit pS6 20X.jpeg

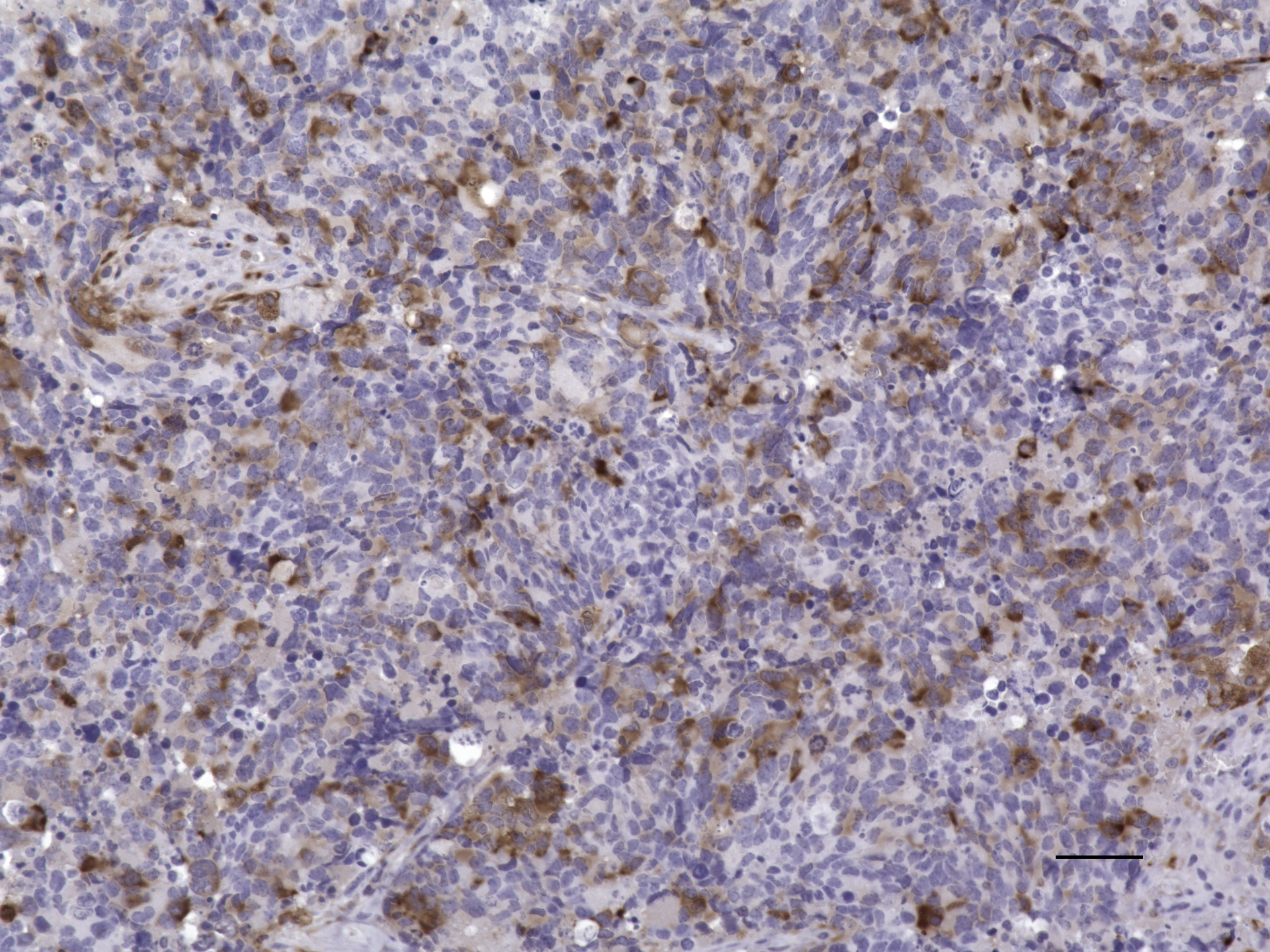

### IPDrosha combo 20X H_E.jpeg

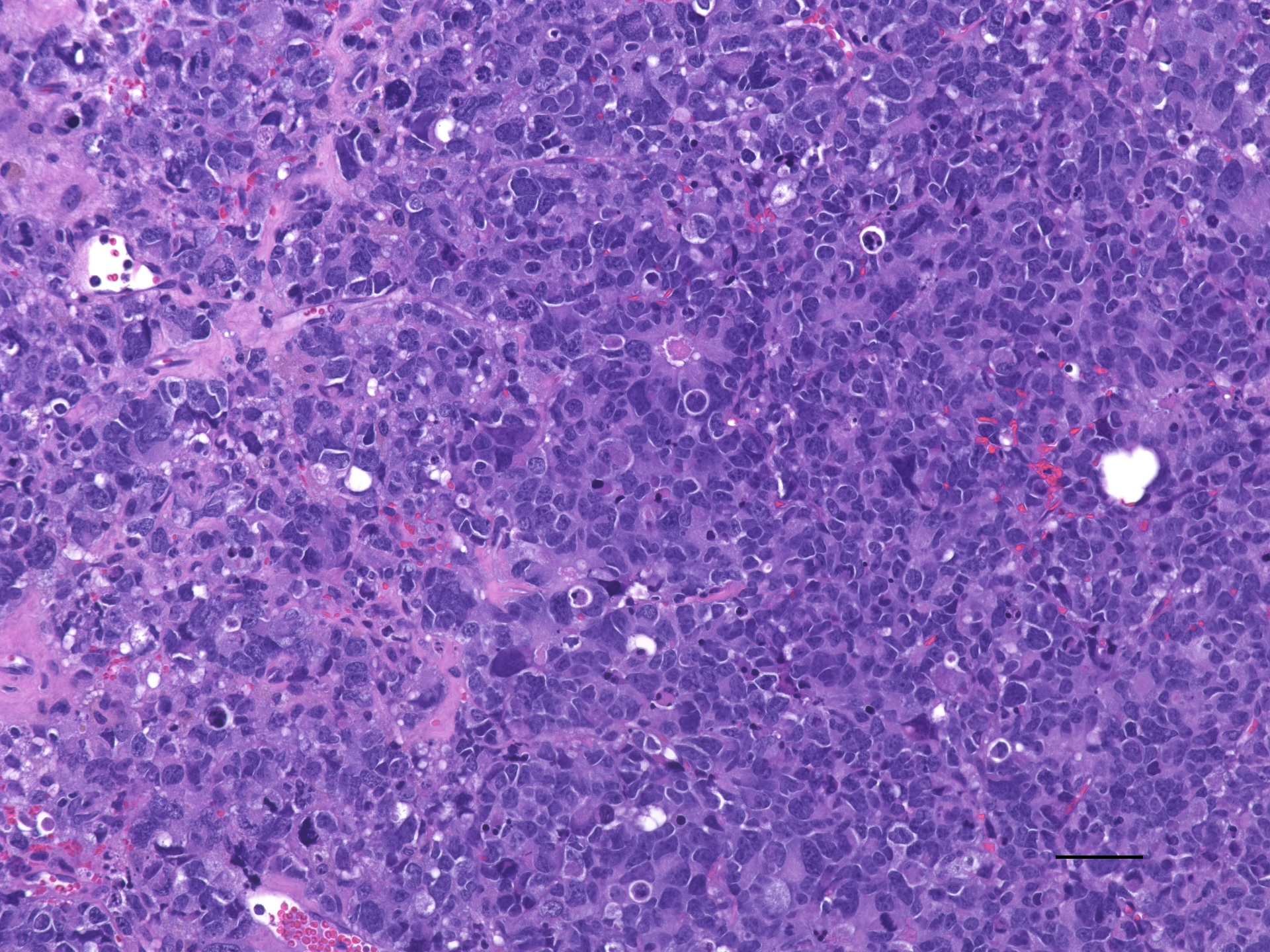

### IPDrosha combo Ki67 20x.jpeg

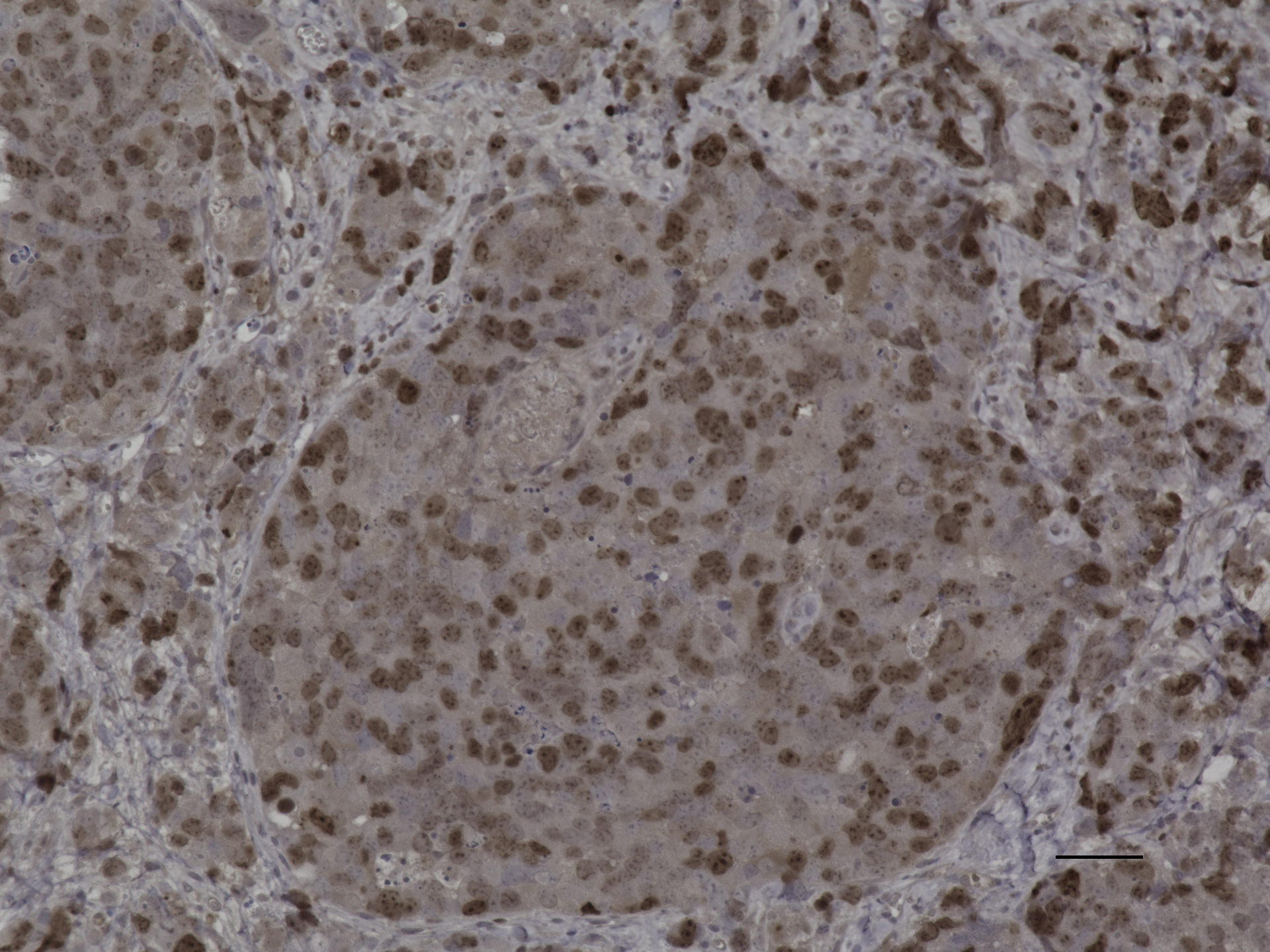

### IPDrosha combo pRb 20X 50um.jpeg

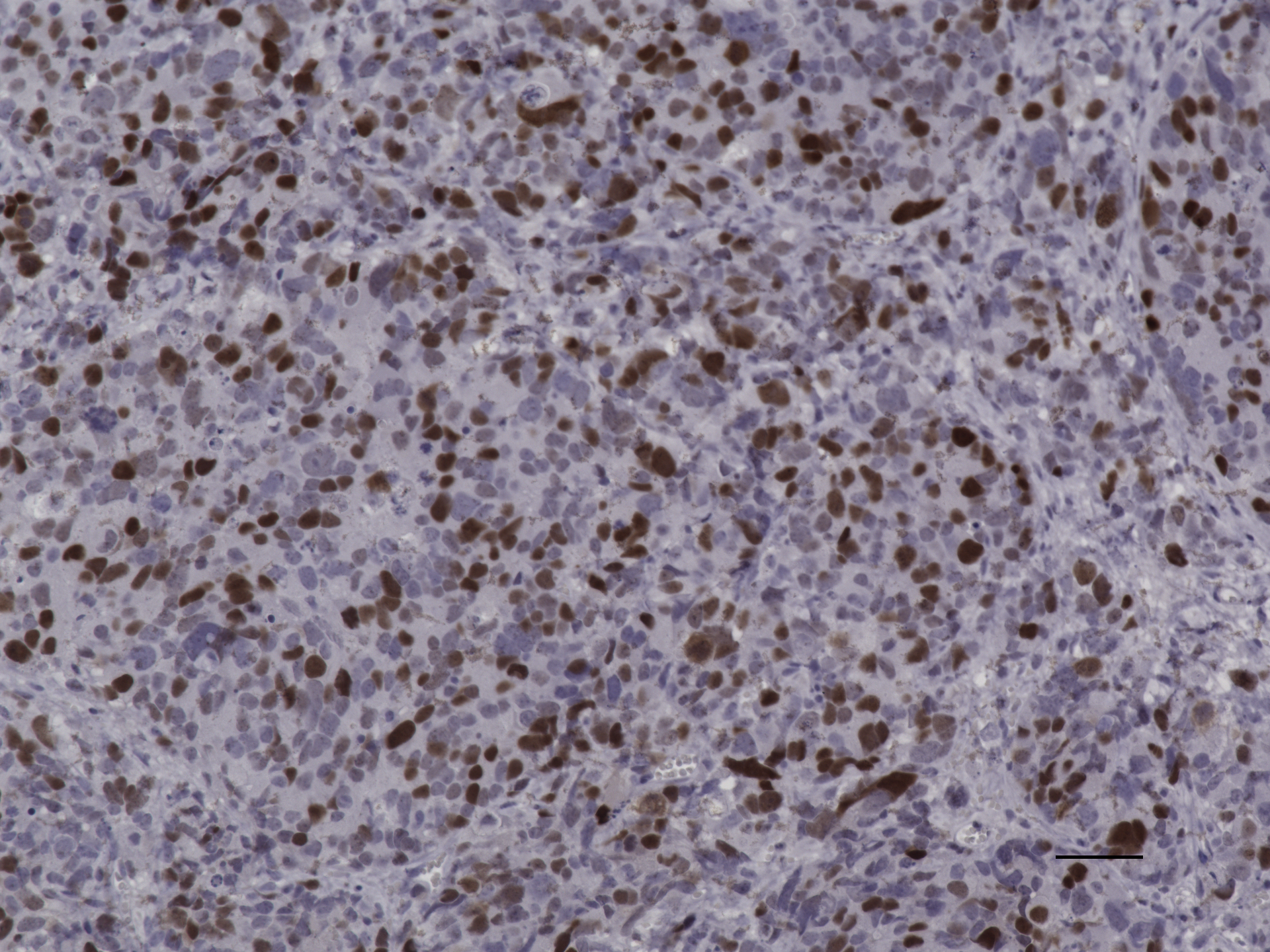

### IPDrosha combo pS6 20X 50um.jpeg

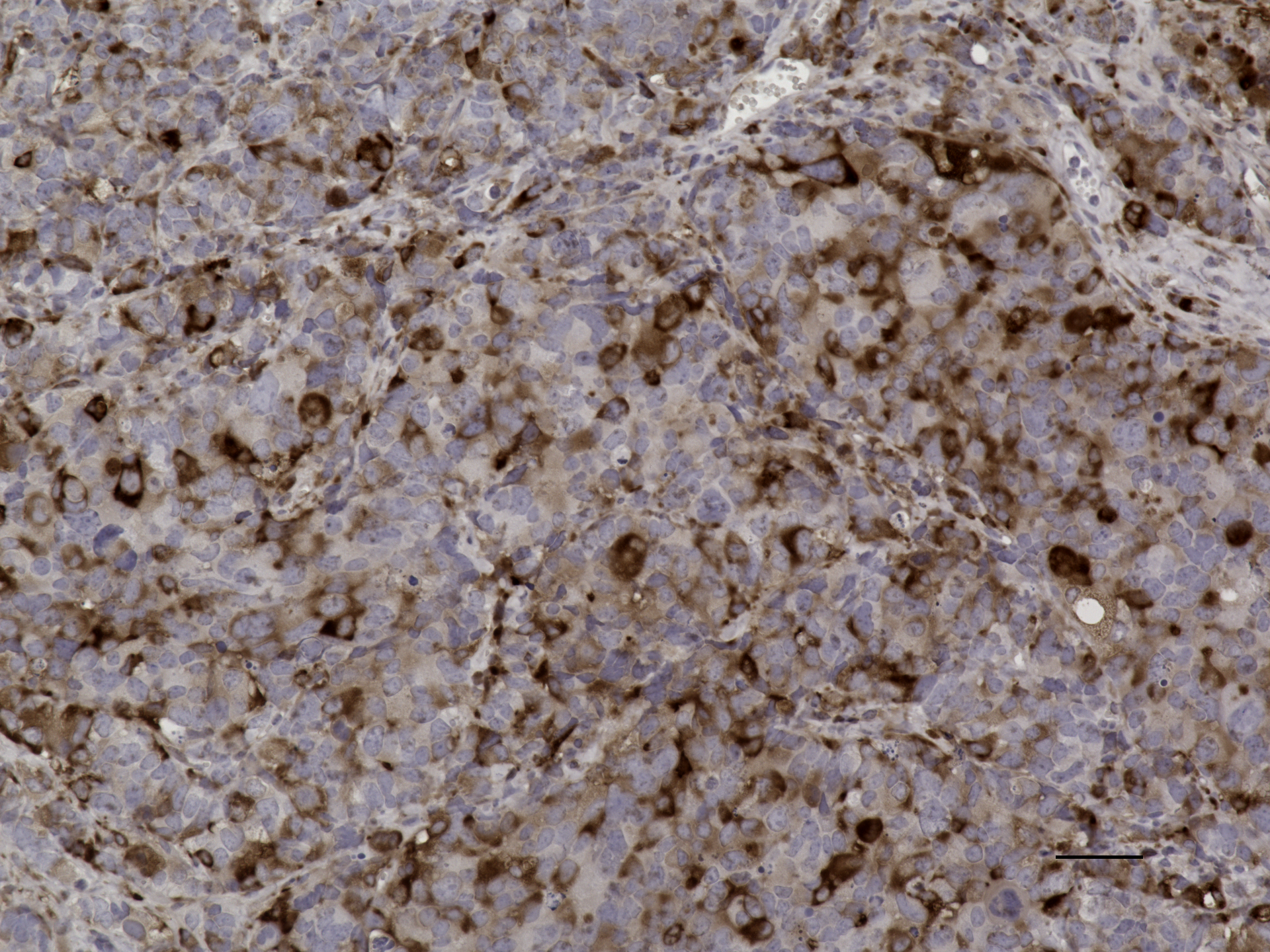

### IPDrosha GFAP 50 um 20X.jpeg

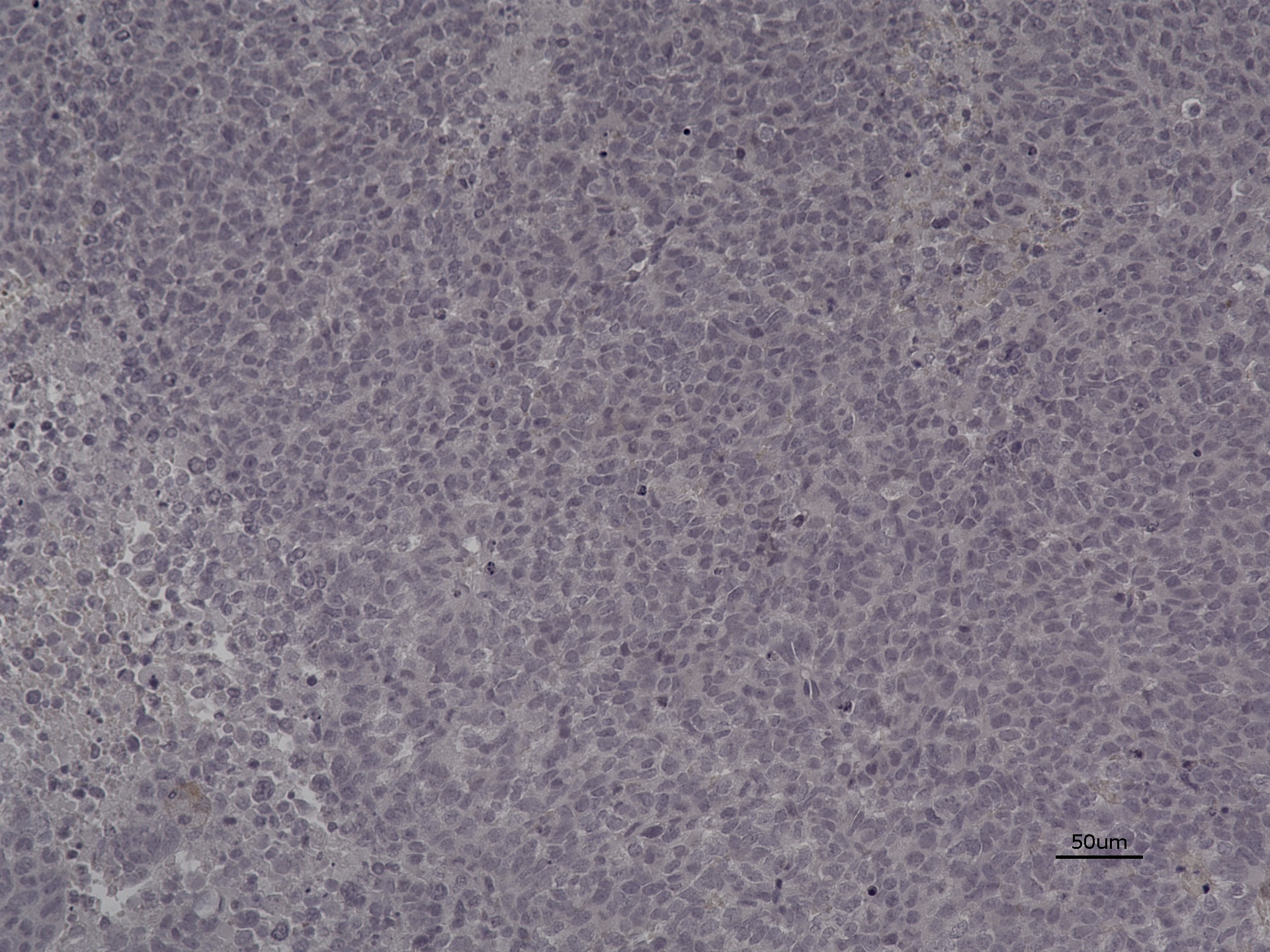

### IPDrosha Ki67 20X.jpeg

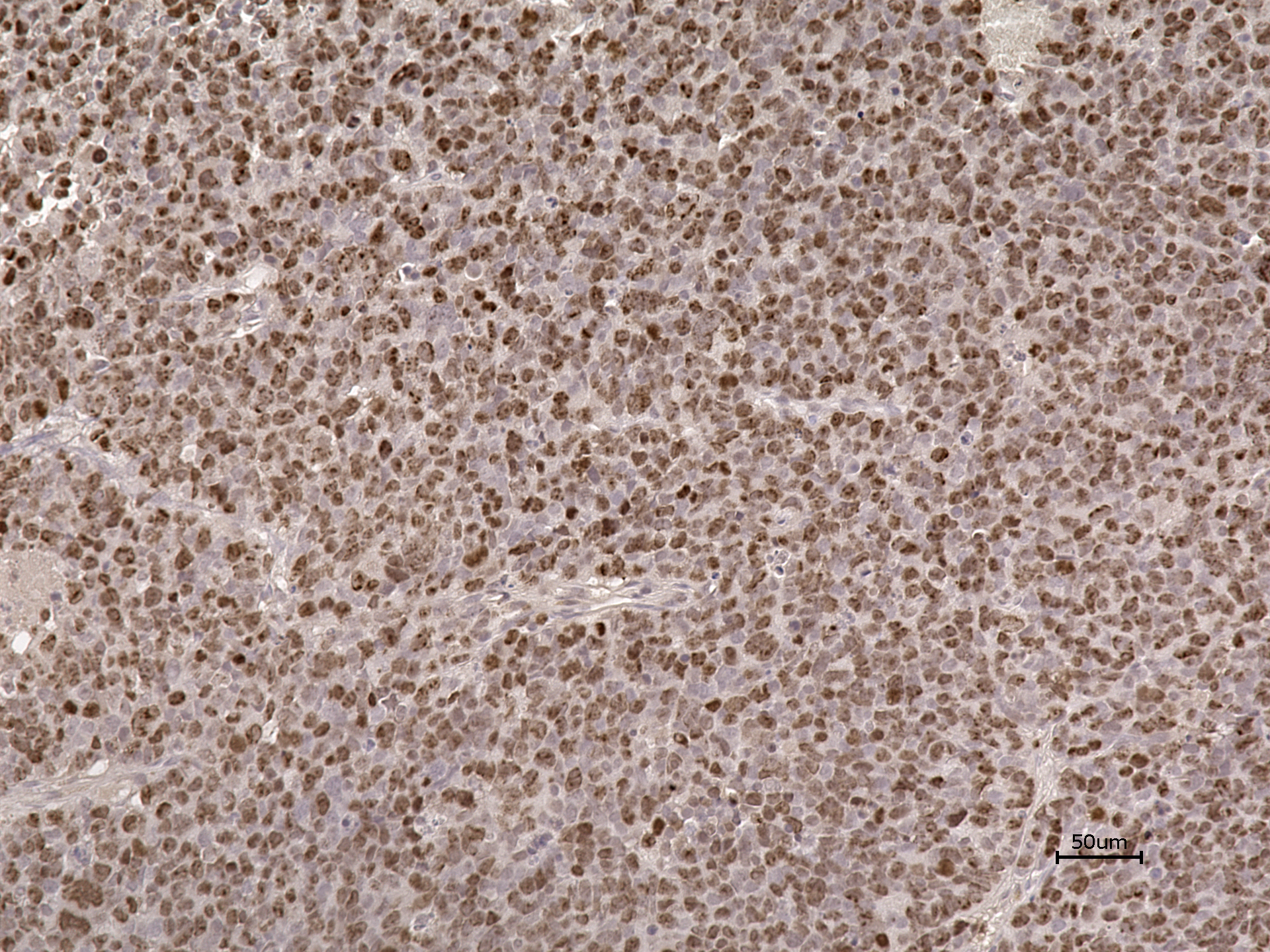

### IPDrosha palbo 20X H_E.jpeg

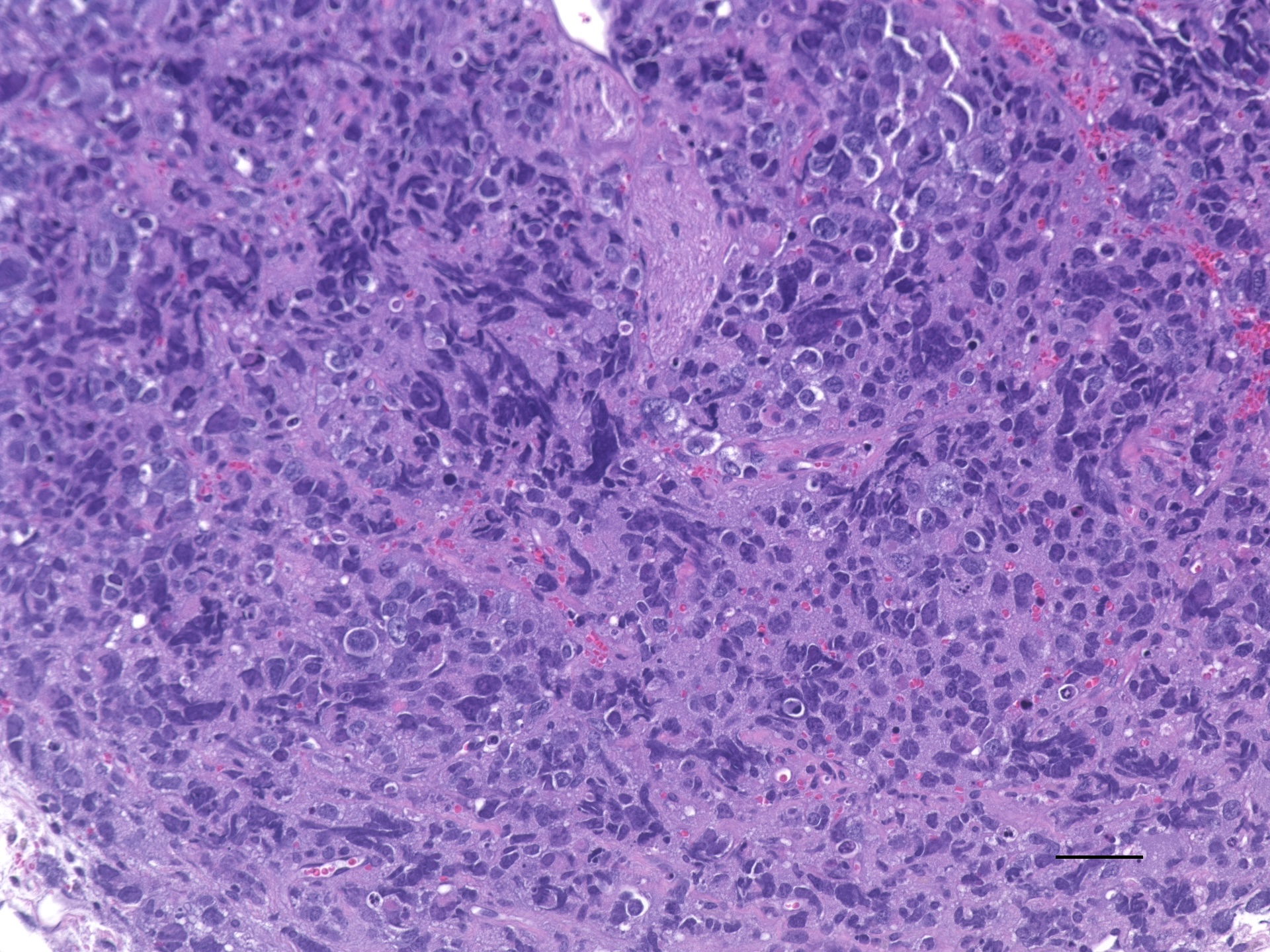

### IPDrosha palbo Ki67 20x.jpeg

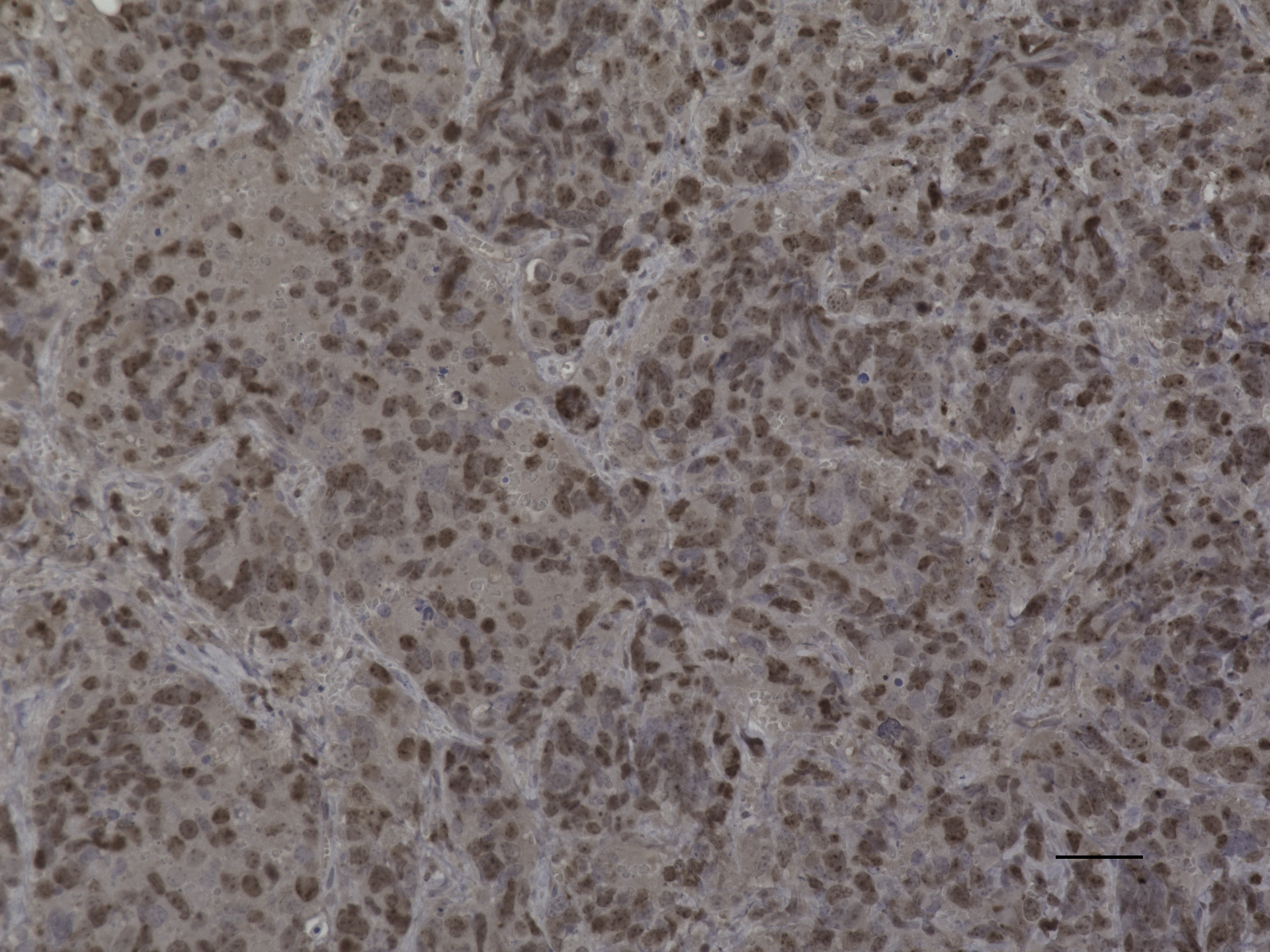

### IPDrosha palbo pRb 20X 50um.jpeg

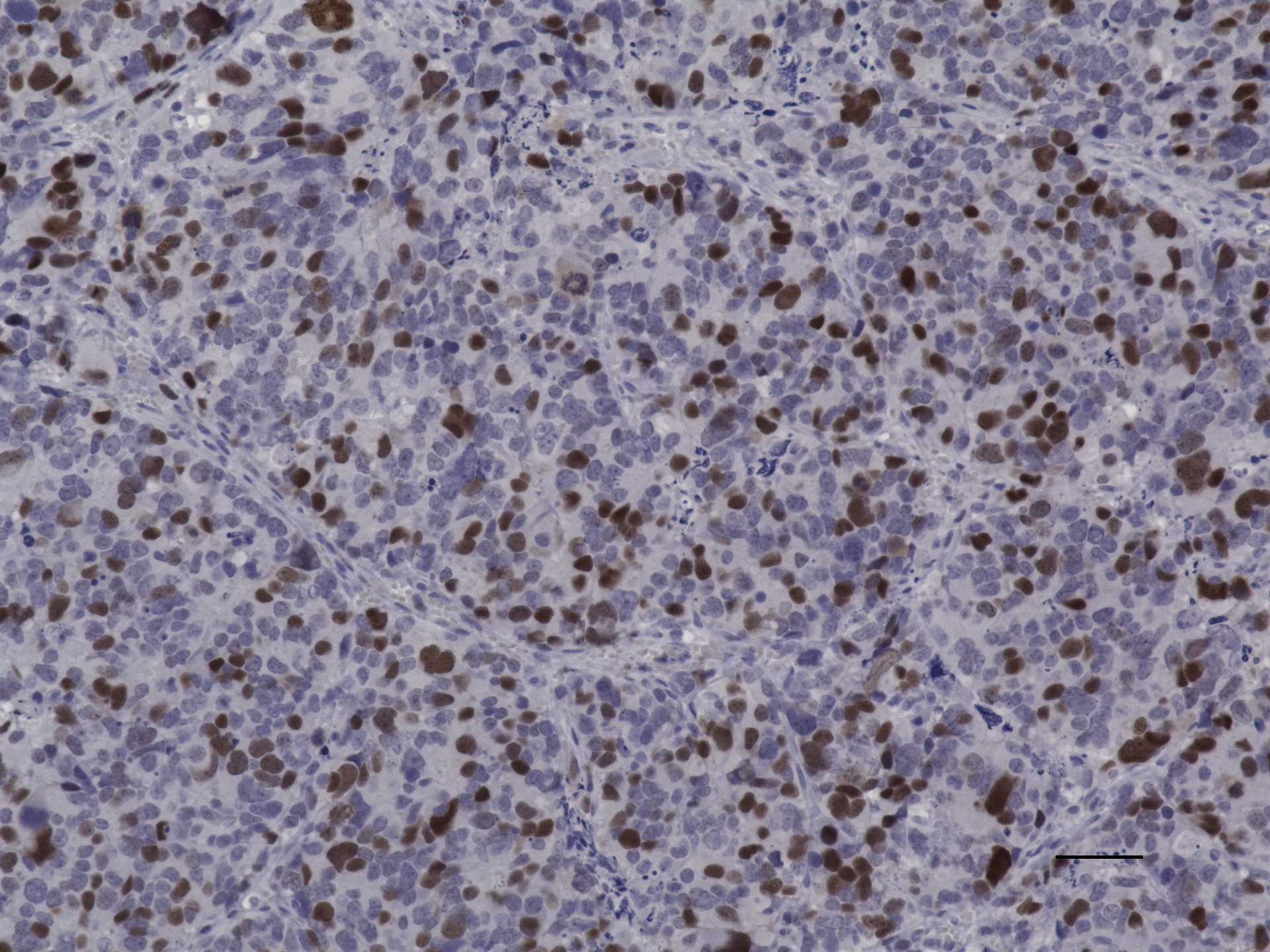

### IPDrosha palbo pRb 20X.jpeg

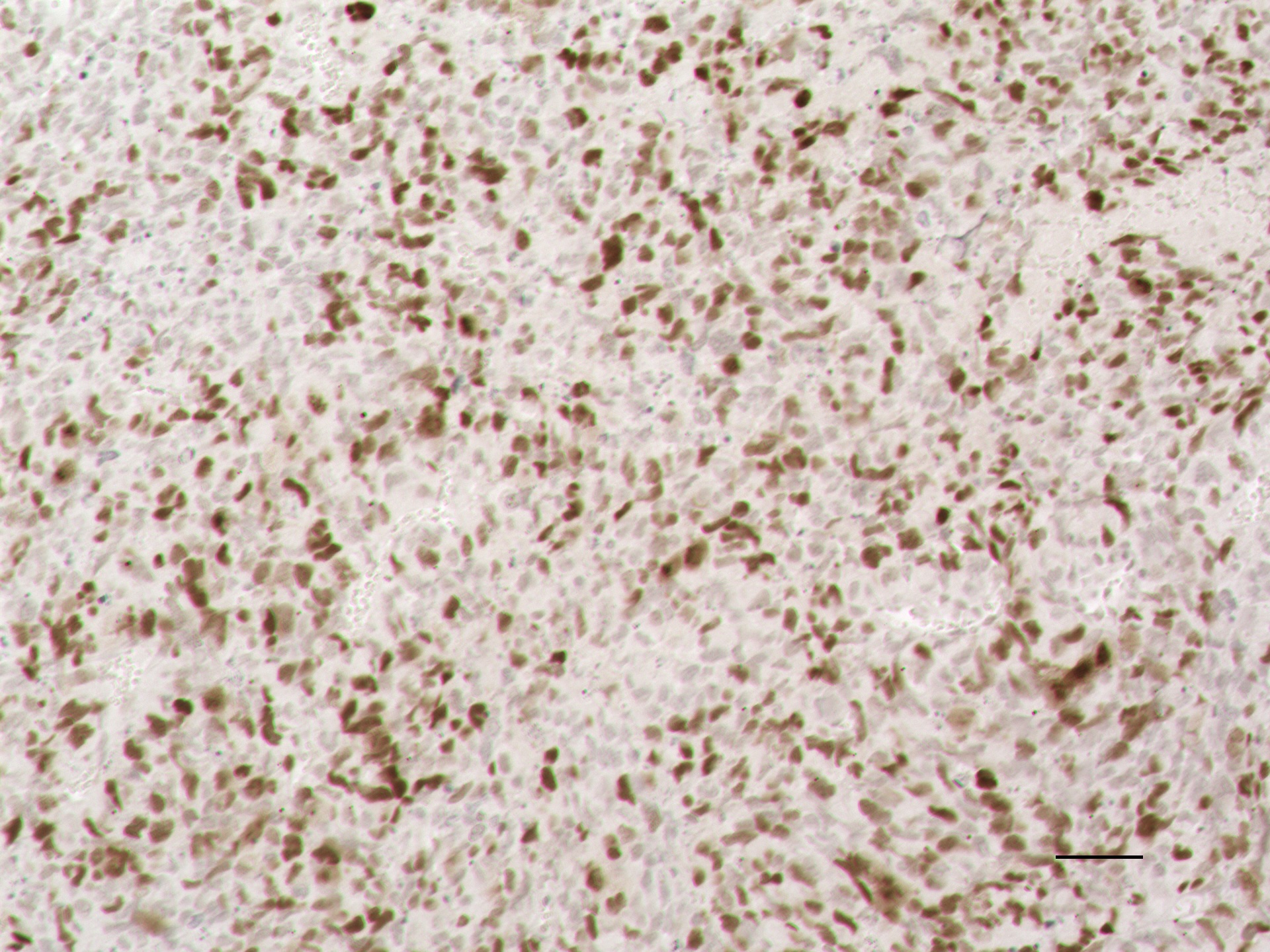

### IPDrosha palbo pS6 20X 50um.jpeg

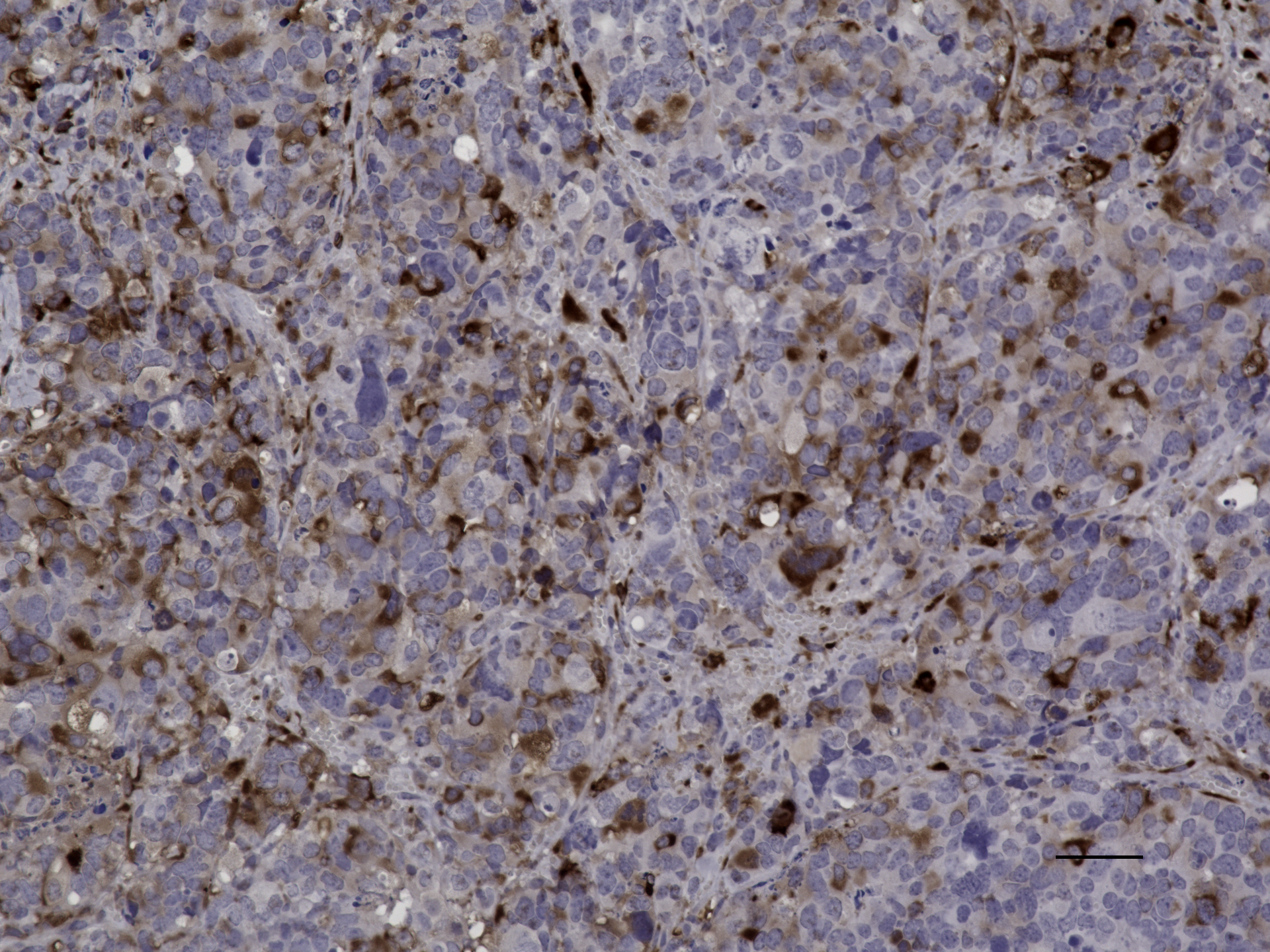

### IPDrosha palbociclib H_E 20X.jpeg

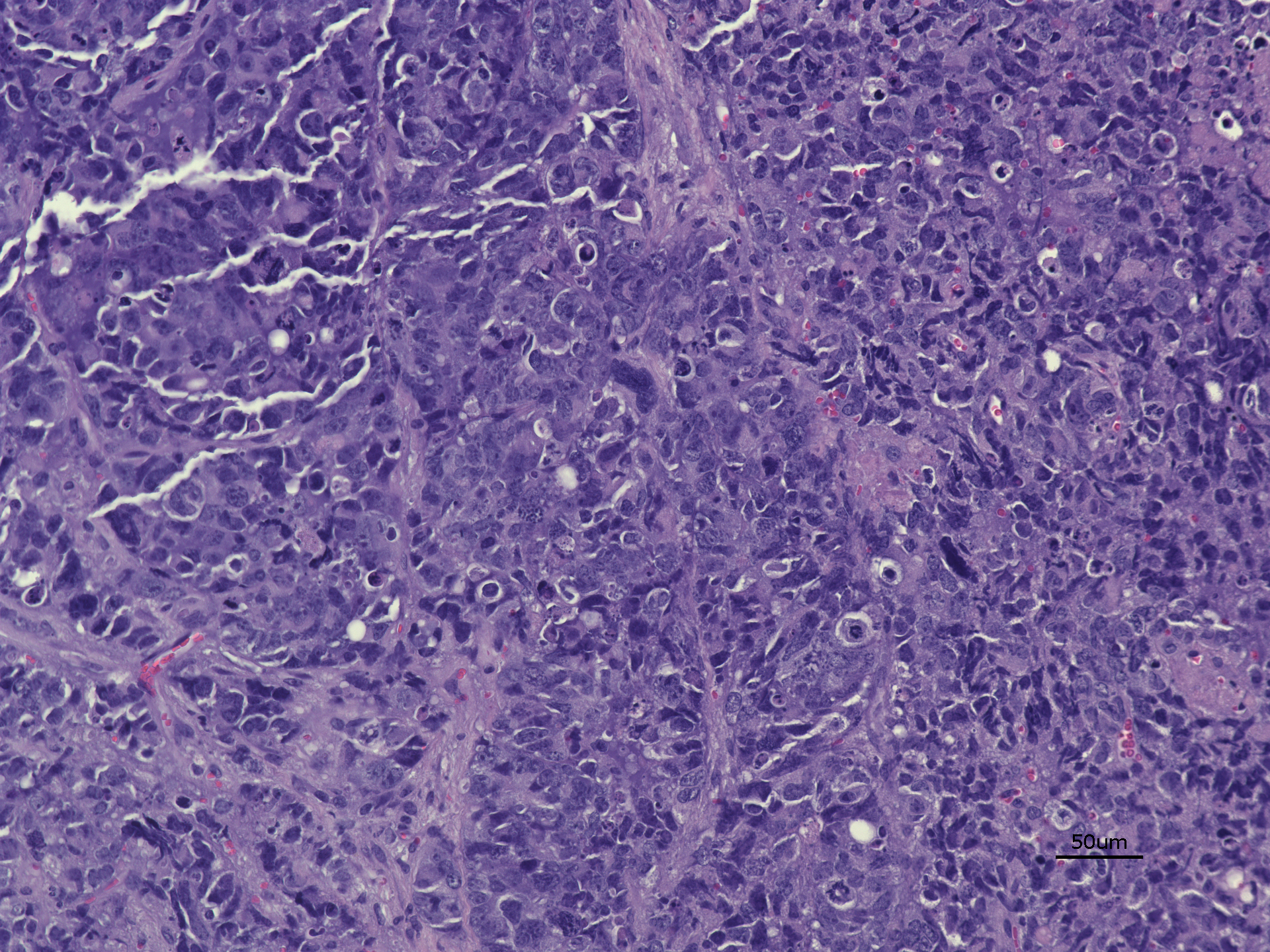

### IPDrosha palbociclib Ki67 20x.jpeg

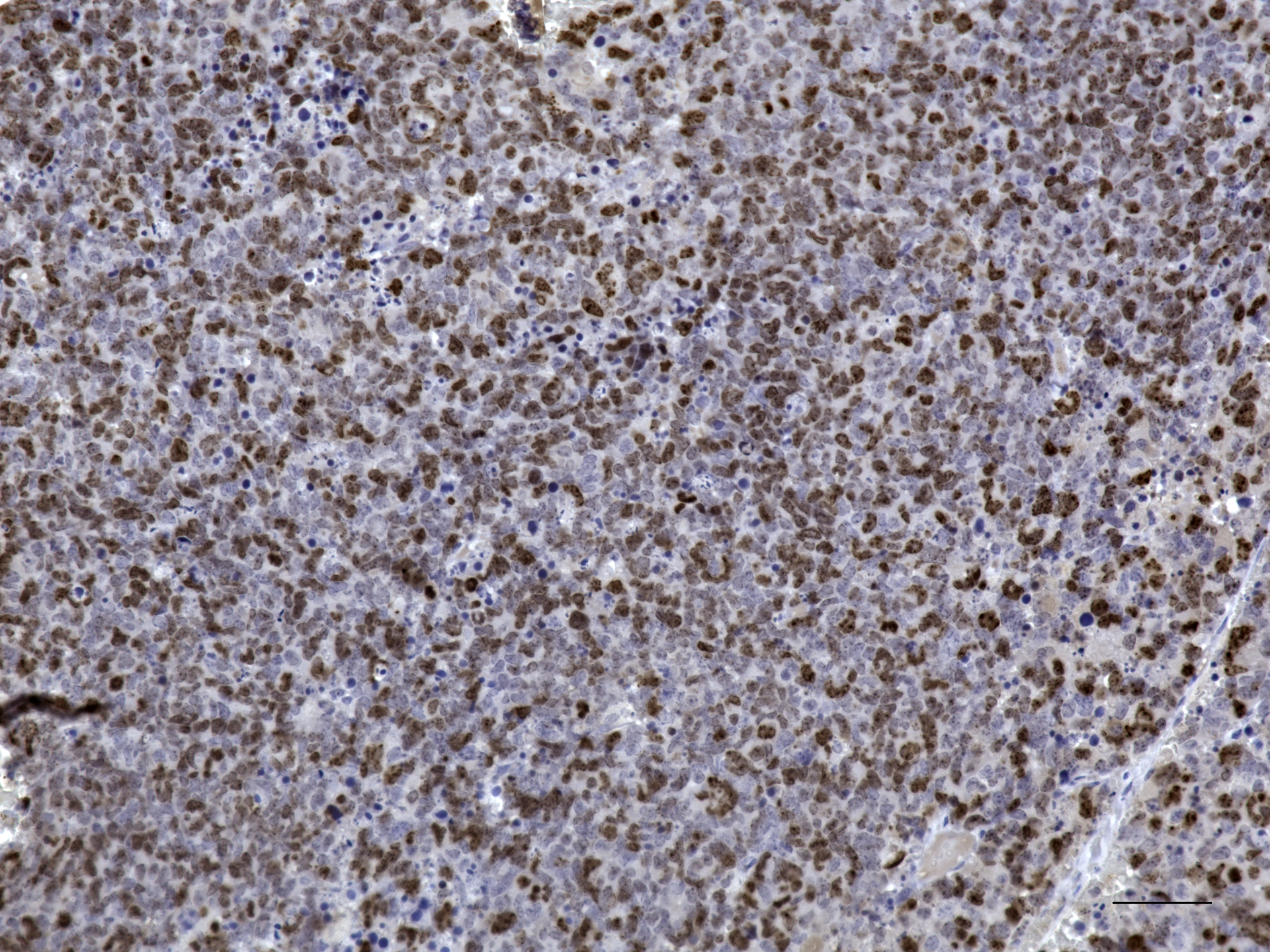

### IPDrosha pRb_20X.jpg

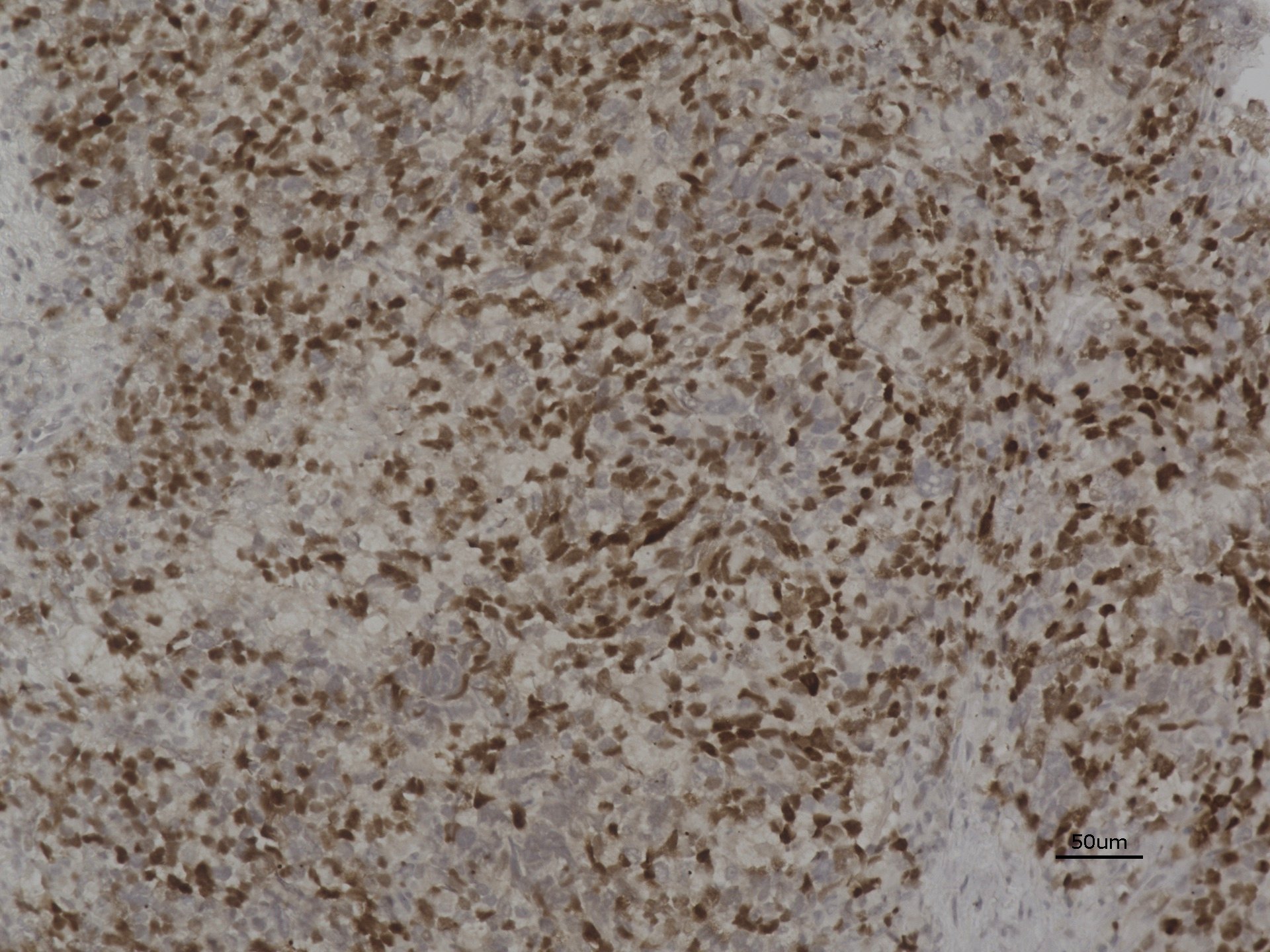

### IPDrosha synaptophysin 20x.jpeg

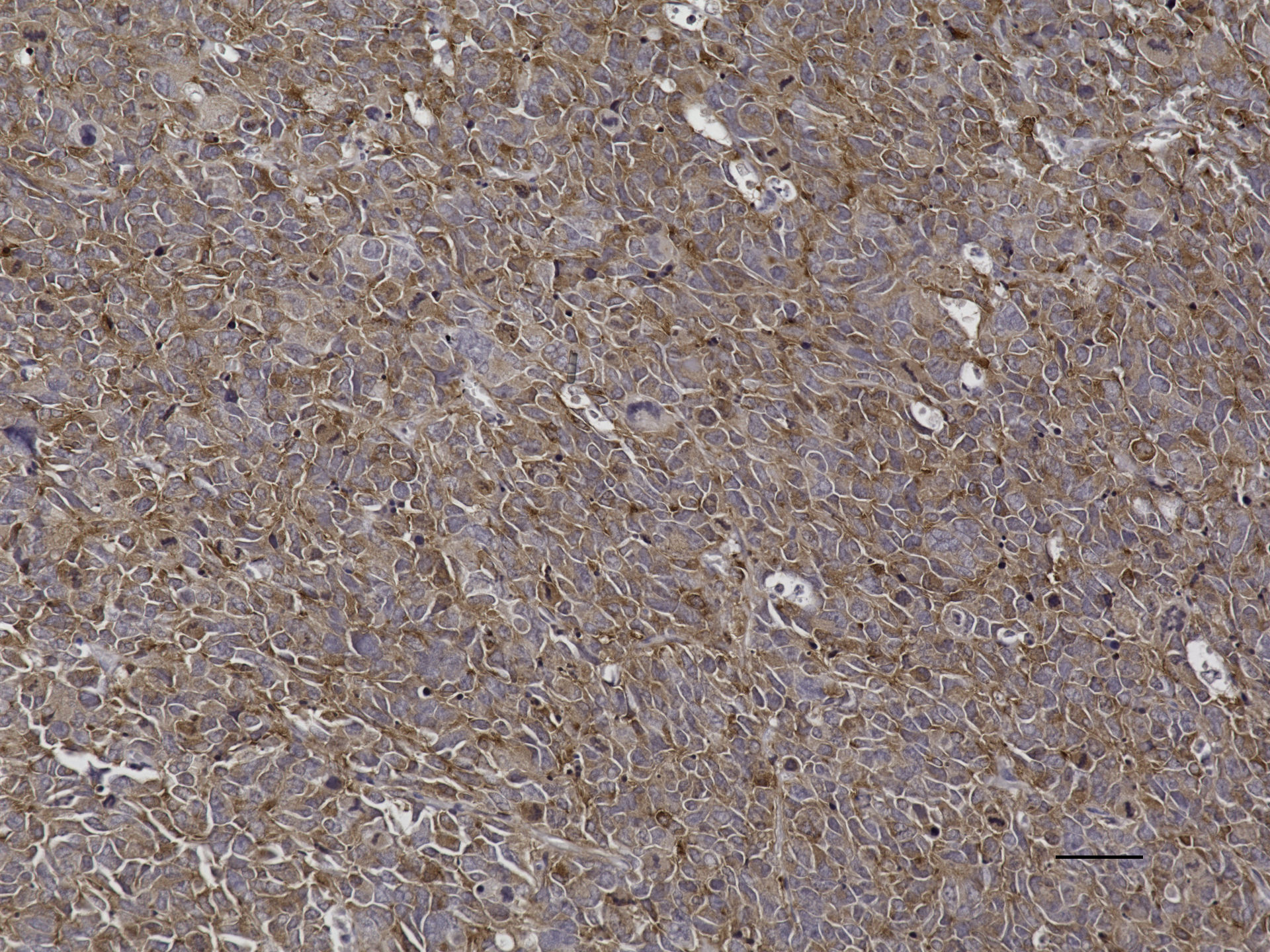

### IPDrosha vehicle 20X H_E.jpeg

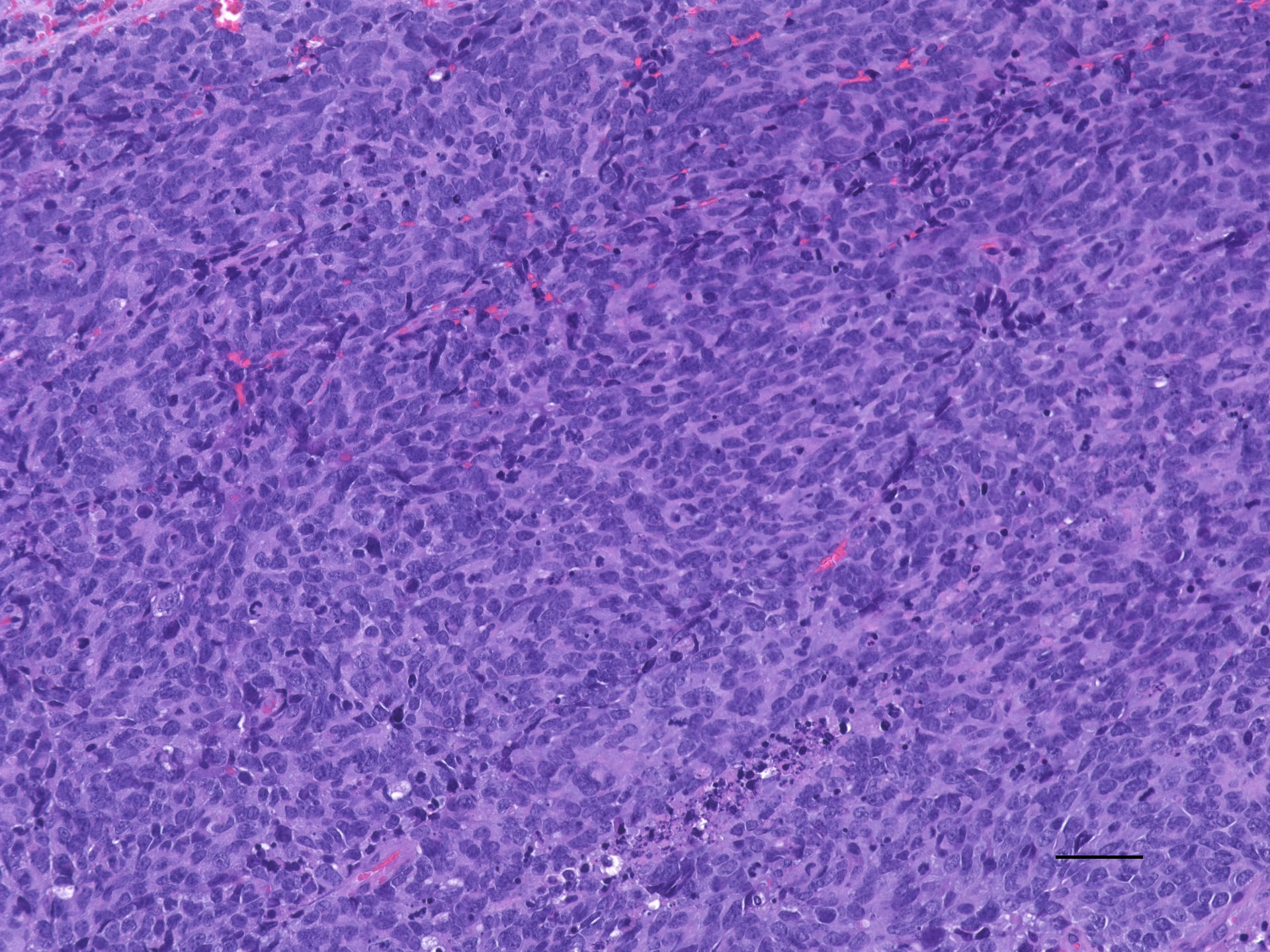

### IPDrosha vehicle H_E 20x.jpeg

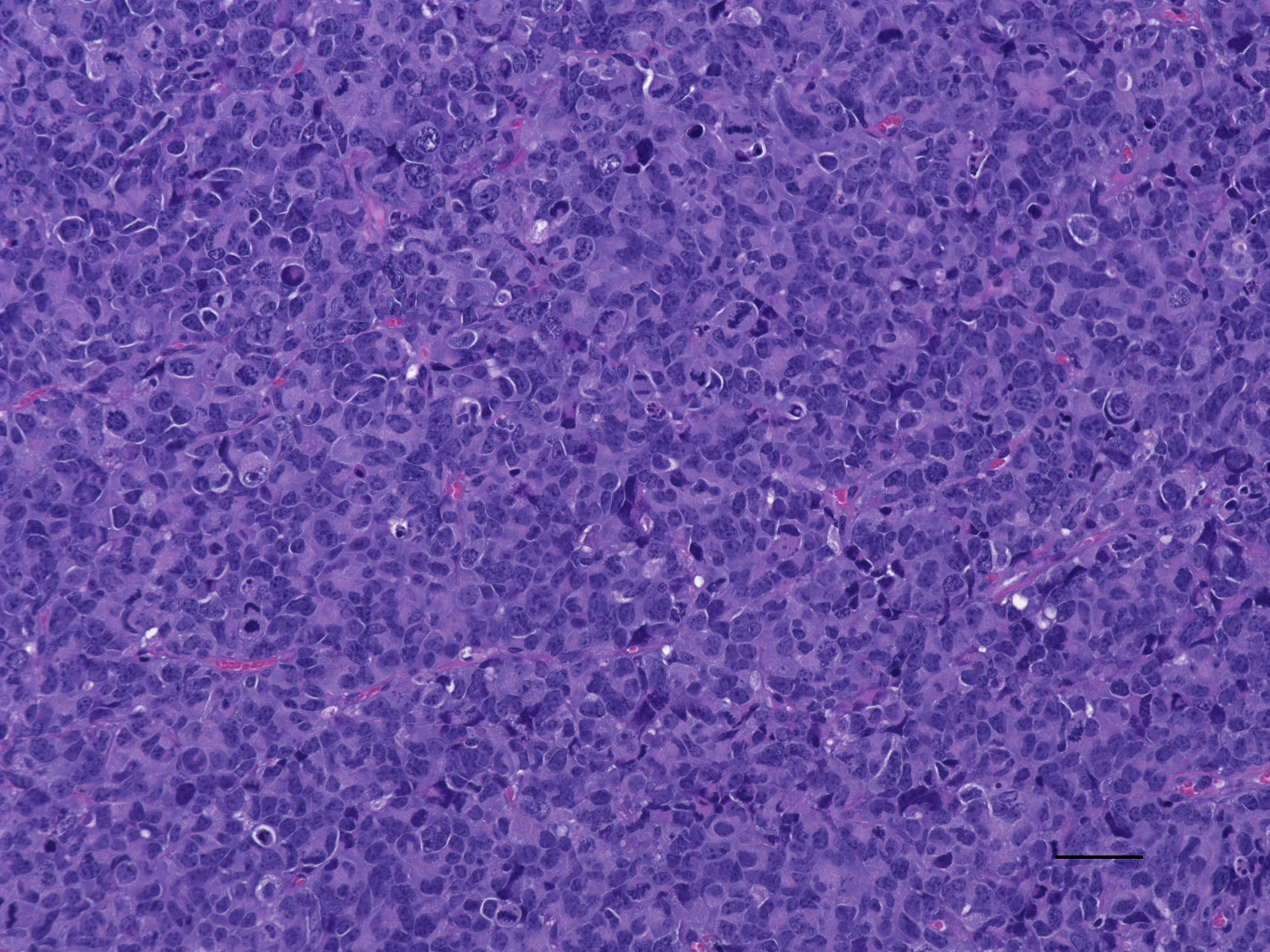

### IPDrosha vehicle H_E 20X.jpeg

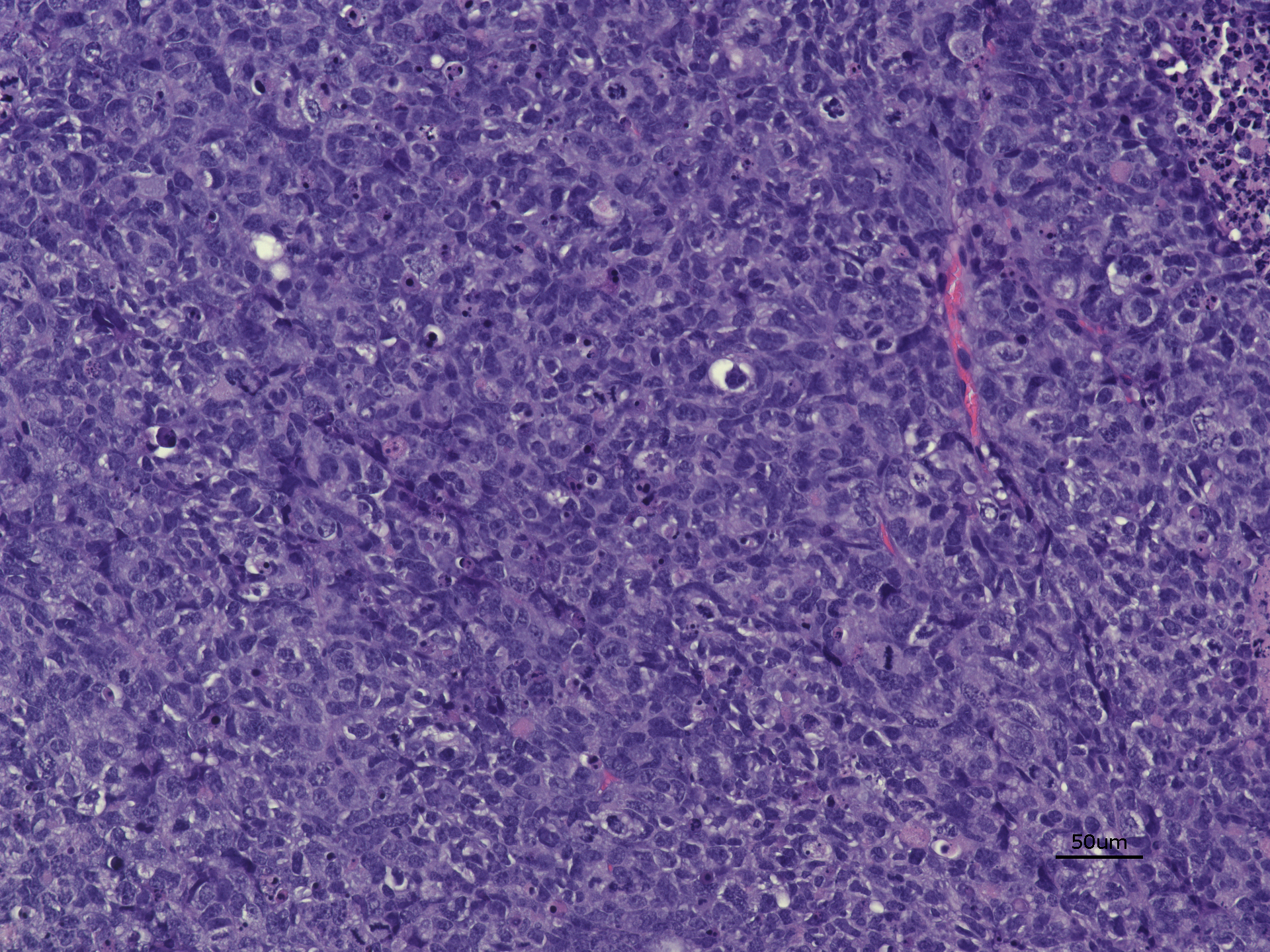

### IPDrosha vehicle Ki67 20x.jpeg

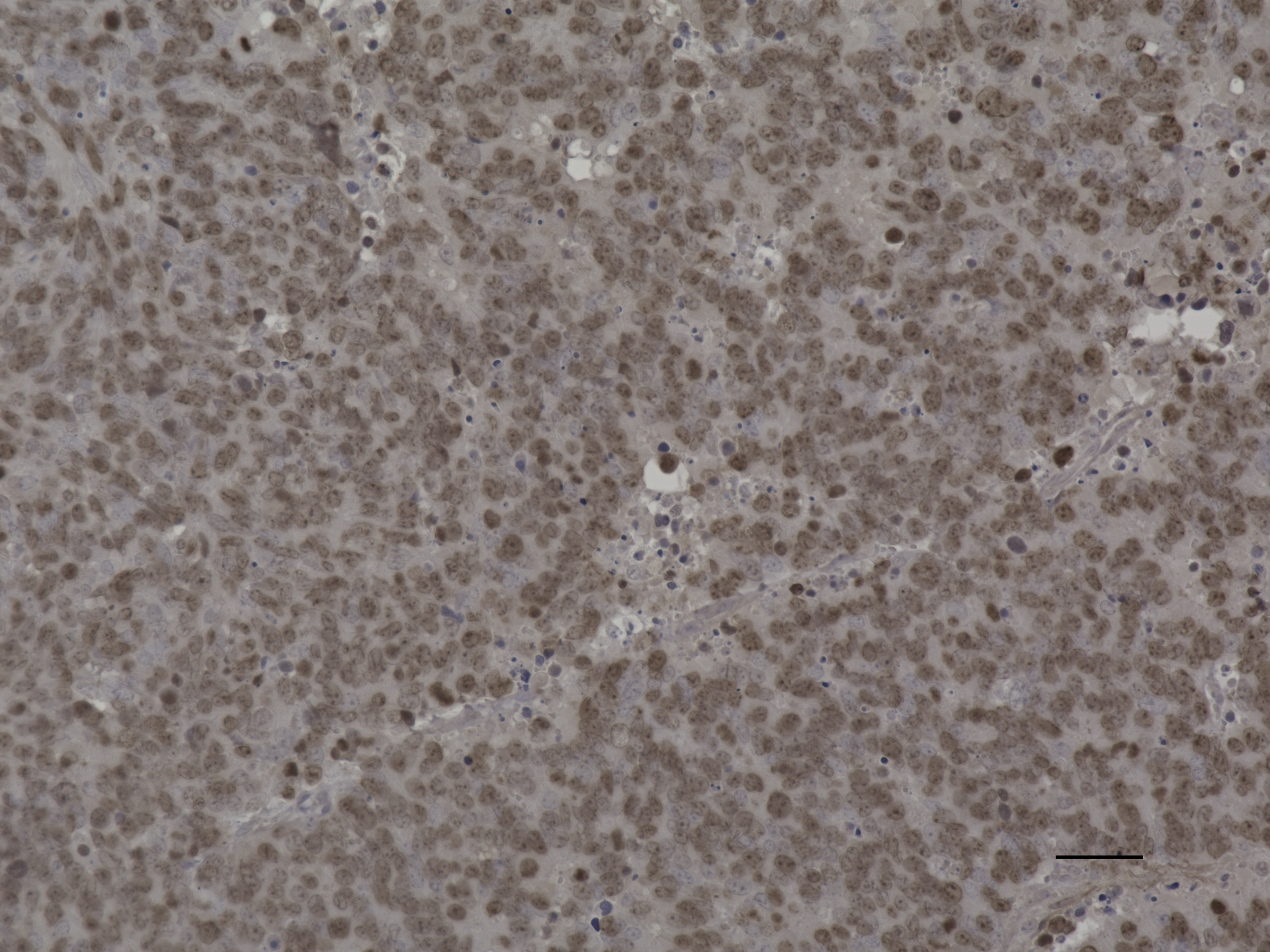

### IPDrosha vehicle Ki67 20x.jpeg

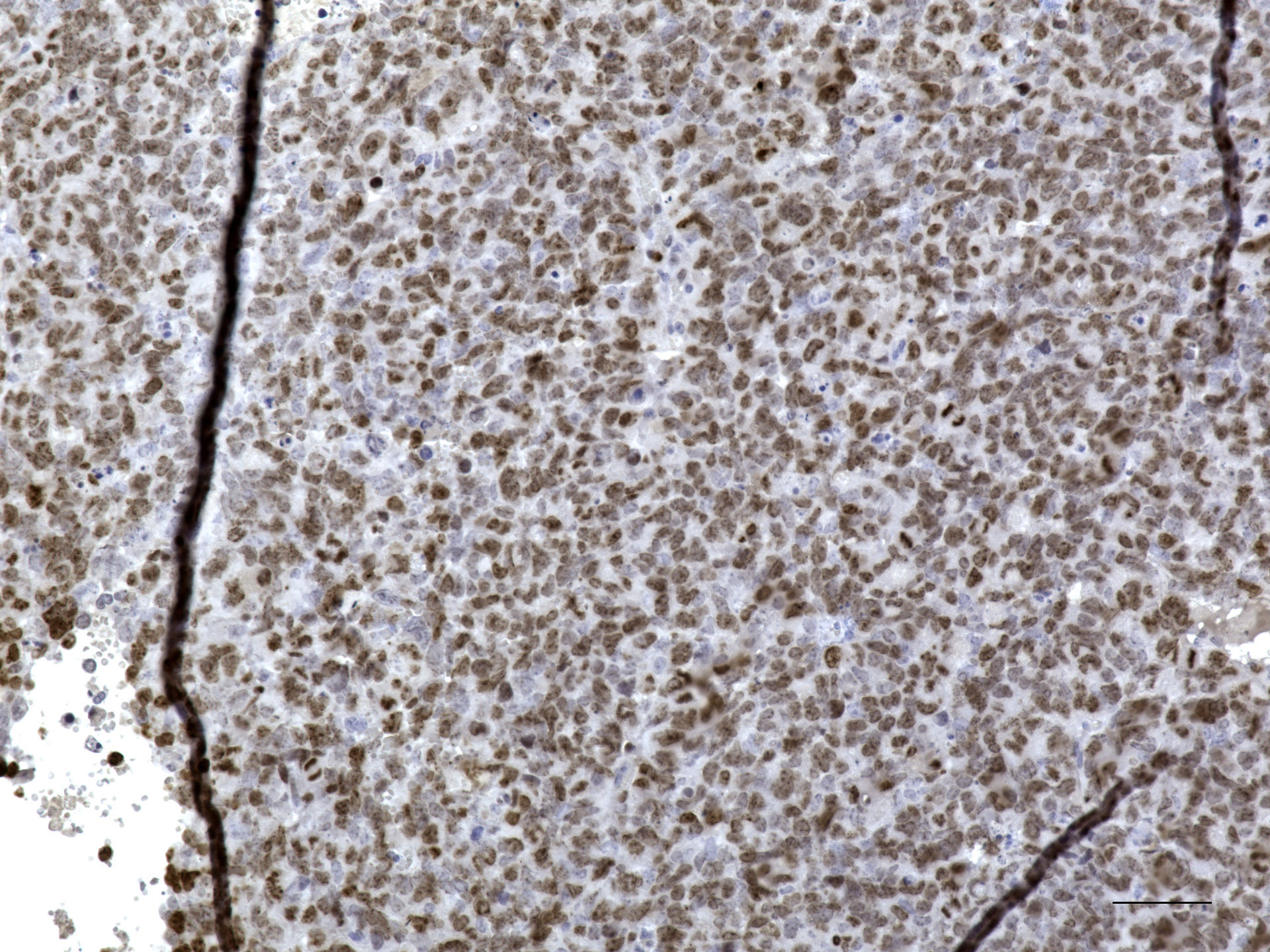

### IPDrosha vehicle pRb 20X 50um.jpeg

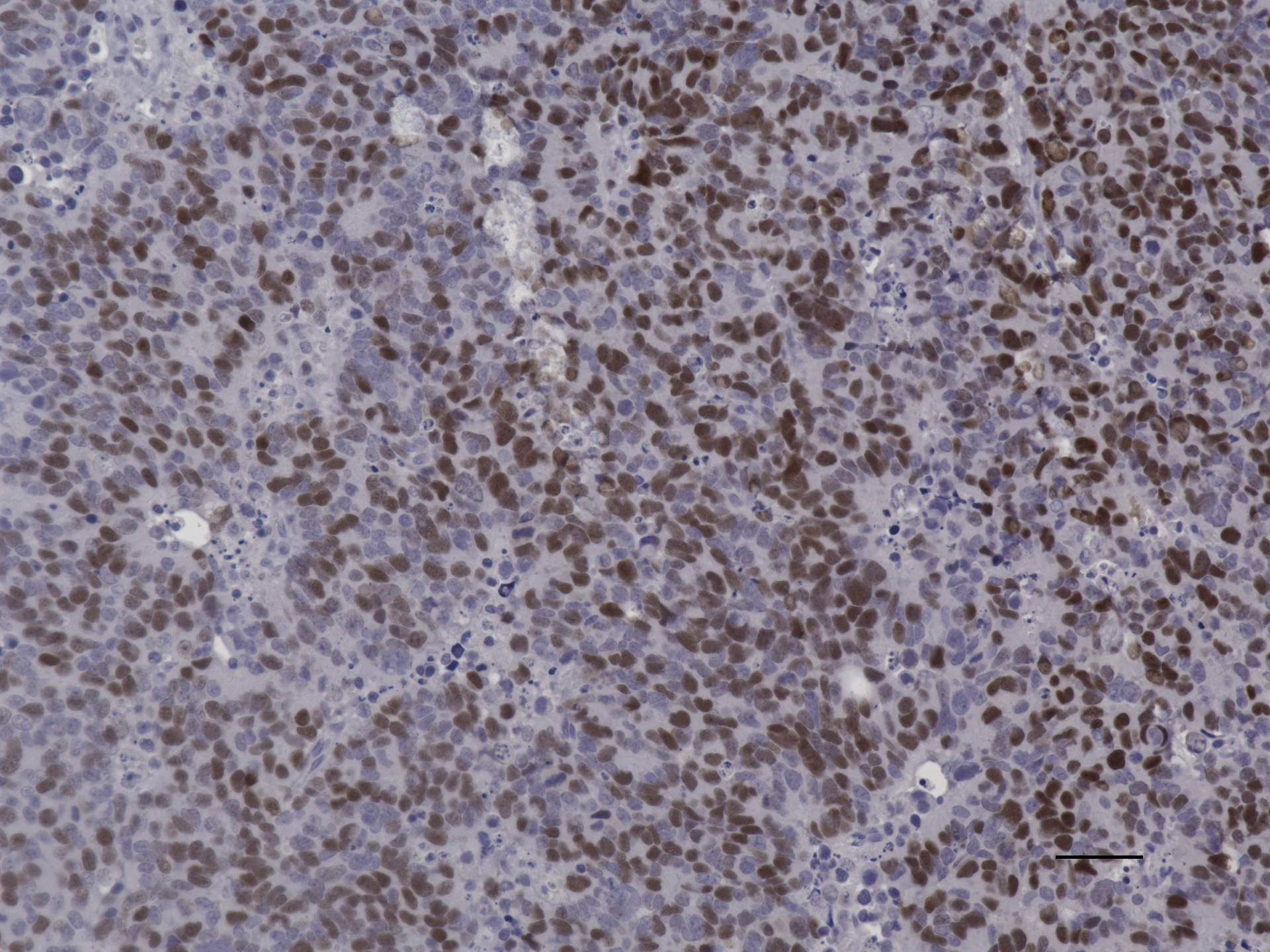

### IPDrosha vehicle pRb 20x.jpeg

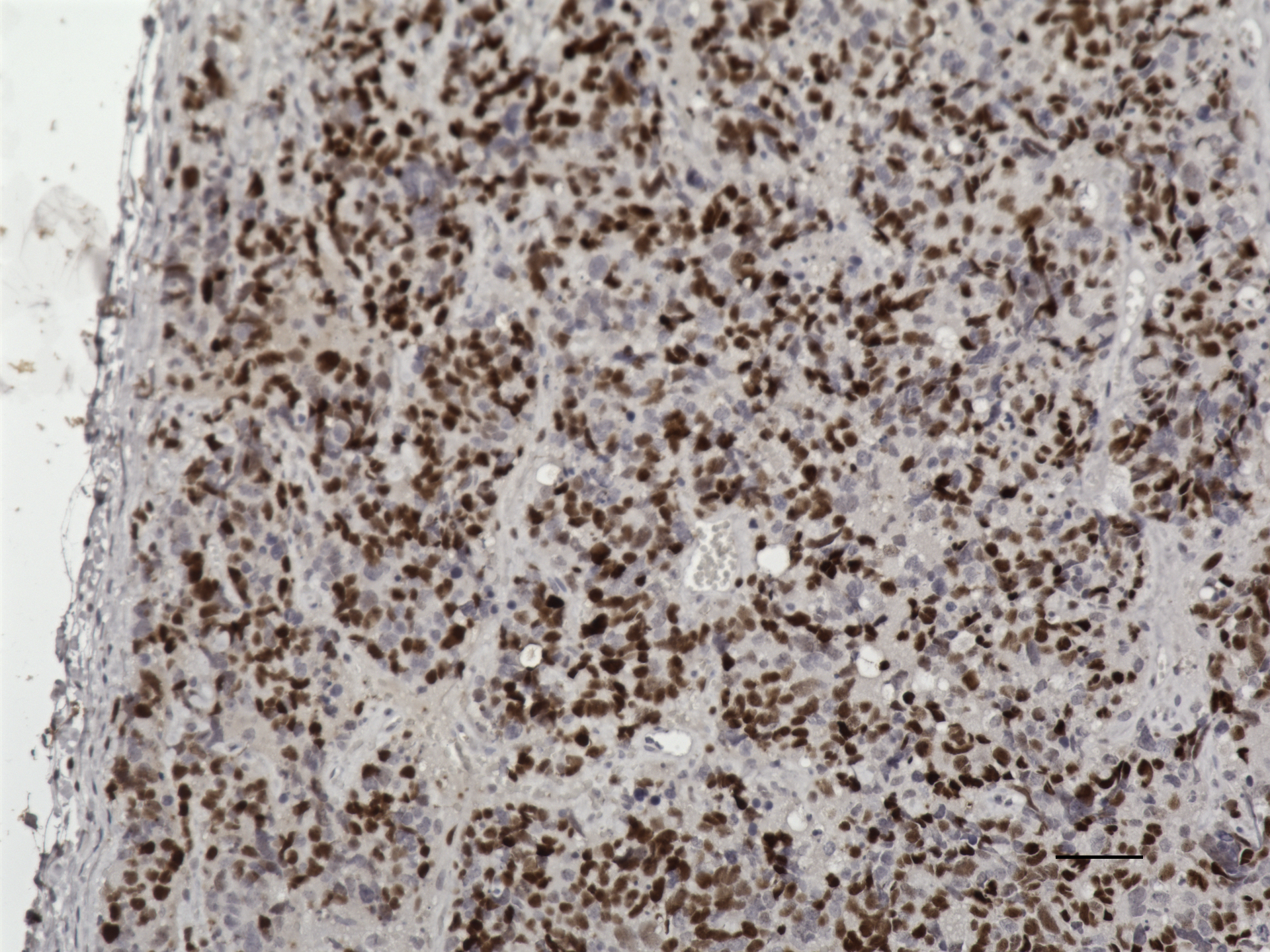

### IPDrosha vehicle pRb 20X.jpeg

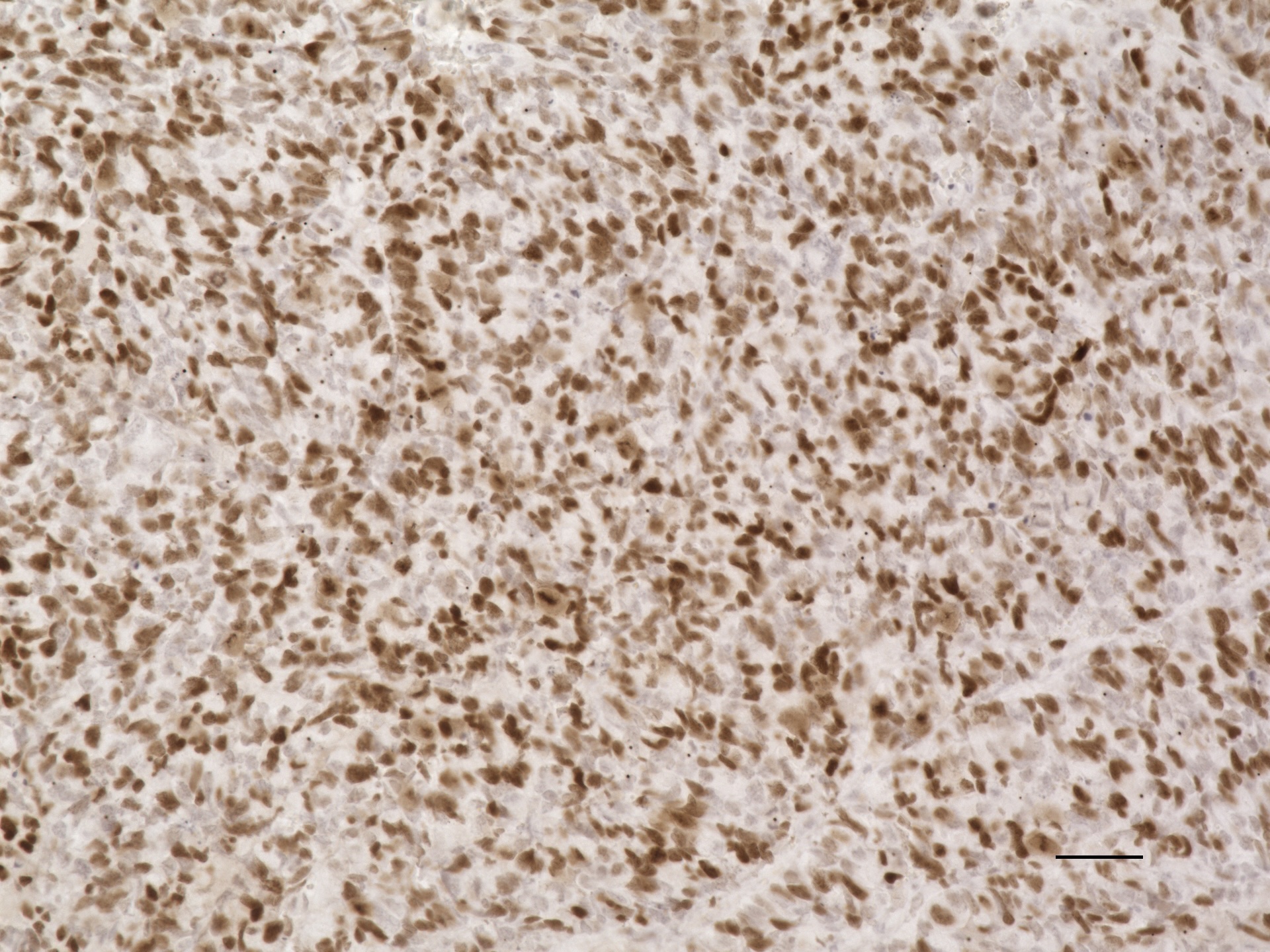

### IPDrosha vehicle pS6 20X 50um.jpeg

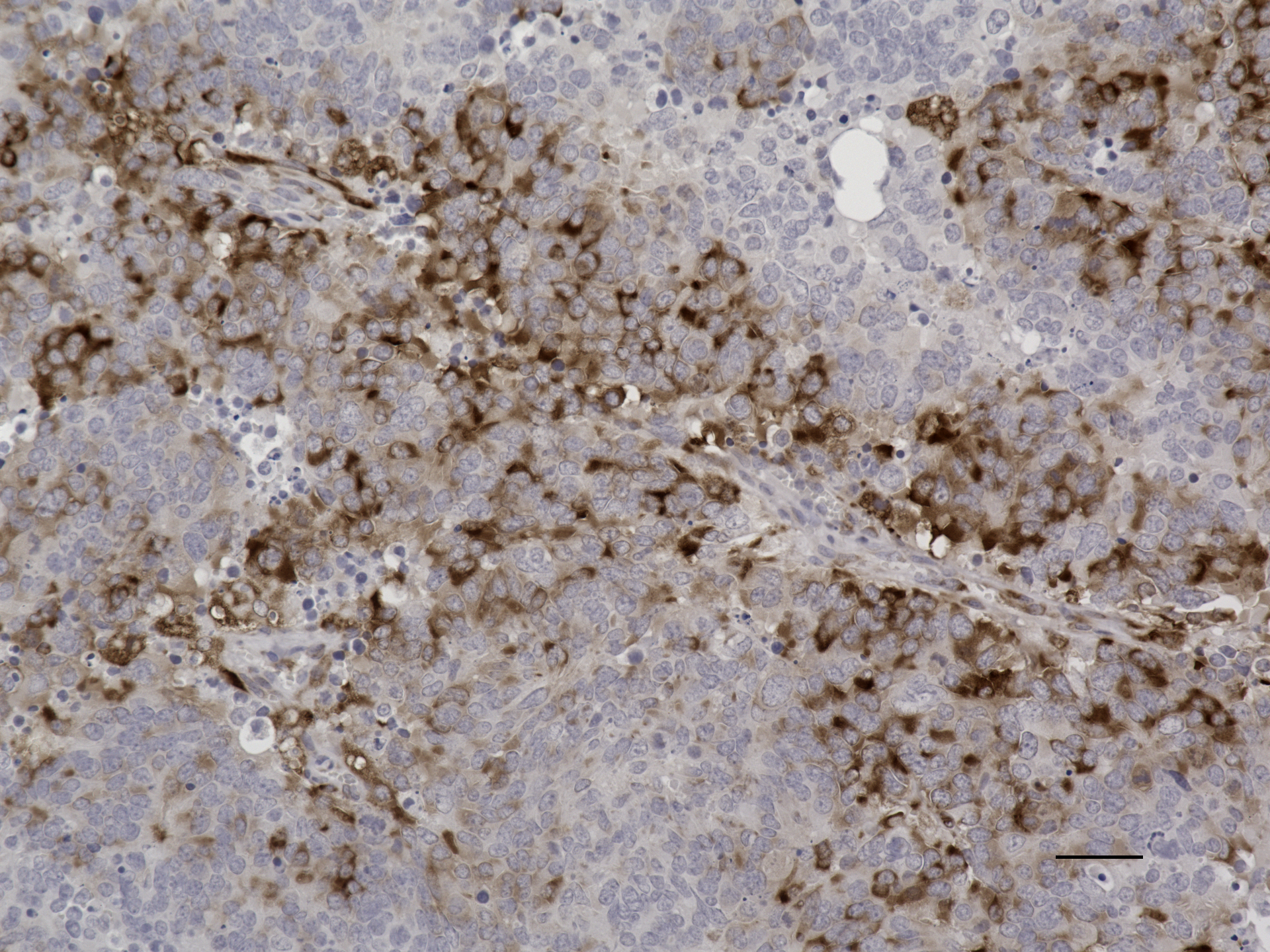

### IPDrosha vehicle pS6 20x.jpeg

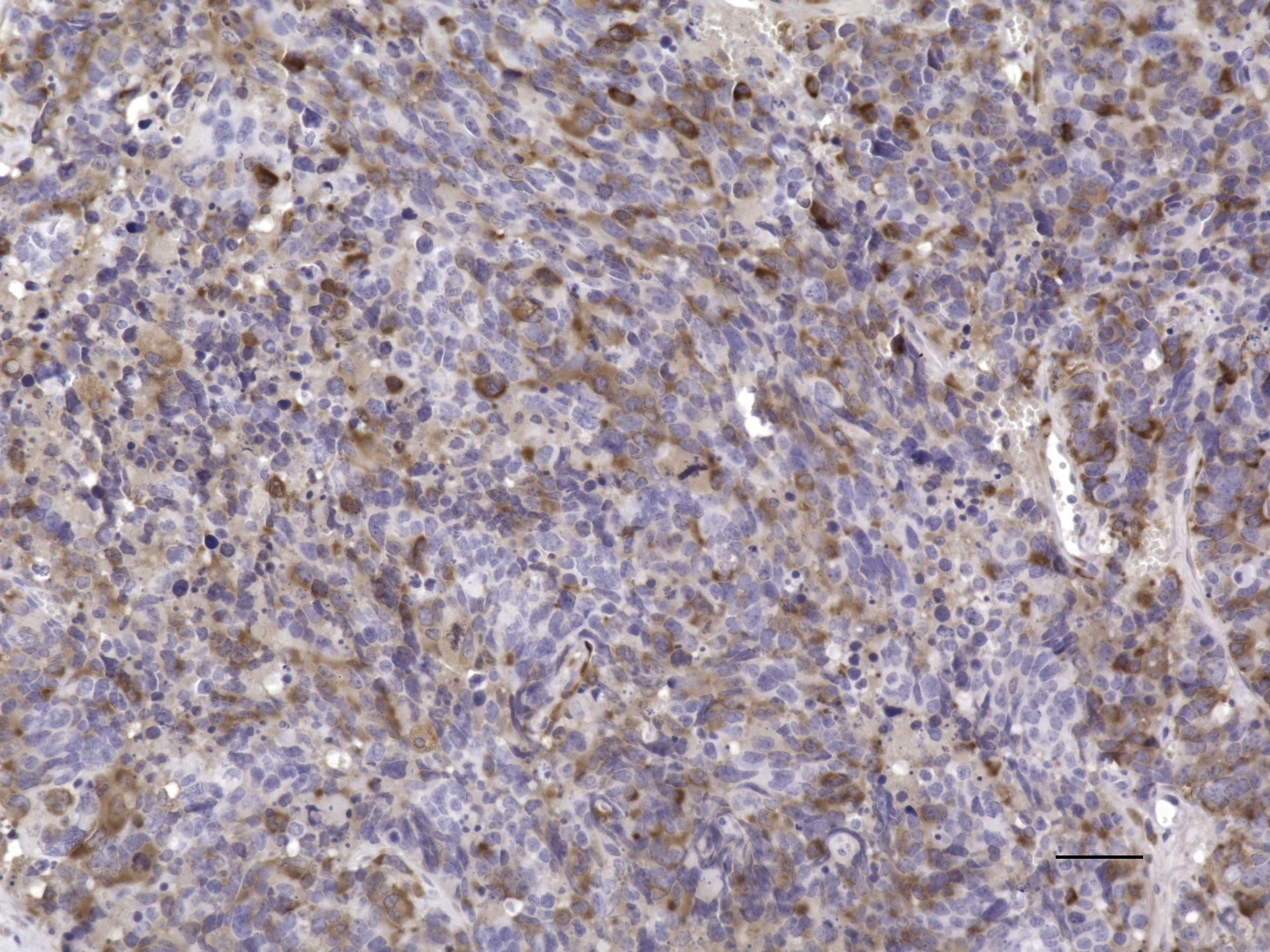

### IPRb1 2X 1000um.jpeg

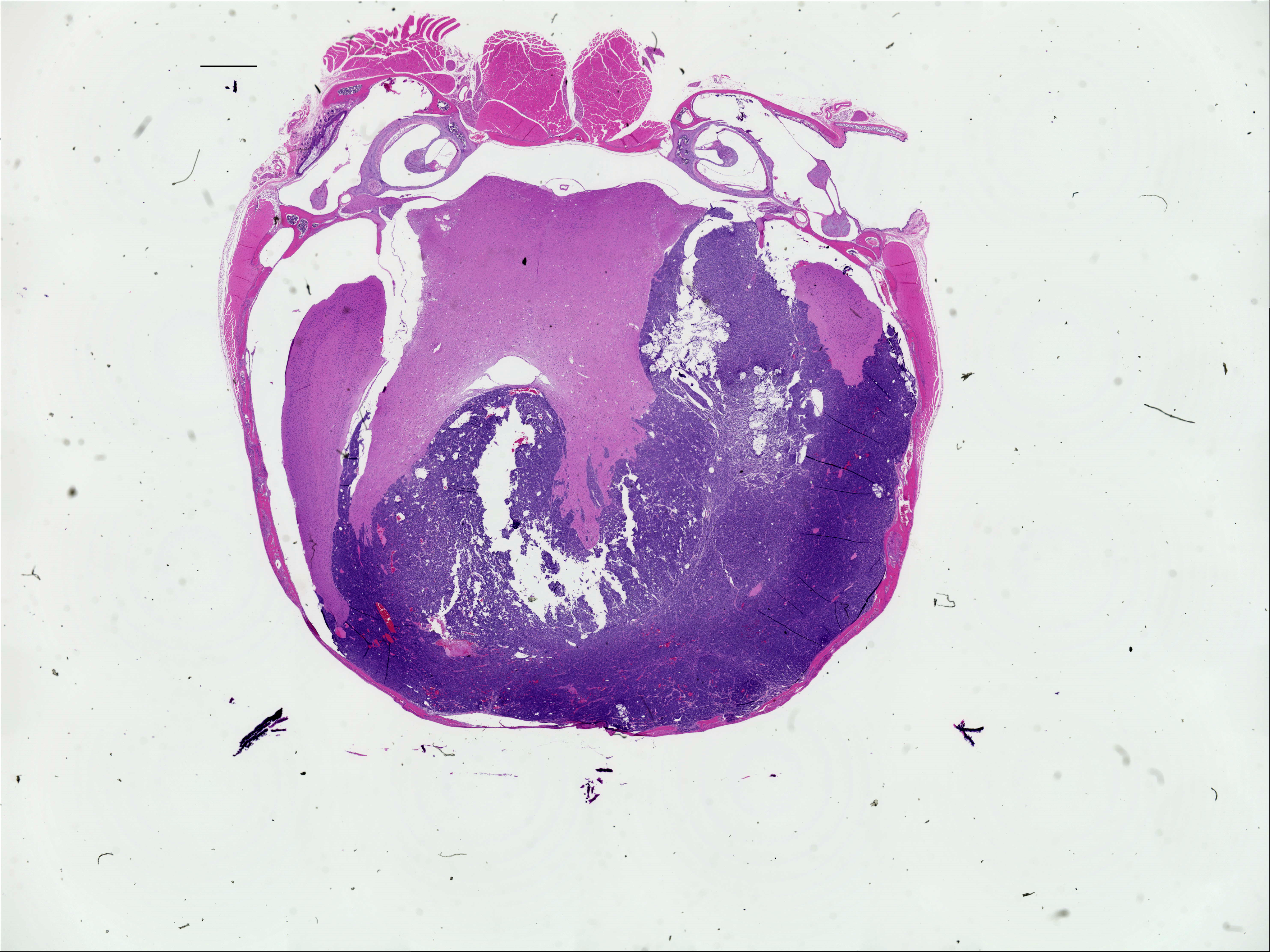

### IPRb1 20X 50um.jpeg

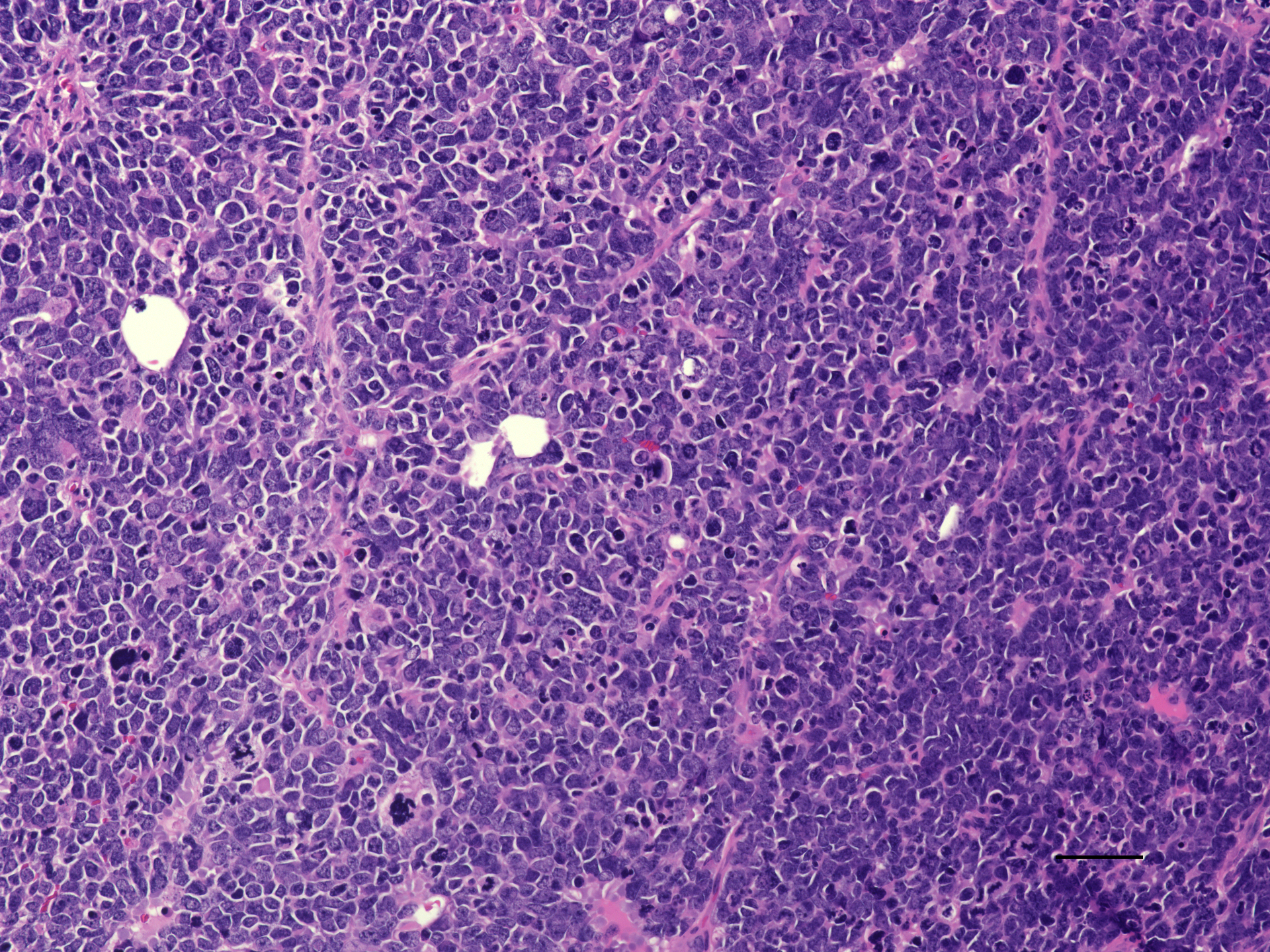

### IPRb1 GFAP 50 um 20X.jpeg

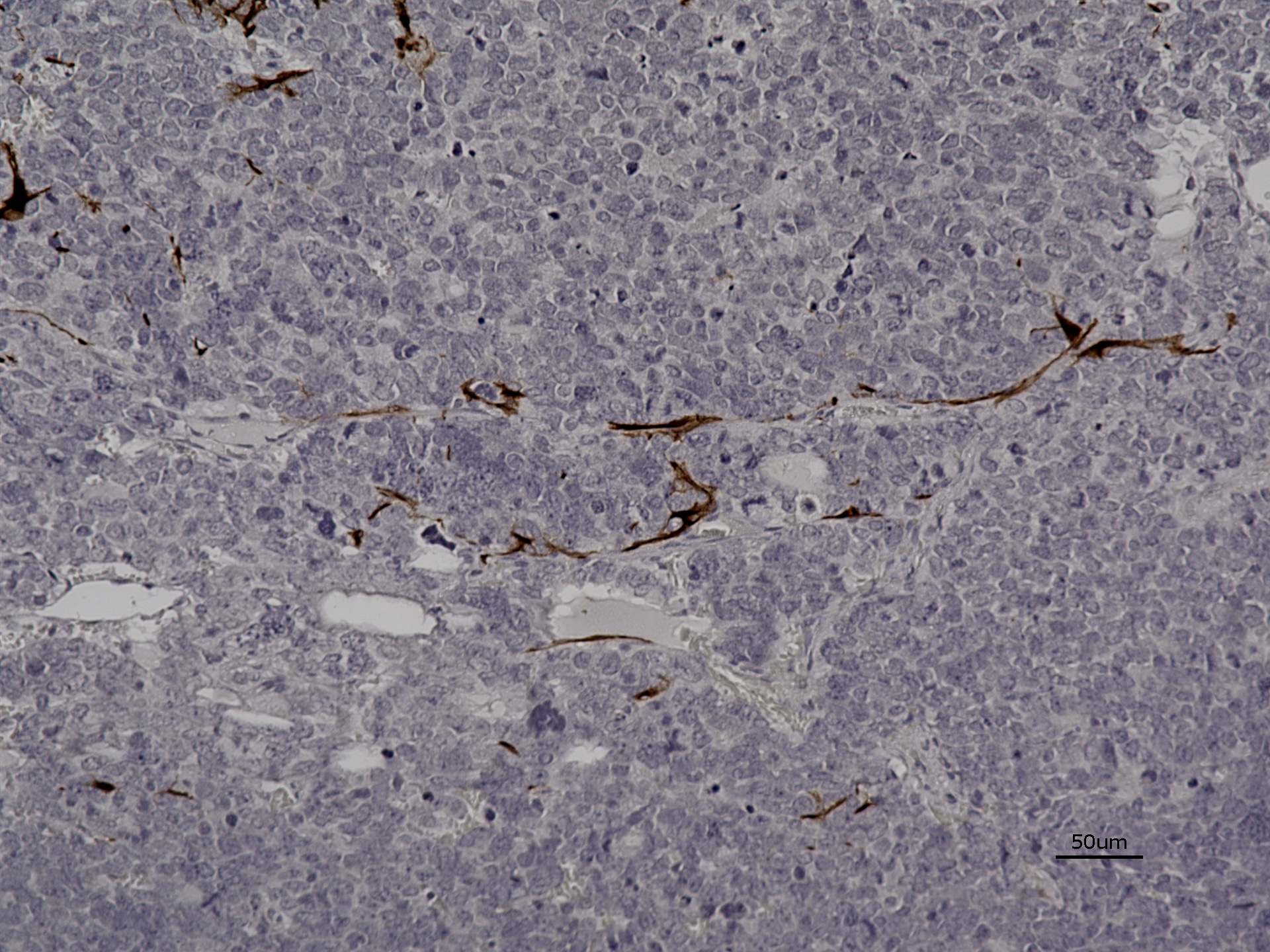

### IPRb1 Ki67 20X.jpeg

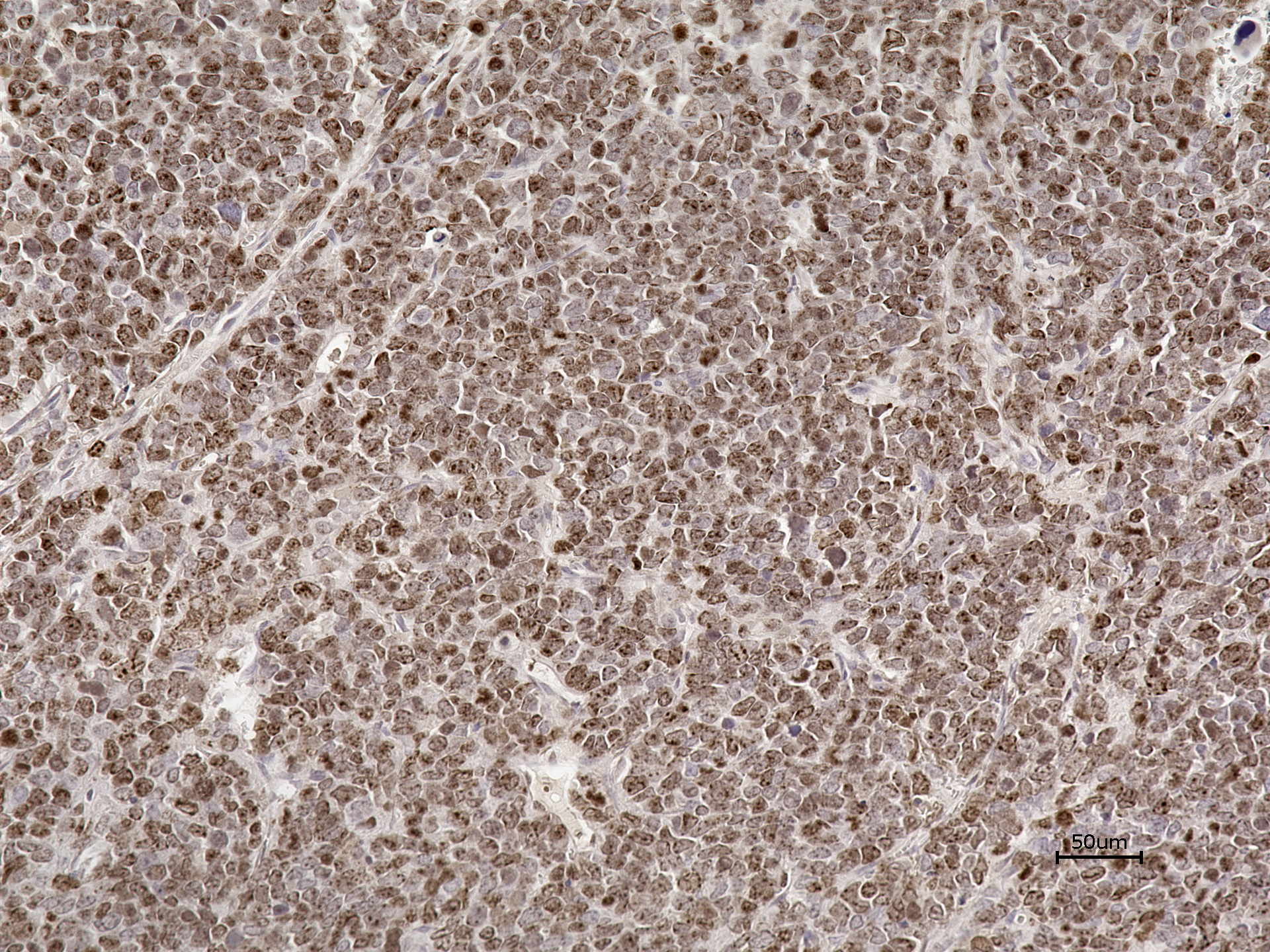

### IPRb1 pRb_20X.jpg

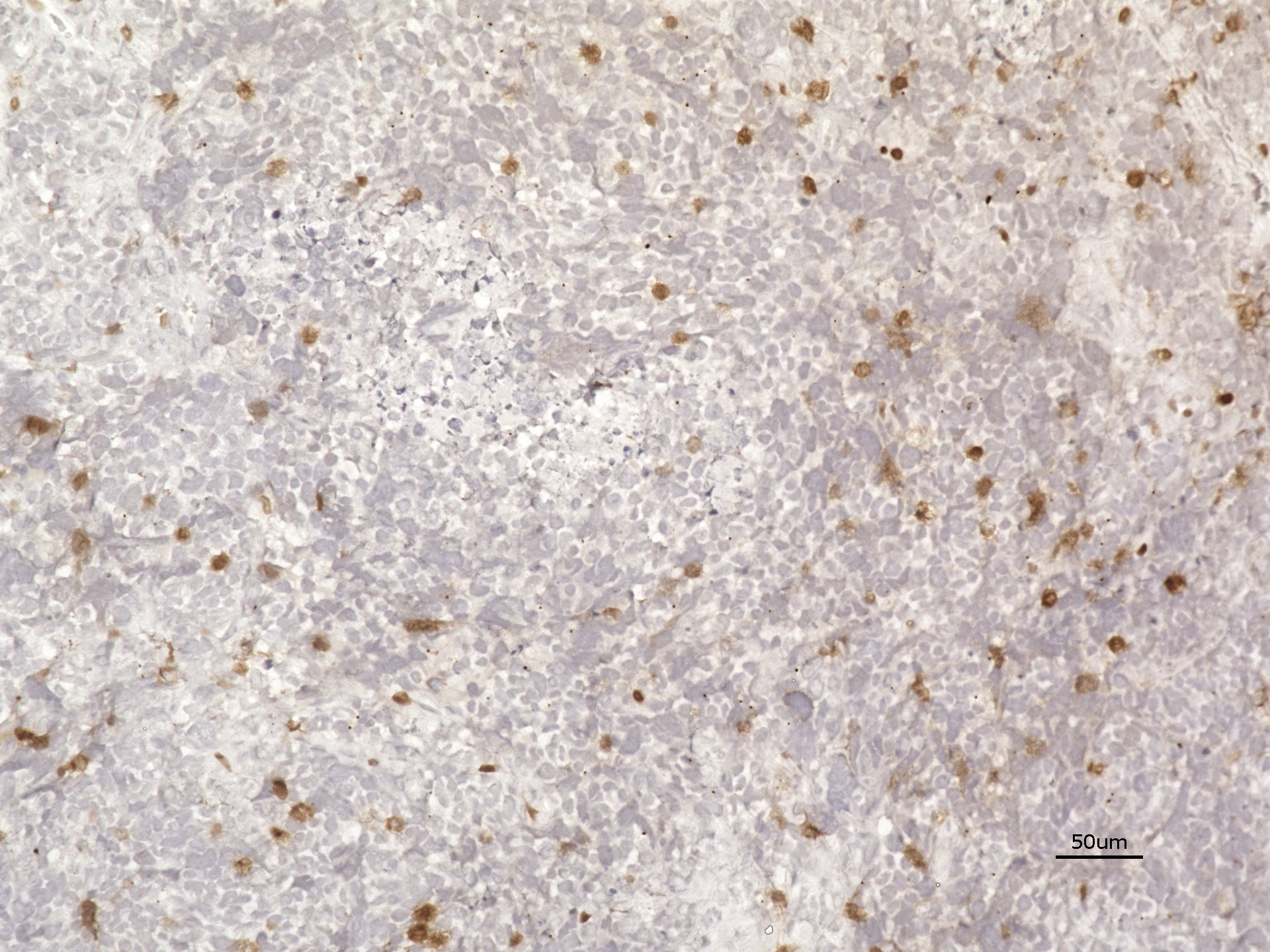

### IPRb1 synaptophysin 20x.jpeg

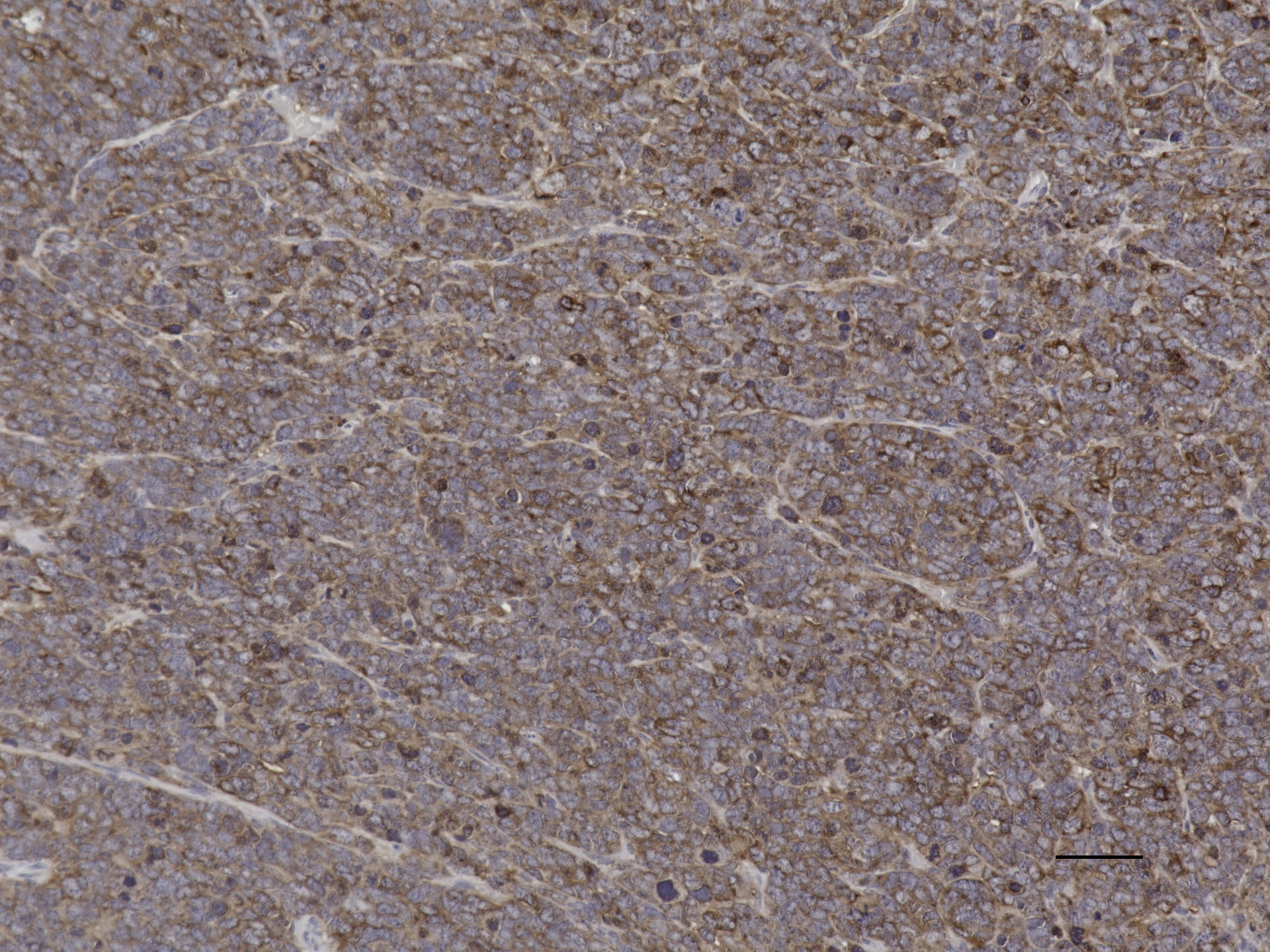

### Source Data 1

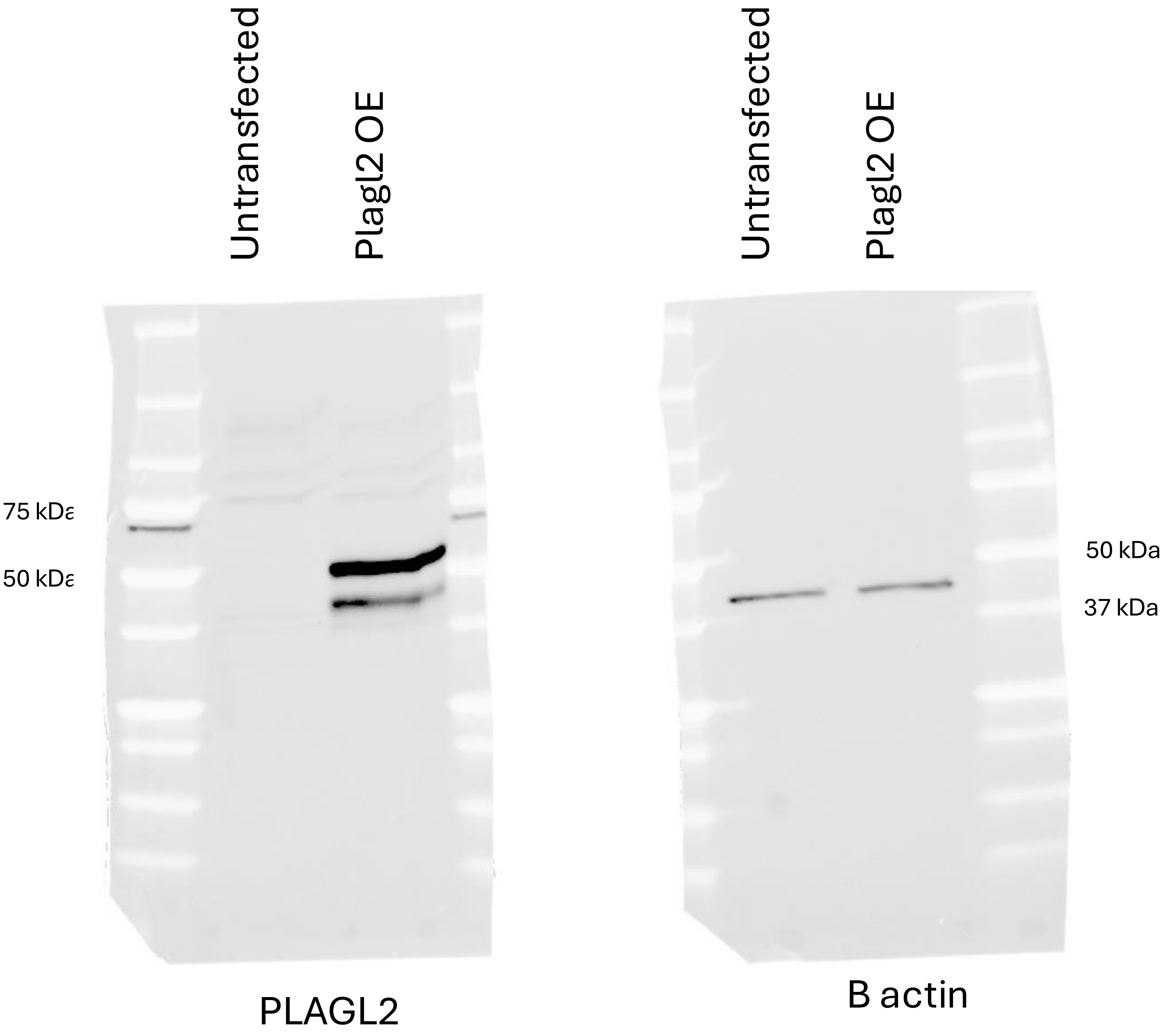


Source data for Suppl. Fig. S7C

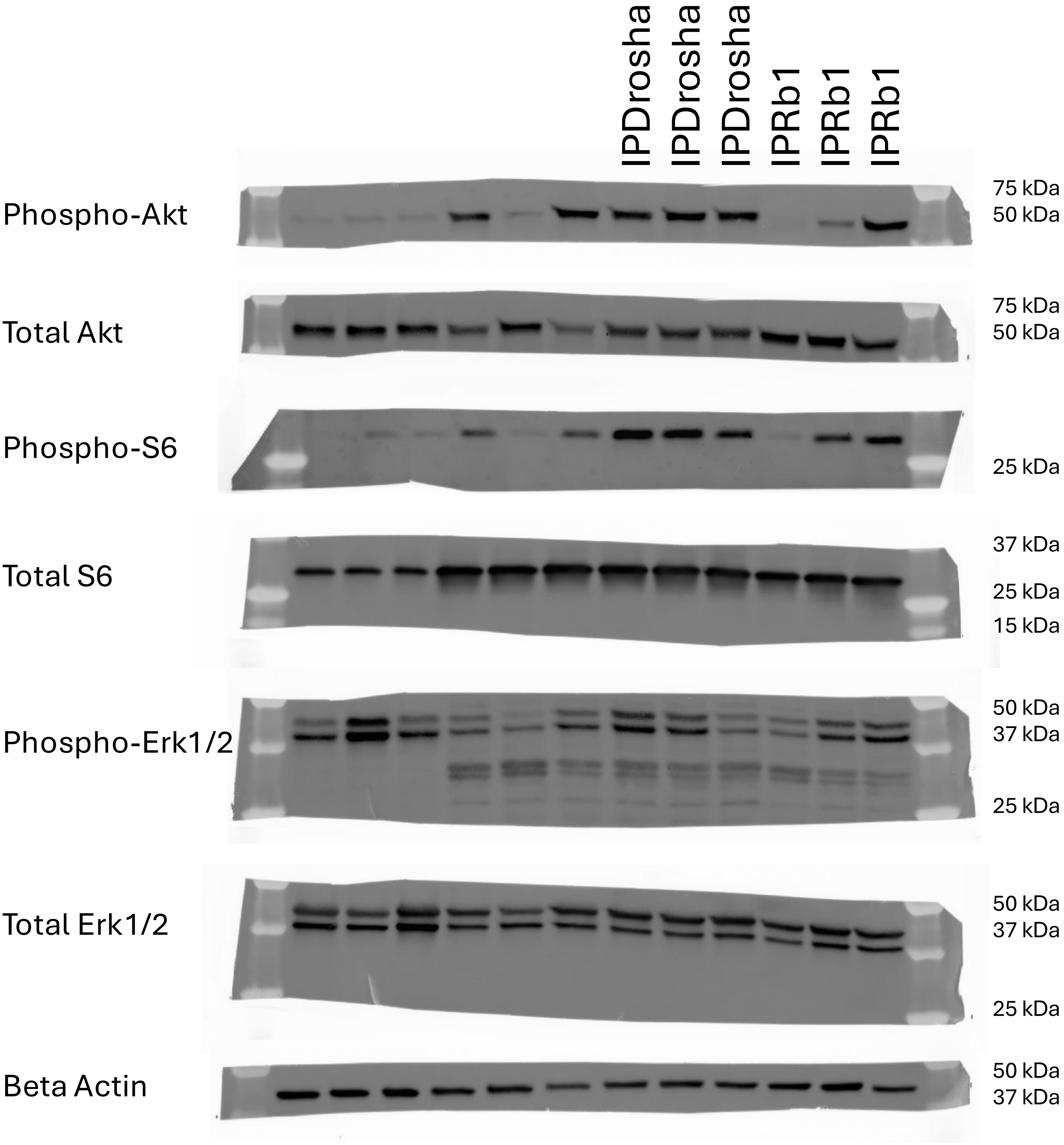


Source data for Suppl. Fig. S8A
