## Supplementary Text for "An imbalance between proliferation and differentiation underlies the development of microRNA-defective pineoblastoma"

**SUPPLEMENTARY FIGURE LEGENDS**

**Supplementary Figure S1. IPDrosha, IPDicer1 and IPRb1 pineal tumors retain embryonic pineal expression pathways.**

**(A-B)** Gene set enrichment analysis (GSEA) for embryonic pineal markers in IPDrosha/IPDicer1 tumor **(A)** or IPRb1 tumor **(B)** vs. age-matched adult pineal gland. **(C-D)** GSEA of enrichment for adult pineal markers in IPDrosha/IPDicer1 tumor **(C)** or IPRb1 tumor **(D)** vs. age-matched adult pineal gland. **(E)** Histone marks H3K27ac and H3K4me3 in IPDrosha and IPRb1 tumors at loci of pineal homeobox transcription factors.

**Supplementary Figure S2. Comparison of IPDrosha, IPDicer1 and IPRb1 pineal tumors to other murine tumor models.**

**(A-C)** Enrichment for proliferation markers (587 genes) **(A)**, proliferation-independent embryonic pineal markers (176 genes) **(B)**, or proliferation-independent adult pineal markers (198 genes) **(C)** across various mouse tumor models from publicly available RNA-seq. Specific comparisons are as follows: IPDrosha/IPDicer1 tumors vs. normal pineal, IPRb1 tumors vs. normal pineal, SHH medulloblastoma vs. normal cerebellum, Group 3 medulloblastoma vs. normal cerebellum, MYCN-driven glioma vs. normal cerebellum or olfactory bulb, orthotopic glioblastoma allografts vs. normal brain, histone H3.3 K27M diffuse midline glioma vs. control neural progenitor cells, and TH-MYCN-derived neuroblastoma vs. normal celiac ganglia.

**Supplementary Figure S3. *Drosha*-driven tumors exhibit loss of canonical microRNAs.**

**(A)** Total normalized reads aligning to microRNAs in IPDrosha tumors and IPRb1 tumors (***$p<0.001$ vs. IPRb1 tumors by two-sided t-test). **(B-D)** Relative expression of Drosha-independent and canonical microRNAs in box-and-whisker plots comparing IPDrosha tumors vs. IPRb1 tumors **(B)**, IPDrosha tumors vs. normal brain **(C)**, or IPDrosha tumors vs. normal adult pineal gland **(D).** (Values shown are $\log_2$ fold change; *$p < 0.05$, ***$p < 0.001$, by two-sided t-tests.) **(E)** Top individual microRNAs expressed in IPDrosha and IPRb1 tumors (*$p<0.05$ vs. IPRb1 tumors, by two-sided t-test). **(F)** Top individual microRNAs expressed in IPDrosha and adult pineal glands (**$p<0.01$ vs. adult pineal by two-sided t-test). **(G)** Expression profile of individual let-7/miR-98-5p family microRNAs in IPDrosha and IPRb1 tumors. **(H)** Expression profile of let-7/miR-98-5p family microRNAs in IPDrosha tumors and adult pineal glands. **(I)** Histone marks H3K27ac and H3K4me3 in IPDrosha and IPRb1 tumors at *Lncppara* (*Mirlet7bhg*) locus.

**Supplementary Figure S4. Patterns of microRNA regulation in miR-eCLIP sequencing of IPRb1 tumors.**

**(A)** Reproduced non-chimeric and chimeric peaks in two IPRb1 tumors. **(B)** Distribution of miR-eCLIP reads across 5' UTR, CDS, and 3' UTR for non-chimeric and chimeric reads in IPRb1 tumors. **(C)** Most significantly enriched motifs in non-chimeric miR-eCLIP reads in IPRb1 tumors. **(D)** Most abundant microRNAs in chimeric reads, with members of the let-7/miR-98-5p family highlighted. **(E)** Cumulative distribution function showing relative expression of microRNA target genes (as identified from miR-eCLIP) in IPDrosha vs. IPRb1. Genes with chimeric peaks were expressed at higher levels than those without chimeric peaks (Kolmogorov-Smirnov test). **(F)** Number of microRNA targets among genes upregulated (n=2,944) or downregulated (n=2,107) in IPDrosha/IPDicer1 vs. IPRb1 tumors (chi-square test).

**Supplementary Figure S5. Regulation of *Ccnd2* and *Mycn*.**

**(A-B)** Enrichment for E2F targets gene set in IPDrosha/IPDicer1 **(A)** or IPRb1 **(B)** vs. adult pineal gland. **(C)** qPCR for *Ccnd1* and *Ccnd3* in normal brain, IPDrosha, IPDicer1, and IPRb1 tumors. Values shown are mean ± SD (*p > 0.05 for all comparisons vs. normal brain, by two-sided t-tests). **(D)** Chimeric miR-eCLIP reads from two IPRb1 tumors in the 3' UTR of *Ccnd2,* with microRNA chimeric read peaks denoted below. **(E)** Expression of *Myc* family members and *Max* by RNA-seq (TPM, transcripts per million). **(F)** Chimeric miR-eCLIP reads from two IPRb1 tumors in the 3' UTR of *Mycn,* with microRNA chimeric read peaks denoted below.

**Supplementary Figure S6. Palbociclib in IPDrosha/IPDicer1 tumors.**

**(A)** Survival analysis of IPDrosha and IPDicer1 tumors treated with vehicle or palbociclib (**p<0.01, ***p<0.001, vs. vehicle, by log-rank test). **(B)** Top "hallmark" gene sets enriched in IPDrosha palbociclib vs. vehicle. **(C)** IHC quantification for phospho-Rb1 (Ser807/811) in the top row and Ki-67 in the bottom row for IPDrosha and IPDicer1 tumors treated with vehicle or palbociclib (***p<0.001, by Chi square analysis) **(D-E)** GSEA of enrichment for adult **(D)** or embryonic **(E)** pineal markers in IPDrosha palbociclib- vs. vehicle- treated tumors.

**Supplementary Figure S7. Regulation of *Plagl2.***

**(A)** Histone marks H3K27ac and H3K4me3 in IPDrosha and IPRb1 tumors at the *Plagl2* locus. **(B)** Expression of *Plagl2*, *Ccnd2*, and *Igf2* in rat pineal gland at different stages of development. **(C)** Expression of ectopically expressed Plagl2 in HEK293 by Western blot. See Source Data file for full uncropped blots.

**Supplementary Figure S8. Ceritinib treatment.**

**(A)** Phosphorylation of mTOR and MAPK signaling intermediates in IPDrosha and IPRb1 tumors by Western blot. Molecular weight markers in kDa denoted on right side of each blot. See Source Data file for full uncropped blots. (**B**) IHC quantification for phospho-S6 (Ser235/236) in the top row and phospho-Rb1 (Ser807/811) in the bottom row for IPDrosha and IPDicer1 tumors treated with vehicle or palbociclib (***p<0.001, by Chi square analysis). **(C)** Tumor volumes for IPRb1 tumors in mice dosed with vehicle (n=9) or ceritinib (n= 8). Ceritinib was given at 50 mg/kg/day by oral gavage, 5 days/week. Values shown are mean ± SD. **(D)** IHC quantification for Ki-67, phospho-S6 (Ser235/236), and phospho-Rb1 (Ser807/811) for IPDrosha tumors treated with vehicle, ceritinib, palbociclib, or both (***p<0.001, by Chi square analysis).

**Supplementary Figure S9. *DROSHA* expression is negatively correlated with let-7/miR-98-5p target genes and E2F target genes in human tumors.**

**(A)** Expression of *DROSHA* and other genes in *DROSHA*-low vs. *DROSHA*-high human pineoblastomas from St. Jude Cloud RNA-seq (center line, median; box limits, upper and lower quartiles; whiskers, 1.5x interquartile range). FPKM, fragments per kilobase per million reads. **(B)** GSEA showing enrichment for target genes of individual microRNA families in total RNA sequencing of *DROSHA*-low vs. *DROSHA*-high human pineoblastoma. **(C)** GSEA showing enrichment for target genes of the let-7/miR-98-5p microRNA family in *DROSHA*-low vs. *DROSHA*-high pineoblastoma tumors. **(D)** Enrichment for E2F targets gene set in *DROSHA*-low vs. *DROSHA*-high pineoblastoma tumors. **(E)** Model for how loss of *Drosha*, *Dicer1*, or *Rb1* leads to proliferation of pineal progenitors.

### SUPPLEMENTARY DATA

**Supplementary Data 1.** Normalized microRNA sequencing counts from IPDrosha tumors and IPRb1 tumors and from IPDrosha tumors, IPDrosha brain, and age-matched pineal glands.

**Supplementary Data 2.** Reproduced chimeric peaks in miR-eCLIP sequencing from IPRb1 tumors.

**Supplementary Data 3**. Gene set enrichment analysis of “hallmark” gene sets in IPDrosha/IPDicer1 tumors vs. adult pineal glands, IPRb1 tumors vs. adult pineal glands, and IPDrosha/IPDicer1 tumors vs. IPRb1 tumors.

**Supplementary Data 4.** Gene set enrichment analysis of “hallmark” gene sets and predicted microRNA target gene sets in *DROSHA*-low vs. *DROSHA*-high tumors in human pineoblastoma RNA-seq.

**Source Data 1.** Uncropped Western blot images (pertaining to **Suppl. Fig. S7C** and **S8A**).

**Source Data 2.** Uncropped microscopy images (pertaining to **Figs. 1**, **3**, **4**, **6**, and **7**).

### SUPPLEMENTARY TABLES

#### Supplementary Table 1. Oligos used in this study.

| Name | Sequence |
|---|---|
| 18s_qF | GTAACCCGTTGAACCCCATT |
| 18s_qR | CCATCCAATCGGTAGTAGCG |
| Otx2_qF | AAATCTCCCTGAGAGCGGAAC |
| Otx2_qR | TCCAAATAGCCAGCTATCAAAGT |
| Crx_qF | TTCCTCTAGCCTCTGCTGTCT |
| Crx_qR | CAGATGAGGGCACCTTTGGAA |
| Rax_qF | GTTCGGGTCCAGGTATGGTT |
| Rax_qR | CTGCAGCTTCATGGACGACA |
| Pax6_qF | GTCAGATCTGCTACTTCCCCC |
| Pax6_qR | TGGTTAAAGTCTTCTGCCTGTGA |
| mIgf2_qF1 | CACATTCGGCCTCTGCGAC |
| mIgf2_qR1 | ATCCCCATTGGTACCTGGAAG |
| Ccnd2_qF | CTGTGCGCTACCGACTTCAA |
| Ccnd2_qR | ATCATCCTGCTGAAGCCCAC |
| Ccnd3_qF | CTTTGTTTGGGTGCCCAGGAA |
| Ccnd3_qR | AGAGCATTTCAGGGCGAGCTT |
| Ccnd1_qF | GCAGAAGGAGATTGTGCCATCC |
| Ccnd1_qR | AGGAAGCGGTCCAGGTAGTTCA |
| Plagl2-qF | CCAAGTACAAGCTGTATAGGCACAT |
| Plagl2-qR | CAGTGGAGGGCCTCTTTGT |
| Chrna3-qF | CCTGTTCCAGTACCTGTTTGAAG |
| Chrna3-qR | TGATCTGGTTTACTTCATCCACC |
| Nefh-qF | CTCGCGACGCCCTCA |
| Nefh-qR | GTCGGTCCAACCTCACTCG |
| neg_ChIP_qF | ATTTTGTGCTGCATAACCTCCT |
| neg_ChIP_qR | TAGCAACATCCTAAGCTGGACA |
| Ccnd2-ChIP F | TTGGTAGAGGGGGACACGTTG |
| Ccnd2-ChIP R | AAGCGTCCCTGATCATCCCC |
| Igf2_ChIP F | CTTTTCGCTGCAGTCCCGAG |
| Igf2_ChIP R | AACTTCGAAGGACCGAGGAC |
| Ccnd2 promoter region 1 F | NNNNCTCGAGCCTAAATGAGGAACAAGGAAAGGC |
| Ccnd2 promoter region 1 R | NNNNGGTACCTTTTGATGTAGCGGAGAAGGCT |
| Ccnd2 promoter region 2 F | NNNNCTCGAGAAAGCATCCTGTGGTCAAGGC |
| Ccnd2 promoter region 2 R | NNNNGGTACCAATCTGCCCGGTAACAATCAGC |
| Igf2 promoter sequence | CTCGAGTTGCGGGGGCGGTCCCGGGGCGGGGCGAGGGCC<br>CTGCGGACGCCCATTGGCGCGGGCGTAAGGCCAGCGGGGC<br>CCGAGCGGGCGCCGAGCCGCGGGGTGGCGCGGCTATAAGA<br>ACCGGGCGTTGGCGCCCGGAGTTCGCCTGCTCTCCGGCGG<br>AGCTGCGTGAGGCCAGGCCGGCCCCCGGCCCCCCCTTCCG<br>GCCGCCCCCGCCTCCTGGCCCACGCCCGCCCGCGCTCGGC<br>CCGCCAGCGCCTCCATCCGGGCTGGCGGCCCCGCGTCGAC<br>GCCGTCCGCCACCTCGCTGCTGCTAACTCCTGTGCAGGGCG<br>CCGTCGGCGGGGCCGCGCTCCGTCGGGCCTGCGGATCTCC |

| | |
|---|---|
| | CCACCGCCTCCTCCTCTATCTACCTCAACACCCCATTCCTGCT<br>TCGCCAGAGGAGGCGGTCCCCACCGCAGGCAGTCCGGCTT<br>GCAGGTCGCCGGCGTTGTCATCCCCCGCGCTTCCCCTCCCA<br>GCCCTCCCCGGTGCGCAGCCCGGCTGCTCCCCTCTTTTCGC<br>TGCAGTCCCGAGCAGCTGAGGCGCCGCCACGCCTGTCCCC<br>CCCACAAGAAGCCCGGGCTTACGACGGCTGAGGGCTCCGTC<br>GACCCTAACCGAGCTGGGTGCCCGTGGCCGGGGTGACGCC<br>TCCATTCCTCCCCCCCTCAACACCGTCCTCGGTCCTTCGAAGT<br>TGCATCCTCTCCTCTGCTTAGGGTGCGCCCCCCCTCGCGCA<br>CCCGCTTACCCGCCACCTTTCCTAAGCTCCCCTCCTGCCCCC<br>TCCCCGTTCCTCCTCGCCTCAGACTCCCTCCCCCTCACGTCC<br>GCCCTCTGCCTTCGCCTACCCAAGTGGATTAATTATACGCTTT<br>CTGTTTCTCTCCGTGCTGTCCTCTCCCGCTGTGAGCCTACCC<br>GCCTCTCGCTGTAAGCTT |

**Supplementary Table 2. Antibodies used in this study.**

| Target | Manufacturer and catalog number | RRID |
|---|---|---|
| Ki-67 (1:500) | Abcam, ab15580 | AB_443209 |
| Phospho-Rb (Ser807/811, 1:400) | Cell Signaling Technology, 8516 | AB_11178658 |
| Synaptophysin (1:200) | Cell Signaling Technology, 36406 | AB_2799098 |
| Phospho-S6 (Ser235/236, 1:200) | Cell Signaling Technology, 2211 | AB_331679 |
| GFAP (1:200) | Cell Signaling Technology, 80788 | AB_2799963 |
| PLAGL2 (ChIP) | Sigma Aldrich, SAB3500815 | AB_1079629 |
| Rabbit IgG | Cell Signaling Technology, 2729 | AB_1031062 |
| H3K4me3 | Epicypher, 13-0041 | AB_3076423 |
| H3K27ac | Cell Signaling Technology, 8173 | AB_10949503 |
| Rabbit IgG | Epicypher, 13-0042 | AB_2923178 |
| PLAGL2 (Western blot, 1:1000) | Proteintech, 11540-1-AP | AB_2165035 |
| Phospho-Akt (1:1000) | Cell Signaling Technology, 4060 | AB_2315049 |
| Akt (1:2000) | Cell Signaling Technology, 9272 | AB_329827 |
| Phospho-MAPK (1:1000) | Cell Signaling Technology, 9101 | AB_331646 |
| MAPK (1:1000) | Cell Signaling Technology, 9102 | AB_330744 |
| S6 (1:1000) | Cell Signaling Technology, 2217 | AB_331355 |
| Beta actin (1:1000) | Cell Signaling Technology, 4970 | AB_2223172 |
| Anti-Rabbit IgG, HRP-linked | Cell Signaling Technology, 7074 | AB_2099233 |
